# Human pathogenic RNA viruses establish non-competing lineages by occupying independent niches

**DOI:** 10.1101/2021.12.10.472150

**Authors:** Pascal Mutz, Nash D. Rochman, Yuri I. Wolf, Guilhem Faure, Feng Zhang, Eugene V. Koonin

**Affiliations:** National Center for Biotechnology Information, National Library of Medicine, Bethesda, MD 20894; Broad Institute of MIT and Harvard, Cambridge, MA 02142; Howard Hughes Medical Institute, Massachusetts Institute of Technology, Cambridge, MA 02139; McGovern Institute for Brain Research, Massachusetts Institute of Technology, Cambridge, MA 02139; Department of Brain and Cognitive Sciences, Massachusetts Institute of Technology, Cambridge, MA 02139; Department of Biological Engineering, Massachusetts Institute of Technology, Cambridge, MA 02139

**Author notes:** Authors contributed equally (Pascal Mutz, Nash D. Rochman).

**Keywords:** effective population size, human RNA viruses, pandemic, globalization, Influenza, SARS-CoV-2, viral phylodynamics

## Abstract

Many pathogenic viruses are endemic among human populations and can cause a broad variety of diseases, some potentially leading to devastating pandemics. How virus populations maintain diversity and what selective pressures drive population turnover, is not thoroughly understood. We conducted a large-scale phylodynamic analysis of 27 human pathogenic RNA viruses spanning diverse life history traits in search of unifying trends that shape virus evolution. For most virus species, we identify multiple, co-circulating lineages with low turnover rates. These lineages appear to be largely noncompeting and likely occupy semi-independent epidemiological niches that are not regionally or seasonally defined. Typically, intra-lineage mutational signatures are similar to inter-lineage signatures. The principal exception are members of the family *Picornaviridae*, for which mutations in capsid protein genes are primarily lineage-defining. The persistence of virus lineages appears to stem from limited outbreaks within small communities so that only a minor fraction of the global susceptible population is infected at any time. As disparate communities become increasingly connected through globalization, interaction and competition between lineages might increase as well, which could result in changing selective pressures and increased diversification and/or pathogenicity. Thus, in addition to zoonotic events, ongoing surveillance of familiar, endemic viruses appears to merit global attention with respect to the prevention or mitigation of future pandemics.

**Significance:** Numerous pathogenic viruses are endemic in humans and cause a broad variety of diseases, but what is their potential of causing new pandemics? We show that most human pathogenic RNA viruses form multiple, co-circulating lineages with low turnover rates. These lineages appear to be largely noncompeting and occupy distinct epidemiological niches that are not regionally or seasonally defined, and their persistence appears to stem from limited outbreaks in small communities so that a minor fraction of the global susceptible population is infected at any time. However, due to globalization, interaction and competition between lineages might increase, potentially leading to increased diversification and pathogenicity. Thus, endemic viruses appear to merit global attention with respect to the prevention of future pandemics.

## Introduction

Viruses, ubiquitous across the tree of life, occupy an astounding diversity of ecological niches (1-3). Viral niches are primarily defined by the behavior and immunity of the respective hosts and are often the subject of deep, but narrow investigation (4, 5). In this work, we sought to uncover common trends at relatively short evolutionary distances by studying the microevolution of human pathogenic RNA viruses. The devastating COVID-19 pandemic has made it abundantly clear that understanding these microevolutionary features is of vital importance not only to forward our understanding of virology in general but to inform appropriate public health measures during a pandemic (6, 7).

Viral populations explore their viable sequence space defined both by intrinsic constraints and those imposed by host behavior (8) through the accumulation of mutations, potentially leading to diversification (9). A single host species can offer multiple independent niches that are explored by distinct virus subpopulations. Niches can be formed and maintained through regional or seasonal separation. Regional separation of subpopulations has been demonstrated, for example, for yellow fever virus (YFV) (10, 11). At sufficiently long evolutionary distances, niches can be defined by immunological differences, which enable a viral subpopulation to overcome immune cross-protection, allowing the same host to be infected by multiple subpopulations largely independent of prior infections. This phenomenon has been demonstrated for enteroviruses with many co-circulating serotypes (12). These immunological niches do not need to be spatially or temporally segregated.

Generally, niches are not necessarily static entities and can overlap or merge depending on dissemination rates, transmission modes, and other life history traits (2). When outbreaks are limited to small communities so that only a small fraction of the global susceptible population is infected at any time, niches can form that are not regionally or seasonally defined but are still maintained through a combination of spatial and temporal separation at a local scale. Thus, the maintenance of these viral niches is highly sensitive to changes in host behavior. The number and sequence diversity of such lineages depends on constraints intrinsic to viral biology as well as host behavior(13, 14). For example, there is a sharp contrast between the emergence of immunological niches among measles morbilli virus (MMV) and influenza A virus (IAV) strain H3N2 (named H3N2 here). Through rapid antigenic drift and shift that involve a non-human host reservoir, IAV is able to overcome adaptive-immune protection, despite infecting a substantial fraction of the susceptible host population each year (15). As a consequence, H3N2 goes through phases of stasis, in which neutral evolution and purifying selection are dominant; parallel lineages are established; and population diversity grows. Once the pool of naïve hosts shrinks, the competition between lineages intensifies, resulting in a short phase of strong positive selection that favors one lineage to replace all others (16-18). In contrast, no such antigenic drift has been observed for MMV, and parallel MMV lineages do not replace each other, but rather stably co-exist (17, 19).

The persistence of multiple, co-existing viral lineages implies minimal inter-lineage competition. When such lineages are maintained through spatial or temporal separation, increased host-host or host-vector contact can result in the merger between and competition among multiple lineages. Climate change can support the spread and mixing of previously separated vectors, which could carry distinct viral lineages. With more vectors, the dissemination rate can rise, decreasing the number of susceptible hosts, and increasing competition globally (2). This can result in accelerated lineage turnover of human and agricultural pathogens, with the potential for substantial epidemiological and economic impact (20).

We sought to identify unifying trends of lineage emergence, persistence, and turnover among human pathogenic RNA viruses and to characterize the niches occupied by these lineages through phylodynamic analysis (21). Taking advantage of the substantial recent progress in virus genome sequencing (22), we constructed phylogenetic trees for the genomes of all monopartite human pathogenic RNA viruses for which extensive genome sequence information was available. These phylogenies were employed to assess the selection pressures affecting the evolution of these viruses through an analysis of the ratio of non-synonymous to synonymous substitution rates (*dN/dS*), and to estimate the effective population size (*Ne*) and the census population size (*N*) for each. The viruses studied here are of clear epidemiological relevance, span a broad variety of life history traits (2, 23), and thus seem suitable to reveal unifying trends in the microevolution of RNA viruses. Our analysis of these viruses indicates that most form multiple, coexisting, non-competing lineages which appear to occupy independent niches.

## Results

### Data aggregation

Despite the substantial progress of the past several years (22), the available numbers of (nearly) complete genome sequences of human pathogenic RNA viruses differ widely among viral species. In the data set for the present analysis, we included only those species for which 200 or more (nearly) complete genome sequences, with at least 50 isolated from a human host, were available in the NCBI virus database (24) or GISAID for Severe acute respiratory syndrome-related coronavirus 2 (SARS-CoV-2) (25) (Fig. 1). These criteria excluded viruses which are widespread, for example lyssa rhabdovirus and rubella virus, but for which few (nearly) complete genomes were available as well as comparatively rare, even if highly pathogenic, viruses including some Ebola virus species (Zaire, but not Reston or Sudan, was included) and Marburg virus.

**Figure 1.**
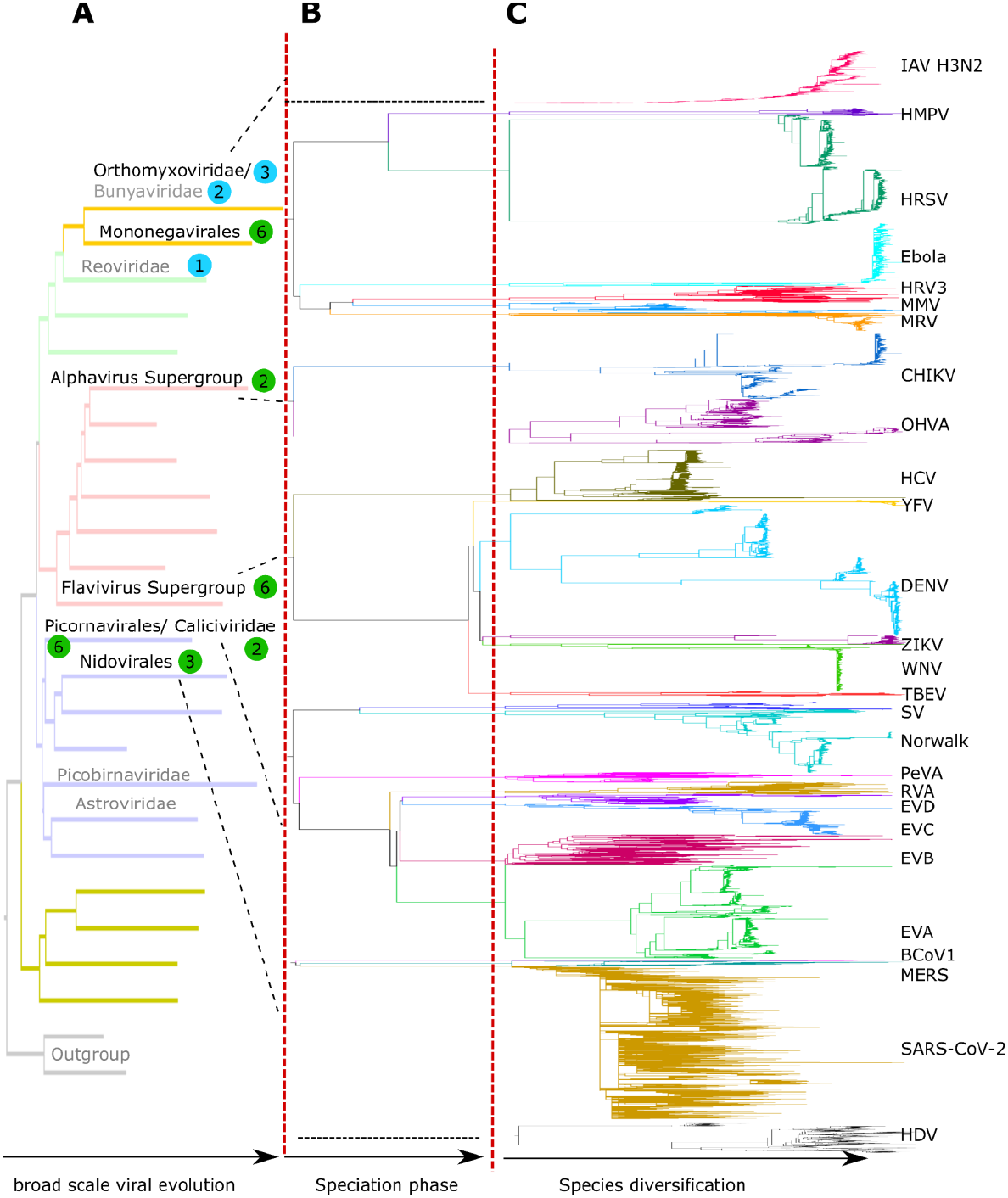
Phylogenies of human pathogenic RNA viruses. Schematic depicting the origins and phylogenetic tree topologies of 27 human pathogenic RNA viruses. **A**. Placement of each virus in the global phylogeny of RNA viruses (realm Riboviria). The tree topology is from (72). Viral groups containing human pathogenic viruses are named in black if containing viruses analyzed in this work and gray otherwise. The numbers of viral species, for which at least 200 nearly complete genome sequences were available, at least 50 of these isolated from humans, are shown in colored circles (green: monopartite viruses; blue: segmented viruses). **B**. Speciation of viral families or orders. **C**. Diversification within species. Trees for species are scaled to the same distance from the root to the most distal leaf and are grafted on the tree scaffold with arbitrary branch lengths for speciation but respecting topology.

Only monopartite RNA viruses were considered in order to exclude potential effects of segment reassortment and enable the construction of a single, unambiguous phylogeny. This restriction excluded 6 species with many genomes available: three Influenza viruses (A, B, and C), Reovirus, Lassa mammarenavirus, and Dabie bandavirus. We further omitted HIV given its retro-transcribing replication strategy. Altogether, our dataset included 26 monopartite virus species. We added IAV H3N2 to this group (with a phylogeny constructed from hemagglutinin) as a thoroughly studied reference virus (14, 16, 17). The 27 viruses analyzed here cover a broad variety of viral lifestyles and ecological constraints and have been subject to varied countermeasures including vaccination (Supplementary Data, Table S1). This diversity enables the exploration of potential unifying trends of viral lineage turnover and niche formation.

Sequences were aligned using MAFFT(26), and with the exception of SARS-CoV-2 and H3N2, for which the large number of sequences necessitated an iterative strategy, phylogenetic trees were constructed using IQ-TREE (27) (see Brief Methods and Extended Methods in the SI Appendix for details). For most of viruses, the resulting trees included several large, clearly distinguishable clades (Fig. 1) that in some cases corresponded to known sero-or genotypes (for example, Dengue virus, DENV, serotypes 1-4).

### Low rates of lineage turnover among human RNA viruses

The major virus clades and the smaller lineages contained within them are subject to turnover whereby an older lineage goes extinct, being gradually replaced by individuals from a newer lineage. Trees with high turnover rates are often described as “cactus”-or “ladder”-like, and in the limit of extreme turnover, as “caterpillar” trees, whereas those with low turnover are often described as “bush”-like, with ultrametric trees representing the limit of no turnover (14, 17). In an effort to explicitly measure lineage turnover (without relying solely on global measures such as coalescence rate (17), which is also estimated), we first sought to establish how many isolates, and distributed on the tree in what way, constitute a lineage. This information is important, in large part because varying substitution rates across the tree complicate the estimation of global lineage turnover (9, 28). We defined lineages as monophyletic groups of sequences separated by periods (branches of the tree) with apparently different substitution rates and within which the sequencing date and the distance to the tree root are significantly correlated (see Brief Methods and Extended Methods in the SI Appendix for details). Lineages cannot be defined in this way to encompass all sequences and Fig. S1 shows the fraction of sequences included in correlated lineages for each virus. Arguably, significantly different substitution rates mark different selective environments and may reflect movement into distinct epidemiological niches. Because there are no apparent periods of different substitution rates within each lineage and, consequently, high-confidence date-constrained genealogical trees with a single substitution rate could be fit for each (see below), we denote these “genealogical lineages” (GL).

Multiple GL were identified in this manner for most viruses (Figs. S2-7), as illustrated in Fig. 2A,B for the three lineages of Enterovirus D (EVD). For human Betacoronavirus 1 (BCoV1), Ebola Zaire (Ebola), MERS (Middle East respiratory syndrome-related coronavirus), H3N2, SARS-CoV-2, and Zika virus (ZIKV), the majority of the phylogeny comprised a single GL. Thus, the entire population of each of these viruses might occupy a single epidemiological niche at any point in time (which may be subject to rapid lineage turnover as is the case for H3N2). For Mumps rubulavirus (MRV) and YFV, the GL identified were not large enough for subsequent analysis. When interpreting our observation of a single GL for Ebola, it should be noted that more than half of the Ebola isolates stem from the 2014-2016 outbreak in Sierra Leone, Liberia, and Guinea (29), the common assumption being that each individual Ebola outbreak stems from an individual zoonotic spillover event (30).

**Figure 2.**
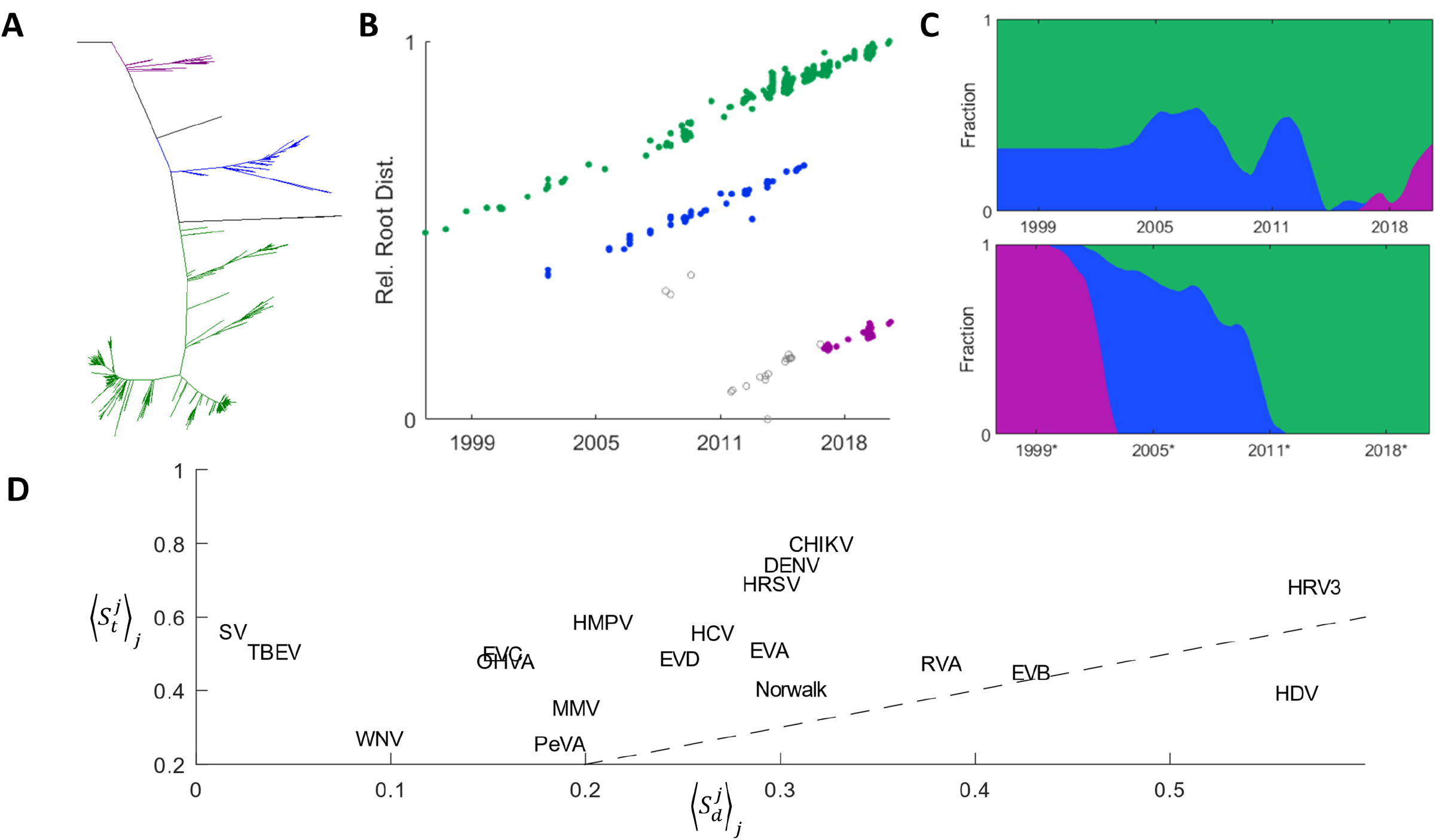
Stable Coexistence of Lineages among Human Pathogenic RNA Viruses. **A**. EVD tree colored to represent the location of the three genealogical lineages (GL). **B**. Distance to the tree root vs. date of isolation for EVD. Distance is scaled by the maximum for any sequence within a correlated-clade and the x axis is bounded by the minimum/maximum date of isolation for any such sequence. **C**. The fraction of isolates within each correlated-clade (and excluding isolates that did not belong to any correlated-clade) computed over a sliding window containing the nearest 5% of all isolates indexed by sequencing date (top) and root distance (bottom, where date* represents the date of isolation corresponding to each sequence in an alternative phylogeny where the date of isolation for each sequence exactly corresponds to the distance to the root for that sequence. **D**. The mean Shannon entropy for the correlated-clade distribution respecting the sequencing date (y-axis, 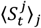) and root distance (x-axis, 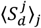) respectively. Dashed line displays 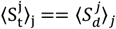.

Having identified the virus GLs, we quantified lineage turnover using the Shannon entropy of the GL distribution over time, *S*_*t*_, as well as travelling up the tree from root to leaf, *S*_*d*_. For this analysis, only sequences included within a GL were considered. First, sliding windows (indexed over *j*) containing the closest 5% of all isolates to the specified date,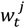, (from the date of the earliest isolated sequence to the latest) or distance to the tree root, 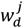, were established and the GL distribution within each window was obtained (Fig. 2C). Next, the probability that a sequence, *x*, within each window belongs to the *i*^*th*^ GL was computed: 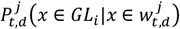.The Shannon entropy of the GL distribution was then calculated using log base *N* equal to the number of GLs identified within the tree (and yielding a maximum value of 1):

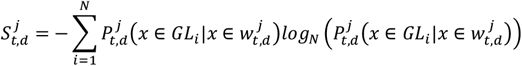

Finally, the mean over all windows was computed for each tree: 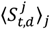 (Fig. 2D). A mean entropy near 0 corresponds to a phylogeny composed of clades that rapidly displace one another (although effects of sampling bias cannot be excluded). A mean entropy near 1 corresponds to a phylogeny, in which all clades are uniformly distributed at every time point. We observed 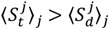 in all but one case (HDV) indicating that entropy is greater than that expected from analysis of the tree structure with no known dates of isolation. This observation coupled with the generally high mean entropy suggests that most of the analyzed viruses evolve with low rates of lineage turnover.

To further quantify lineage turnover, we constructed date-constrained genealogical trees. As suggested above, GLs are separated by periods (branches of the tree) with apparently different substitution rates. These branches are often deep within the tree and are sparsely populated with leaves (if at all), making the assignment of a global model for substitution rates statistically dubious and highlighting the importance of rates inferred for individual GLs. Date constrained trees were produced using a least-square distance approach based on the date of isolation for each sequence (31). A mutation rate and the date of the last common ancestor (LCA) were estimated for all global trees and each GL individually (Fig. 3A). Samples without a known date of isolation introduce additional uncertainty into the calculation and can result in future-dated portions of the genealogical tree (see Brief Methods and Extended Methods in the SI Appendix for details). GLs tend to accumulate over time, with few if any extinction events (Fig. 3B). Note that the apparent decline of parallel GLs for many viruses in recent years (approximately 2016-present) is most likely a sampling artefact although, in principle, such decline could also point to a change in GL dynamics. These trends are further indicative of low lineage turnover and suggestive of minimal competition among GLs. Thus, each GL is likely to occupy a distinct niche, and we sought to identify the factors that shape and maintain these niches.

**Figure 3.**
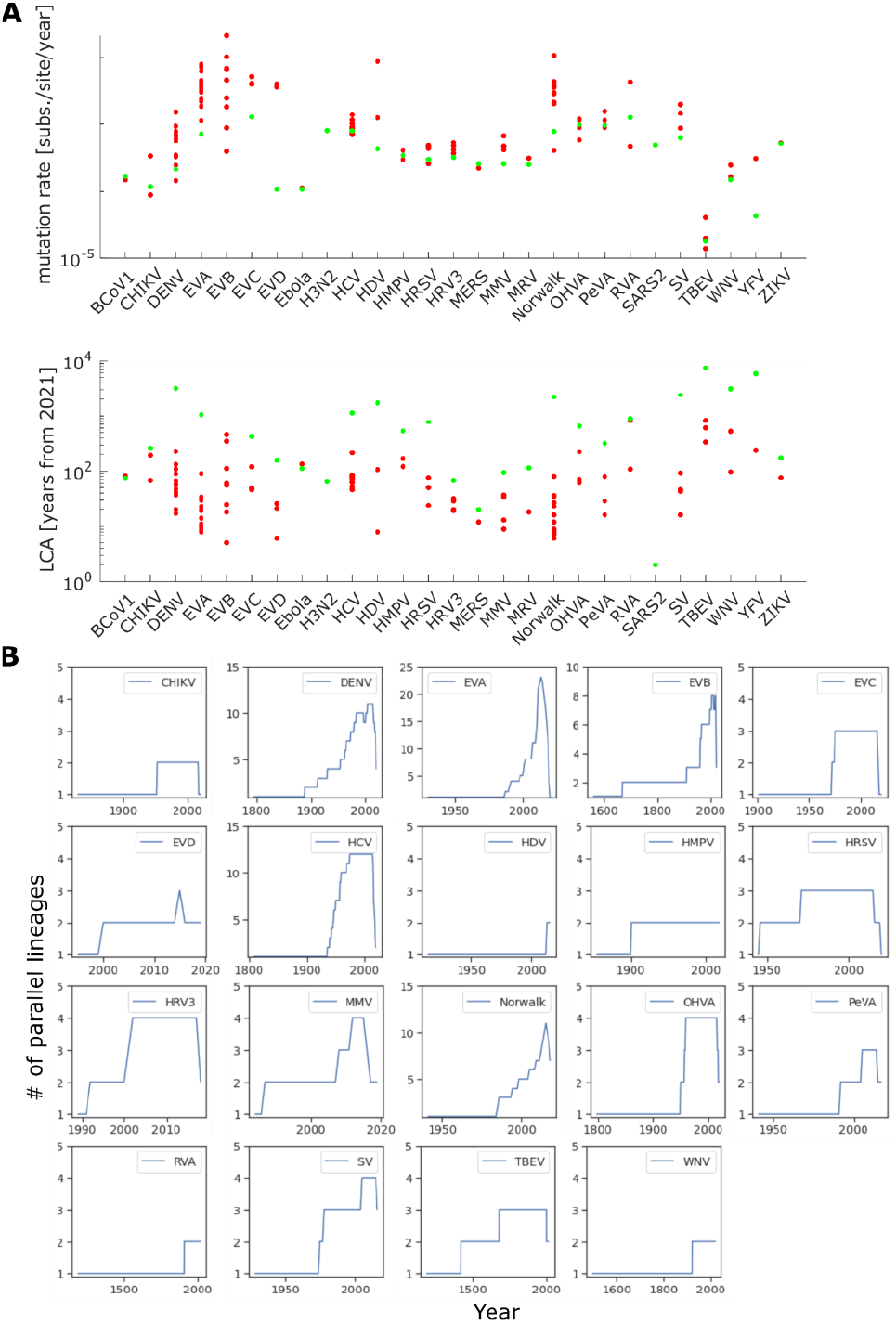
Genealogical lineages of human pathogenic RNA viruses throughout time. **A**. Mutation rates (substitutions per site per year) for all main and GL phylogenetic trees used to construct genealogical trees (top). Time to Last Common Ancestor (LCA) in years from 2021 for all main and GL populations used to construct genealogical trees (bottom). **B**. Number of GLs per virus species circulating at the same time, based on the genealogical trees for each GL. For unannotated samples within a GL, the sampling date was estimated based on the date of the MRCA and root distance.

### Most virus lineages are not regionally or seasonally defined

Perhaps the simplest explanation for the existence of distinct niches would be regional separation. To assess the role played by regionality, we examined whether isolates within a single GL clustered by region. The great circle map distance between pairs of isolates within each GL and between pairs of GLs was computed to retrieve the mean intra-and inter-GL distance, respectively (see Brief Methods and Extended Methods in the SI Appendix for details). Given that GLs do not span the entire phylogeny for all viruses; were defined algorithmically without the incorporation of metadata beyond the date of isolation; and typically include a small number of isolates, we additionally examined the regionality of larger clades, usually defined by serotype or genotype (“Manual Lineages”, ML; see Brief Methods and Extended Methods in the SI Appendix for details).

The ratio of the inter-lineage to intra-lineage map distances is expected to be greater than unity for regionally defined GLs or MLs, and near or below unity for those lineages that are not regionally defined (Fig. S8). For most viruses analyzed, niches do not appear to be regionally defined, with a few notable exceptions (Fig. S8). In particular, YFV is known to split into three regional lineages (East/Central Africa, West Africa, and South America) although the underlying mechanisms for this separation, especially between the African lineages, are not well understood (10, 11). Similarly, Chikungunya virus (CHIKV) displays regionality although in this case the separation seems to be incomplete (32). WNV lineage 1 can be found globally whereas all other lineages are regional (33). However, evidence of the local co-existence of multiple WNV subtypes (34) indicates that additional WNV niches not linked to regionality might exist. HDV displays weak regionality that might be determined by its helper virus, HBV, on which HDV depends for reproduction. The interplay between HDV and HBV genotypes is not yet well understood (35). Ebola outbreaks show a clear regional structure, which is due to *de novo* spill-over events for each outbreak as well as successful containment measures (30). Some, but not all, GLs for DENV, EVA and EVB may be regionally defined (Fig.S8). Thus while common, regionality does not appear to explain the existence of most apparent niches, although this does not imply absence of spatial separation of localized outbreaks.

Similar to regionality, seasonality could potentially support niche formation. Although the temporal resolution of our analysis was limited by the amount of metadata available and the precision with which dates of isolation are specified, we found no evidence that seasonality plays a role in lineage maintenance within viral species. We observed no bi-annual or longer global periodicity of any GLs, but rather a continuous distribution of lineages through time (Figs. S2-7) although shorter temporal patterns are likely for respiratory viruses (36).

GLs could represent localized outbreaks (phases of enhanced virus spread) whereby a virus infects only a minute fraction of the global susceptible population at any given time. Under these conditions, even lineages which do not form distinct immunological niches and do not admit near-simultaneous infection, can coexist within short distances of one another. Infection or vaccination leading to life-long immunity as observed, for example, in the case of MMV or MRV (14), can support the emergence of localized outbreaks. In these cases, naïve hosts are born and are not vaccinated, so that a local community of susceptible hosts emerges. Given sufficient evolutionary distance, lineages can become so diverse antigenically that they form different serotypes, which induce weak to no cross-immunity against each other and thus admit near-simultaneous infection. This pattern has been reported for some picornaviruses (12). In the case of zoonotic viruses, distinct lineages can originate when the same virus species is introduced from different animal reservoirs, which could support ongoing diversification and lineage turnover not observed in the human population. This is how some Orthohepevirus A (OHVA) GLs (37) and possibly some TBEV and WNV GLs (33, 38) could originate. However, even in this case, the maintenance of multiple niches with low turnover within human populations requires spatiotemporal or immunological separation. Regardless of the specific mechanisms underlying the apparent coexistence of non-competing GLs, we sought to explore lineage-defining mutational signatures and to establish whether significant differences existed between the distributions of mutations within and between lineages.

### Selective pressures acting on human RNA viruses

Selective processes are often categorized as diversifying, positive, or purifying, in contrast to neutral evolution via genetic drift (39-41). We sought to probe the selective pressures involved in human pathogenic virus evolution by estimating the ratio of non-synonymous to synonymous substitution rates (*dN/dS*), a gauge of protein-level selection(42, 43). Given that different genes are subject to distinct selective constraints and pressures, the *dN/dS* value was estimated separately for each viral protein-coding gene (44). Seeking to identify defining features of lineage emergence and maintenance, we would ideally estimate *dN/dS* across deep and shallow portions of each phylogeny separately. However, because most GLs antedated modern sequencing technologies and therefore few samples located near the root were available, this approach was not feasible. To partially compensate for this lack of data, we compared *dN/dS* ratios for whole trees, which include deep branches connecting GLs, with those computed over each GL and ML (which are typically larger) individually (see Brief Methods and Extended Methods in the SI Appendix for details). The *dN/dS* estimates for whole trees ranged between 0.02 and 0.5 for most virus protein-coding genes, which is indicative of strong to moderate purifying selection, in line with previous results (45) (Fig. S9). The few virus genes with elevated *dN/dS* ratios encode proteins that are either presented on the virion surface, such as HRSV G and M-2 (∼3.5x above the species mean *dN/dS*), or HMPV SH and G (∼3 and 5x above the species mean, respectively) (Figs. S9, S10), or are involved in interactions with the host immune system, for example, MMV V protein (46) (∼4x above the species mean) (Fig. S10). These proteins are likely to experience positive selection, as described, for example, for HMPV G, where sites under positive selection were identified in the putative ectodomain (47). Elevated *dN/dS* values were also observed for some very short proteins, for example, the 6k peptide of DENV (Fig. S11). However, such observations are sensitive to statistical artifacts and should be interpreted with caution. For OHVA ORF3, the *dN/dS* estimate was ∼4x above the species mean (0.3, Fig. S13), suggesting that this gene, which encodes an ion channel, plays a role in host adaptation following zoonosis (48).

Next, we computed *dN/dS* for each GL and ML individually (Figs S10-13 and Figs. S14-17, respectively). Despite considerable differences in size, generally, the results for GLs and MLs were comparable. For 12 of the 27 viruses studied (members of the order *Mononegavirales*, HMPV, HRSV, HRV3, MMV and MRV; some flaviviruses ZIKV, YFV, TBEV, YFV; HDV; MERS; and CHIKV) the *dN/dS* estimates for individual proteins as well as the mean for the whole tree differed little relative to the respective estimates for individual lineages (Figs S10-17), with no indication of how selective pressures might have varied over time for any genes. In contrast, the GLs of enteroviruses (EVA-D) show elevated *dN/dS*, mainly among capsid proteins (Fig. S16). Although frequent recombination among enteroviruses necessitates interpreting these results with caution (49), this finding, coupled with the observation that mutations in enterovirus capsid protein genes appear to be the primary lineage-defining features (see below), suggests a substantial change in the selective pressure acting on the capsid proteins between the periods of lineage emergence and subsequent maintenance. Notably, OHVA lineages show similarly elevated *dN/dS* for DUF3729 (up to 0.4) and ORF3 *dN/dS* (up to 0.3, which is also elevated relative to the species mean as discussed above) (Fig. S17). Both these genes are likely to be involved in host adaptation following zoonosis (37, 48). Further, a two to five fold increase in mean *dN/dS* was detected for DENV, WNV and HCV GLs across most genes relative to the complete phylogeny (Fig. S15). The interpretation of genome-wide elevation of *dN/dS* in GLs is more challenging and depends on whether the GL is newly emergent, possibly reflecting a period of rapid host adaptation and intense positive selection (45, 50). Given the distant dates predicted for the LCAs for these GLs and lack of lineage turnover, reduced selective pressure moving from stronger purifying selection towards neutral drift appears more likely. Overall, *dN/dS* analysis revealed little about potentially differing selective pressures acting within and between GLs despite the apparent differences in substitution rates critical to the definition of the GLs themselves (as discussed above).

### Intra- and inter-lineage mutational signatures

Gene scale *dN/dS* analysis is often unable to uncover positive selection acting on specific sites or neighborhoods, which can occur in widely different backgrounds, from neutral drift to strong purifying selection (51). Identification of individual positively selected mutations can provide additional insight into differences between the evolutionary contexts of GL emergence and subsequent maintenance. Multiple, parallel non-synonymous mutations comprise the most obvious indication of site-wise positive selection. With the prominent exception of SARS-CoV-2, for which we have previously identified up to 100 sites with recurrent amino acid replacements that are likely subject to positive selection (52), too few recurrent amino acid substitutions were detected for comparable analysis in the remaining viruses analyzed here despite being the species with the largest number of genome sequences available.

Given the infeasibility of the direct, site-specific approach, we performed a genomic neighborhood analysis to compare inter- and intra-lineage mutational signatures. First, amino acid sites were labelled according to three categories of amino acid substitutions (Fig. 4A; see Brief Methods and Extended Methods in the SI Appendix for details): 1) Multiple, deep (MD) substitutions which are “lineage defining”, being conserved in at least 90% of the samples within at least two GLs, but represented by different amino acid residues in each of these GLs. For example, consider the third amino acid of the CHIKV ORF gp1, 97% of the sequences in GL 1 contain a serine in that site, whereas 96% of the sequences in GL2 contain a proline in that site. 2) Multiple, shallow (MS) substitutions occurring on multiple, independent occasions across GLs. 3) All shallow (AS) substitutions occurring at least once within a GL, representing all “recent” events. We then computed site densities for each of these three categories over a sliding window of 101 amino acid sites, respecting protein boundaries.

**Figure 4.**
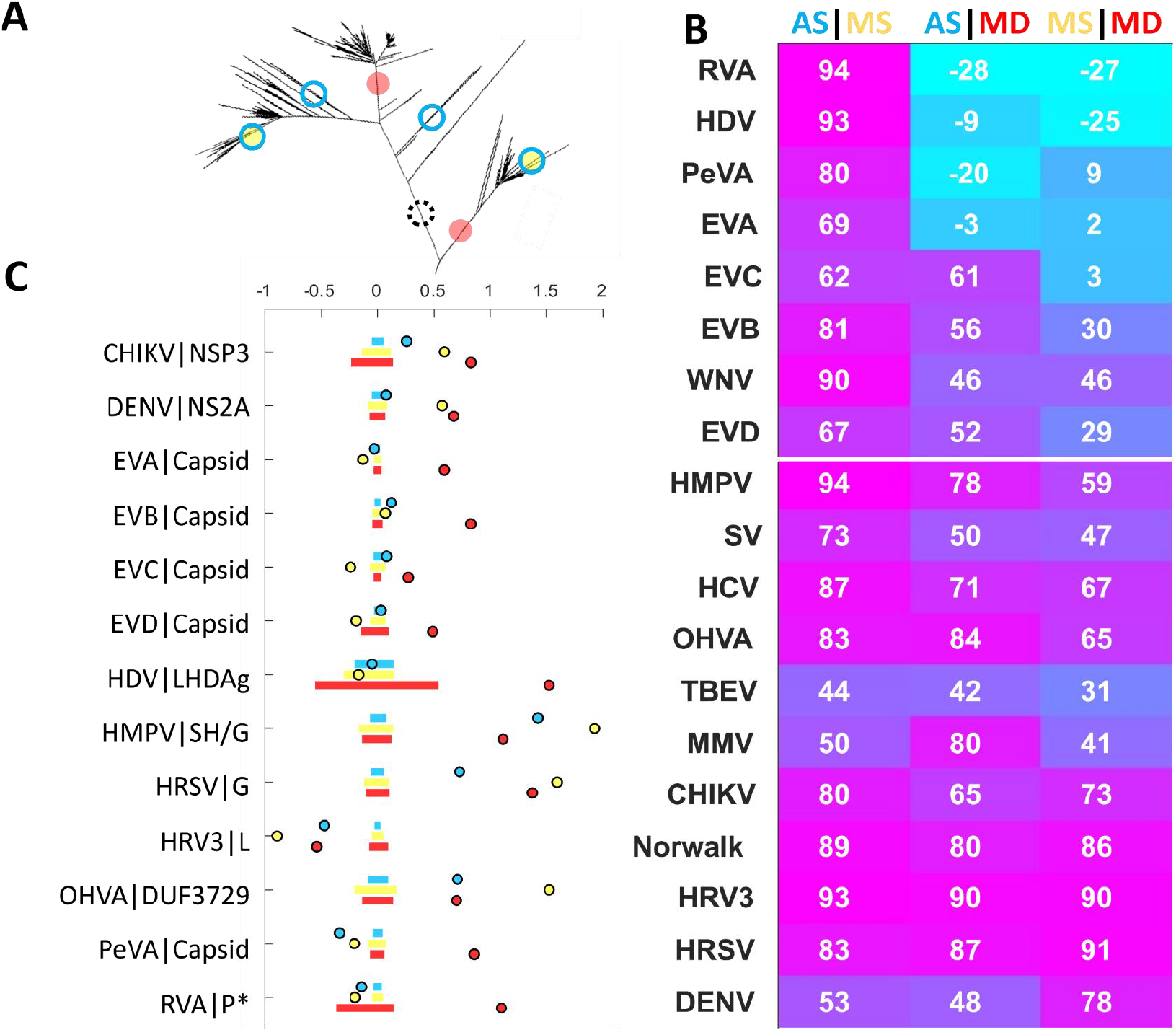
Mutational Signatures in human pathogenic RNA viruses vary little with tree depth. **A**. Illustration of three amino acid site classes considered. 1) Multiple, deep (MD, red). **B**. Multiple, shallow (MS, yellow). 3) All, shallow (AS, blue). Dashed circle represents deep, singular mutations which are excluded from this analysis. **C**. Pearson correlation coefficient between site densities for all pairs of site classes across the genome computed over a moving average of 101 amino acids respecting mature peptide boundaries. Rows are sorted by the first column subtracted from the third column. **D**. Log ratio of the mean site density across the specified peptide relative to the whole genome for select peptides. Bars are bounded by the 25^th^ and 75^th^ percentiles of simulated data drawn from the binomial distribution with *n=total number of sites across the genome* trials of probability *p=length of peptide / length of genome*. RVA|P* represents the union of peptides P2-C, P-3A, and protease-3C.

We then examined the correlation between the site densities in these categories of amino acid substitutions across the genome (Fig. 4B and Figs. S18-24). In most cases, there was a strong, positive correlation between all three categories of amino acid substitutions, indicating that most genomic regions are subject to similar selection pressures during inter- and intra-lineage evolution, with several notable exceptions (Fig.4C). For enteroviruses, we observed an elevated MD site density within capsid proteins (VP1-4, Fig. S18-20), suggesting that capsid mutations are primarily lineage-defining, but become less frequent once an epidemiological niche is established and occupied. This trend is consistent with the historical classification of enterovirus lineages by serotype, which is determined by the antigenic properties of the capsid proteins (53). As mentioned above, frequent intra- and inter-species recombination among all 4 types of enteroviruses requires caution when interpreting these results (49). This trend was similarly observed for the capsid proteins of the picornavirus Parechovirus A (PeVA) (VP0, 1C and 1D, Fig. S23).

Elevated MD and MS substitution densities were observed for HRSV glycoprotein (G), potentially suggesting multiple residues evolving under positive selection throughout the entire course of evolution (including both lineage emergence and maintenance) of this immunologically exposed protein(54). Both MD and MS substitution densities are also increased in DENV NS2A and CHIKV nsp3. DENV NS2A is involved in virus replication and assembly and shows viroporin-like properties(55, 56). A detailed functional understanding of CHIKV nsp3 is lacking although this protein is known to be part of the replication complex and is also involved in modulating the host cell’s antiviral response (57). In line with the observation of elevated *dN/dS* in for OHVA DUF3729 in GLs compared to the whole population, MS substitution densities are elevated in this gene (Fig. S22), suggesting that this poorly characterized protein contains multiple positively selected residues. These residues might have played a role in relatively recent host adaptation, but were not necessarily involved in the emergence of multiple lineages.

The high MD substitution density observed for large human delta antigen (LHDAg) might result from statistical fluctuations given the short length (20 aa) of this peptide and should be interpreted with caution. As observed for the *dN/dS* analysis, we found few mutational signatures, which would shed light on different selection pressures acting within and between GLs. These observations seem to suggest that, although the tempo is variable, the mode of molecular evolution is broadly conserved from the deep to the shallow portions of each phylogeny, thus, spanning considerable evolutionary distances

### Effective population size of human pathogenic RNA viruses

Another tool to indirectly assess the selective pressures shaping a phylogeny is to estimate the effective population size (*Ne*), which defines the time scale of population turnover across generations and thus can reveal major evolutionary events including population bottlenecks (58). Assuming an evolutionary model, such as Wright-Fisher (59) or Moran (60), one can estimate the number of individuals per generation (that is, *Ne*) required for the observed rate of turnover in an idealized population. In what follows, we refer to “selection” as the sum of evolutionary pressures that promote lineage turnover. Although the background could vary from strong purifying selection to neutral drift, the occurrence of lineage turnover implicitly assumes some degree of positive selection in most scenarios. In the context of lineage turnover, under strong selection, *Ne* is small, whereas lack of competition leads to larger *Ne* values over time (17). *Ne* can be inferred from the coalescence rate (*Cr*) estimated for any genealogical tree (17, 61, 62): *Ne* ∗ *t* ∼ 1/*Cr* where *t* is the viral generation time (the time in days a virus needs to complete a transmission cycle from human to human). This expression enables a measurement of diversity and strength of selection among phylogenies represented by a single GL (e.g. H3N2, SARS-CoV-2) as well. Further, the census population size *N* (individuals present at each generation) can be estimated as *N* = *D ∗ t* where *D* is number of yearly cases estimated. The *N/Ne* ratio may be used to quantify lineage turnover, where *N/Ne* >> 1 indicates population bottlenecks, and *N/Ne* ∼ 1 suggests stable population diversity (17). As tree topology, and hence *Ne* estimates, depend on selection strength and sampling effort (17, 58), we directly assessed the effect of sampling by randomly drawing up to either 10 or 100 samples per year, with three replicates each, for H3N2 and EVA as representative viruses with fast and slow turnover respectively. We then used these reduced ensembles of isolates for genealogical tree construction (see Brief Methods and Extended Methods in the SI Appendix for details, mutation rates and time of LCA shown in Fig. S25). We constructed two additional ensembles for each virus composed of the same number of isolates selected above, this time maximizing the sequence diversity (as measured by the hamming distance between alignment rows, see Brief Methods and Extended Methods in the SI Appendix for details). As a result, we obtained six trees evenly sampled over time (3x, e10 and e100) as well as two maximally diverse subtrees of the same size (d10 and d100) for both H3N2 and EVA.

We calculated the coalescence rates for all complete trees and for the H3N2 and EVA subtrees using the PACT package (http://www.trevorbedford.com/pact(17)) and estimated *Ne* (Fig. 5A). The *Ne* estimation was not performed for some zoonotic viruses including TBEV and WNV, for which the generation time could not be reliably estimated. *N/Ne* ratios were calculated based on the best estimates of *Ne* for complete trees and those obtained after even and diverse sampling (H3N2 and EVA), (Fig. 5A and 5B). The estimated *Ne* values span more than two orders of magnitude with H3N2e10 and EVD representing the extremes (Fig. 5A, *Ne* of around 400 and 270,000 respectively). Sampling was found to have a major effect on the estimates. The *Ne* estimates for EVA d100, d10, e100, and the whole tree were similar, whereas EVA e10 was about an order of magnitude lower. It should be noted that EVA e100 contained most sequences present in the whole tree. This trend was even more pronounced for H3N2 where the *Ne* estimates for d100, d10, and the whole tree were similar, whereas those for e10 and e100 were several orders of magnitude lower. It is first important to note that the estimates were not sensitive to the number of sequences present in the phylogeny as illustrated by the equivalency of d10 and the whole phylogeny for both viruses, indicating that the differences observed between e10 and e100 and the whole phylogeny are not merely methodological artifacts. However, the estimate is sensitive to sampling and, as could be expected, this sensitivity is more pronounced for viruses with fast lineage turnover. While perhaps unsurprising, this finding implies that *Ne* has been, and likely continues to be in this work, underestimated due to data limitations for most viruses and H3N2 in particular.

**Figure 5.**
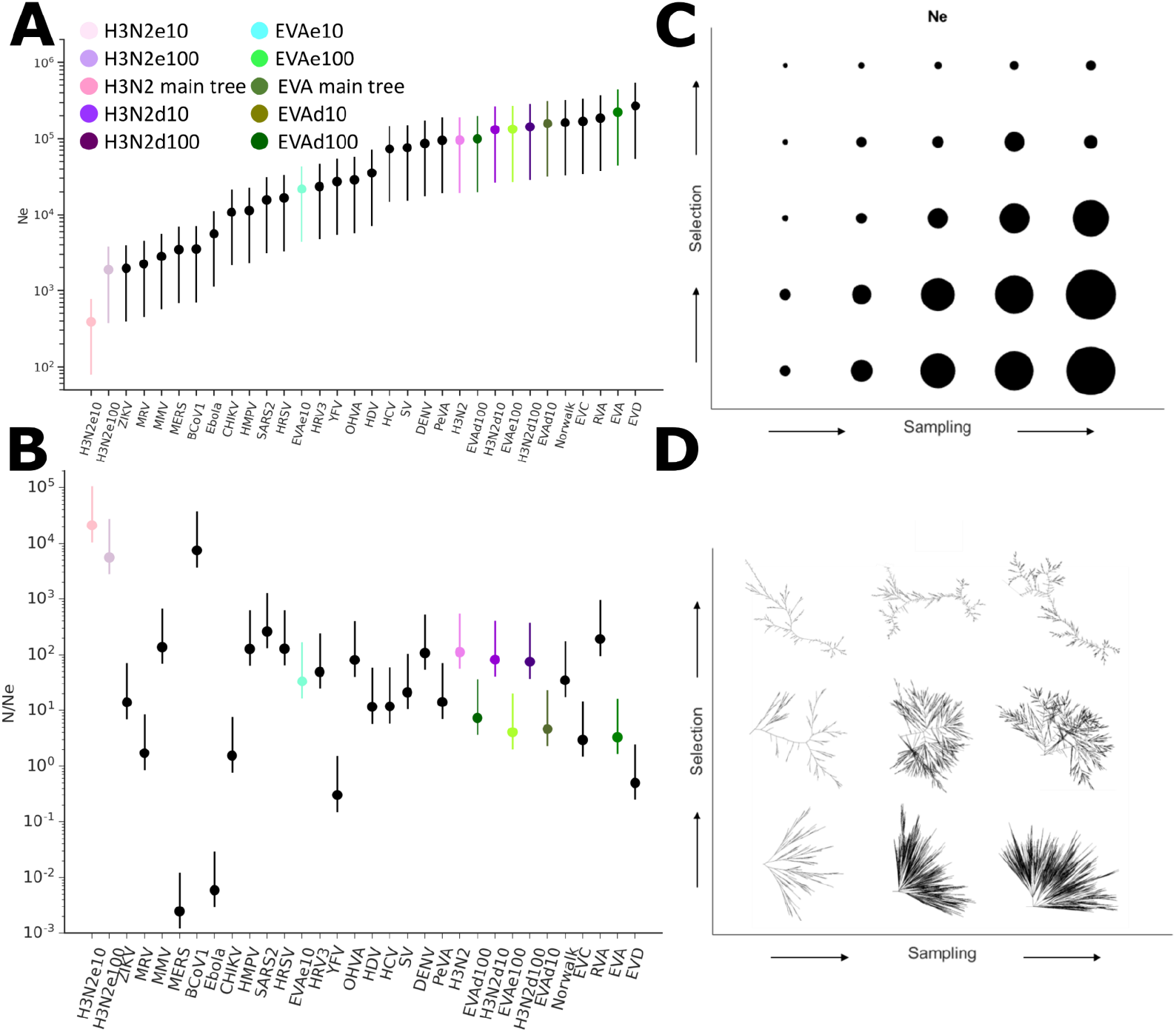
Estimation of effective population sizes for human pathogenic RNA viruses. **A**. *Ne* estimated for genealogical trees. Bars represent varying generation time *t* for each virus ranging between 0.5 and 5 times the value corresponding to the filled circle. For H3N2 and EVA, Ne estimates for evenly sampled trees (up to 10 or 100 samples per year, e10 and e100, respectively) and diverse sampled trees (d10 and d100) are also displayed. **B**. *N/Ne* ratios, where *N* is the census population. Bars represent varying *N* between 0.5 and 5 times the value corresponding to the filled circle. *N* and *t* estimates are shown in Supplementary Data, Table S10. As in (C), *N/Ne* estimates are shown for evenly and diverse sampled trees for EVA and H3N2. Colour code as in (C). **C**. *Ne* (with *t* fixed to 1 for all trees) for simulated trees varying selection strength and sampling density (*Ne* represented by circle area). **D**. Example simulated trees resulting from simulations of varying selection strength and sampling density.

To illustrate the potential equivalency of reduced sampling and increased selection on *Ne* estimation, we simulated an ensemble of genealogical trees under a simple phenomenological model. Trees were iteratively constructed through the addition of clades representing local sequencing efforts. Increased sequencing efforts were modelled by changing the number of sequences in each clade from 2 to 6. Increased selection strength was modelled by changing the placement of these clades on the tree, relative to the root, from the prior iteration. A root distance threshold was set to be the 0th, 25th, 50th, 75th, or 95th percentile of the root distance distribution for all leaves at the prior iteration with higher thresholds corresponding to new isolates being placed farther from the tree root and representing increased selection (Fig. 5C, Fig. S26A-C). Although selection and under-sampling result in qualitatively different tree topologies (Fig. 5D), their effects cannot be disentangled from *Ne* analysis alone. Furthermore, sensitivity to under-sampling is more pronounced under high selection than under low selection (Fig. 5D, Fig. S26D). These effects must be considered when evaluating the expectation that genetic diversity (and hence *Ne*) plateaus earlier in a growing census population *N* when selection is strong (17). This challenge is reflected in the damped increase of *Ne* from H3N2 e10 to e100 when compared to the increase from EVA e10 to e100 (∼4- and 8-fold, respectively; Fig. 5A).

Keeping these sensitivities in mind, we proceeded to examine the *N/Ne* ratios. High *N/Ne* ratios can be an indicator of population bottlenecks. The highest *N/Ne* ratio was observed for H3N2e10 (e100 was similar); in contrast, the estimate for the whole phylogeny was about 200-fold lower (Fig. 5D), within the range of the majority of the other viruses. Thus, sampling efforts can substantially affect the interpretation of the *N/Ne* estimation, moving H3N2 from an outlier associated with extreme bottlenecks to typical behavior. As discussed above, it has been well established that H3N2 is subject to pronounced population bottlenecks as a result of alternating periods of stasis and rapid host adaptation (16-18). However, the results presented here emphasize that on shorter timescales the transmission dynamics of local outbreaks play a larger role in determining the extent of the diversity of the H3N2 population (63) as was the case for the majority of viruses studied in this work,. BCoV1 also demonstrated a high *N/Ne* ratio, within the range of H3N2 e10 and e100 (Fig. 5B). Although this observation could simply result from insufficient sampling, given the high incidence of this virus (Fig. S27D), it might point to pronounced population bottlenecks during the evolution of the BCoV1 population (although not of comparable magnitude to those for H3N2; see below). In contrast, other viruses seem to experience less severe bottlenecks and maintain greater genetic diversity (e.g. MRV and enteroviruses A, C, D). The low *N/Ne* values for MERS, ZIKV and CHIKV likely result from an underestimation of *N* due to large animal reservoirs that might impact estimates of *N* for H3N2 as well.

Whereas insufficient sampling can lead to an underestimation of both *N* and *Ne*, the complete unavailability of genomes from premodern periods can, perhaps counterintuitively, lead to an overestimation of *Ne*. As discussed above, GLs are separated by periods (branches of the tree) with apparently different substitution rates. These branches are often deep within the tree topology and sparsely populated with leaves (if at all), making the assignment of a global model for substitution rates statistically dubious. This can result in inaccurate deep branch lengths for genealogical trees and substantially change the predicted date for the LCA. This date, as well as the predicted dates of other deep nodes, is used to estimate the effective population size. Given these limitations, we sought to establish a lower bound for the effective population size, which still preserves all GLs, through the construction of truncated global genealogical trees or “grafted trees”. The LCA of each grafted tree is set to the LCA of the oldest GL and the remaining GLs are connected to this (multifurcated) root through branches preserving the LCA of each respective GL (see Fig. S27A and Brief Methods or Extended Methods in the SI Appendix for details). We proceeded to estimate Ne for each grafted tree as well as for each GL separately (Fig. S27B). By construction, *Ne* estimates for grafted trees are generally significantly smaller than those for complete trees and larger than those for individual GLs. Notably, the *Ne* estimate for H3N2 (which is represented by a single GL) is the second highest value observed (after RVA) among the viruses studied when this lower bound is considered. This counterintuitive finding emphasizes another facet of the sensitivity of *Ne* estimation to data availability.

These sensitivities are evidently greater within individual GLs, which represent only a subset of the viral diversity for each species. These limitations notwithstanding, in an effort to characterize lineage turnover within individual GLs, we analyzed skyline plots representing the time to the most recent common ancestor (TMRCA) of all clades present at a given time point and diversity within the population over time (as measured by the average time for any two isolates to coalesce, that is, to find their most recent common ancestor) for individual GLs and the complete population (Fig. S28). The average population diversity can be displayed as the mean diversity per year (Fig. S28A). Populations with high turnover, such as H3N2, show a low average diversity per year. whereas those with low turnover are characterized by high diversity. In general, the skyline plots and mean diversity values for complete phylogenies correspond well to *Ne* and *N/Ne* estimations, supporting slow lineage turnover for most of the viruses analyzed. For example, H3N2 displays ∼4x and ∼8x lower mean diversity/year compared to BCoV1 and ZIKV respectively. This observation suggests BCoV1, despite having high *Ne* and *N/Ne* values, has a slower population turnover compared to H3N2. Of note, evidence of high intra-GL turnover was obtained for few GLs (as demonstrated by a mean diversity in the range of H3N2 or BCoV1). The two principal examples are HRSV GL2 and Norwalk GL3 (Fig. S28C, D). The majority of the GLs show mean diversity within the range of viral populations with low turnover.

## Discussion

Here we present a comprehensive phylodynamic analysis of monopartite human pathogenic RNA viruses (and H3N2 hemagglutinin) in an effort to establish global trends in viral evolution in human populations. Despite data limitations, the viruses studied in this work span a wide variety of viral life history characteristics. This lends considerable generality to the study, while making it outside the scope of this work to investigate many features specific to individual lifestyles (for example, intra-host diversity, symptom characteristics, or acute vs. chronic infection (2)). Given this diversity, the commonalities we observe among the virus phylogenies constructed are notable. Consistent with the conclusions of previous efforts (45, 64), we observed moderate to strong purifying selection among all viruses.

Nearly all virus populations are characterized by low rates of lineage turnover, and most consist of multiple, coexisting GLs, monophyletic groups of sequences separated by periods with apparently different substitution rates. Despite these differing substitution rates, *dN/dS* and genomic neighborhood analysis revealed little about how selective pressures might have differed between the early period of GL formation and the subsequent period defined by persistent coexistence. This lack of resolution seems to suggest that, although the tempo is variable, the mode of molecular evolution is broadly conserved from the deep to the shallow portions of each phylogeny, spanning considerable evolutionary distances. The distribution of lineage-defining mutations across the virus genome is similar to that of shallow, repeated mutations for almost all viruses, indicating that positive selection affects sites in the same neighborhoods during both periods (Fig. 4). The lineage-defining role of enterovirus capsid proteins was the principle exception observed, in line with the traditional serotype classification (12). Other virus proteins with different intra- and inter-lineage mutational signatures, which might provide insight into ongoing host and/or vector adaptive evolution, are OHVA DUF3729, DENV NS2A, and CHIKV NSP3. In the case of CHIKV E1-A226V, NSP3 has been demonstrated to play an important role in the adaptation to the vector *Aedes albopictus* (65). In general, the low GL turnover and broadly stable mutational signatures appear to be indicative of weak if any competition among GLs, suggesting that each GL occupies an independent epidemiological niche (Fig. 5).

Such niches could be maintained in a variety of ways, the most obvious possibility being regionality and/or seasonality. Although these factors can explain the persistence of some GLs identified in this work, the majority do not show regional localization, and none display biannual (or more coarse-grained) temporal trends (the limit of time resolution we can reliably detect). At sufficiently large evolutionary distances, niches can be defined by immunological differences, which can overcome immune cross-protection allowing the same host to be infected by multiple subpopulations largely independent of prior infections as seems to be the case for picornaviruses and HRSV (54). These effects are also insufficient to account for the stability of most GLs. We suggest that, in many if not most cases, niches are maintained through a series of localized outbreaks such that only a small fraction of the global susceptible population is infected at any given time. Under this scenario, even lineages that do not overcome immune cross-protection can coexist within short distances of one another. Furthermore, extensive environmental transfer or fragmented animal reservoir populations could play a role. Virions that can persist for extended periods of time outside of the (identified) host or vector might maintain the genetic diversity of a lineage during time periods when no active infections from that lineage occur.

As a result of globalization, disparate communities are becoming increasingly connected, which might lead to increased interaction between previously separated lineages, enhancing between-lineage competition within the viral population. This effect has been demonstrated already for DENV in Thailand where multiple lineages typically coexist throughout the country with a well-defined pattern of dissemination. However, within densely populated areas of Bangkok, genomic analysis pointed to increased competition and lineage turnover (66).

## Conclusions

Phylodynamic analysis revealed multiple co-circulating lineages (GLs) for the majority of human pathogenic RNA viruses separated by periods of apparently different substitution rates within the phylogeny. *dN/dS* and genomic neighborhood analysis yielded surprisingly little evidence of different selection pressures acting within and between GLs, suggesting that whereas the tempo is variable, the mode of molecular evolution is broadly conserved. This slow lineage turnover suggests each GL occupies an independent epidemiological niche, with little inter-GL competition. No pronounced patterns of regional or temporal separation of the GLs were detected, suggesting that the stability of the GLs primarily stems from limited outbreaks within small communities so that only a small fraction of the global susceptible population is infected at any time. These results raise the, perhaps pressing, question: how will increased host-host contact resulting from globalization affect viral evolution? Could new or renewed competition emerge among lineages of endemic viruses to drive diversification, evolution of increased pathogenicity, or even virus speciation? With these questions in mind, we emphasize that, in addition to zoonotic events, the ongoing surveillance of familiar, endemic viruses deserves global attention in effort to mitigate or prevent future pandemics.

## Author contributions

PM, NDR, and GF collected data; PM, NDR, YIW, GF, FZ, and EVK analyzed data; PM, NDR, and EVK wrote the manuscript that was edited and approved by all authors.

## Acknowledgements

The authors thank Koonin group members for helpful discussions. NDR, YIW, PM, and EVK are supported by the Intramural Research Program of the National Institutes of Health (National Library of Medicine).

## Brief Methods

### Multiple Sequence Alignments of RNA Virus Genomes

Genomes were retrieved for all viruses except influenza A virus H3N2 and SARS-CoV-2 from NCBI virus (24). Members of related viral families were used to construct an outgroup when possible. Influenza A virus H3N2 (flu H3N2) segment HA was retrieved from the NCBI flu database (67). The SARS-CoV-2 tree and alignment analyzed in this work was subsampled from a larger alignment consisting of all high quality genomes that were available as of January 8, 2021 in the GISAID database(25), as previously described (52). Subsampling was conducted to maximize the sequence diversity. Acknowledgments for the GISAID deposited sequences used in this study are displayed in Supplementary Data, Table S3. Subalignments were considered for H3N2 and EVA, principally for the purpose of effective population size analysis. In all cases sequences were harmonized to DNA (e.g. U was transformed to T to amend software compatibility) and aligned with MAFFT(26), using default settings. Sequences were clustered according to 100% identity with no coverage threshold using CD-HIT (68), and otherwise default settings for MERS and H3N2.

The longest sequence from each cluster was selected as a representative. Exterior ambiguous characters were removed, and sequences with more than 10 remaining ambiguous characters (“N”) were discarded. Outliers based on hamming distance to the nearest neighbor and consensus were identified and removed from the set. Sites corresponding to protein-coding ORFs were then mapped to the alignments and noncoding regions were discarded. Common gaps corresponding to multiples of three nucleotides were maintained as “true” insertions or deletions and mapped into frame if necessary. Unique alignment rows were identified. Samples related to laboratory experiments, vaccine-related sequences and patents were pruned based on an automated keyword search

Dates and locations of isolation are available for many isolates reported as calendar dates and city or country/administrative region of origin. These dates are referenced as calendar dates in the main text and date indices (number of days before/after January 1, 1950) in the supplement. For the regional analysis, the latitude and longitude of each city of origin or a representative city for each country/administrative region of origin was identified from simplemaps (https://simplemaps.com/data/world-cities) (69).

With the exception of SARS-CoV-2 and H3N2, tree topology was optimized using IQ-TREE (27) with the evolutionary model fixed to GTR+F+G4 and the minimum branch length decreased from the default 10e-6 to 10e-7 (options: -m GTR+F+G4 -st DNA - blmin 0.0000001). For SARS-CoV-2, the tree was drawn from the global topology previously described (52). The global H3N2 tree was approximated using FastTree(70) specifying GTR; a 4 category gamma distribution; no support values; and using a previously-constructed maximum diversity subtree as a constraint (compiled at double precision, options: -nt -gtr -gamma -cat 4 -nosupport -constraints). Trees were rooted according to the position of an outgroup when possible and by date or midpoint otherwise.

Viral lineages were both manually selected based on available metadata and algorithmically into correlated-clades we call “genealogical lineages” or GLs. GLs are defined as monophyletic clades with a strong correlation between the sequencing date and the distance to the tree root. Trees were used to construct date-constrained, genealogical trees using least-square dating (with software LSD2) (31). We considered the Shannon entropy of the clade distribution calculated over sliding windows based on the known or estimated date of isolation OR distance to the tree root as an explicit measure of lineage turnover.

Fitch Traceback (71) was used to estimate ancestral states. Three classes of amino acid sites were identified on the basis of the nonsynonymous mutations within each site. 1) Multiple, deep (MD) substitutions are “lineage defining”. 2) Multiple, shallow sites. 3) All shallow sites. We computed the site density of each class over a sliding window to assess signatures of positive selection. Selection pressures were also assessed through *dN/dS* analysis using PAML(44).

The effective population size *Ne* and the ratio of the census population size *N* over *Ne* was estimated as previously described (17) http://www.trevorbedford.com/pact. Viral diversity (average time of any pair of leaves at a given timepoint to find their LCA) and average time to most recent common ancestor (TMRCA) over time were calculated with the PACT package as well. In order to demonstrate the potential equivalency between the impacts of selection strength and sampling density on effective population size, we additionally simulated an ensemble of trees.

## List of abbreviations

AS: All, shallow
BCoV1: Betacoronavirus 1
CHIKV: Chikungunya virus
DENV: Dengue virus
dN/dS: ratio of non-synonymous to synonymous substitution rates
DUF: Domain of unknown function
Ebola: Zaire ebolavirus
EVA: Enterovirus A
EVAd: Enterovirus A ‘diverse’
EVAe: Enterovirus A ‘even’
EVB: Enterovirus B
EVC: Enterovirus C
EVCe: Enterovirus C ‘every’ sample
EVCr: Enterovirus C ‘reduced’ samples
EVD: Enterovirus D
GL: Genealogical lineage
H3N2d: H3N2 ‘divers’
H3N2e: H3N2 ‘even’
HBV: Hepatitis B virus
HCV: Hepatitis C virus
HCVe: Hepatitis C virus ‘every’ sample
HCVr: Hepatitis C virus ‘reduced’ samples
HDV: Hepatitis D virus
HMPV: Human metapneumovirus
HRSV: Human respiratory syncytial virus
HRV3: Human respirovirus 3
H3N2: Influenza A Virus H3N2
LCA: Last common ancestor
LHDAg: Large human delta antigen
LSD: least-square distance
MD: Multiple, deep
MERS: Middle East respiratory syndrome-related coronavirus
ML: Manual lineage
MMV: Measles morbillivirus
MRCA: Most recent common ancestor
MRV: Mumps rubulavirus
MS: Multiple, shallow
N: Census population size
Ne: Effective population size
Norwalk: Norwalk virus
NS: Non-structural protein
NSP: Non-structural protein
OHVA: Orthohepevirus A
PeVA: Parechovirus A
RVA: Rhinovirus A
SARS-CoV-2: Severe acute respiratory syndrome-related coronavirus 2
SV: Sapporo virus
TMRCA: Time to most recent common ancestor
YFV: Yellow fever virus
ZIKV: Zika virus

## Declarations

### Ethics approval and consent to participate

Not applicable.

### Consent for publication

Not applicable.

### Data Availability

The datasets generated and/or analyzed during the current study are available as supplementary data at Zenodo, https://doi.org/10.5281/zenodo.5711959, as well as through FTP, https://ftp.ncbi.nih.gov/pub/wolf/_suppl/virNiches/. Original virus sequences are publicly available for all viruses except SARS-CoV-2 at NCBI virus (24). SARS-CoV-2 sequences are available at GISAID (25).

### Competing interests

The authors declare that they have no competing interests.

### Funding

NDR, YIW PM, and EVK are supported by the Intramural Research Program of the National Institutes of Health (National Library of Medicine).

## Notes

https://doi.org/10.5281/zenodo.5711959

### Competing Interest Statement

The authors have declared no competing interest.

## References

1. Suttle CA (2007) Marine viruses--major players in the global ecosystem. Nat Rev Microbiol 5(10):801–812.

2. Chisholm PJ, Busch JW, & Crowder DW (2019) Effects of life history and ecology on virus evolutionary potential. Virus Res 265:1–9.

3. Johnson PT, de Roode JC, & Fenton A (2015) Why infectious disease research needs community ecology. Science 349(6252):1259504.

4. Yan L, Neher RA, & Shraiman BI (2019) Phylodynamic theory of persistence, extinction and speciation of rapidly adapting pathogens. Elife 8.

5. Marchi J, Lässig M, Mora T, & Walczak AM (2019) Multi-Lineage Evolution in Viral Populations Driven by Host Immune Systems. Pathogens 8(3).

6. WHO WHO (2021) Coronavirus (COVID-19) Dashboard.

7. Harvey WT, et al. (2021) SARS-CoV-2 variants, spike mutations and immune escape. Nat Rev Microbiol 19(7):409–424.

8. Rochman N, Wolf Y, & Koonin E (2021) Evolution of human respiratory virus epidemics [version 2; peer review: 2 approved]. F1000Research 10(447).

9. Simmonds P, Aiewsakun P, & Katzourakis A (2019) Prisoners of war - host adaptation and its constraints on virus evolution. Nat Rev Microbiol 17(5):321–328.

10. Bryant JE, Holmes EC, & Barrett AD (2007) Out of Africa: a molecular perspective on the introduction of yellow fever virus into the Americas. PLoS Pathog 3(5):e75.

11. Li Y & Yang Z (2017) Adaptive Diversification Between Yellow Fever Virus West African and South American Lineages: A Genome-Wide Study. Am J Trop Med Hyg 96(3):727–734.

12. Brouwer L, Moreni G, Wolthers KC, & Pajkrt D (2021) World-Wide Prevalence and Genotype Distribution of Enteroviruses. Viruses 13(3).

13. Geoghegan JL & Holmes EC (2018) Evolutionary Virology at 40. Genetics 210(4):1151–1162.

14. Grenfell BT, et al. (2004) Unifying the epidemiological and evolutionary dynamics of pathogens. Science 303(5656):327–332.

15. Kim H, Webster RG, & Webby RJ (2018) Influenza Virus: Dealing with a Drifting and Shifting Pathogen. Viral Immunol 31(2):174–183.

16. Wolf YI, Viboud C, Holmes EC, Koonin EV, & Lipman DJ (2006) Long intervals of stasis punctuated by bursts of positive selection in the seasonal evolution of influenza A virus. Biol Direct 1:34.

17. Bedford T, Cobey S, & Pascual M (2011) Strength and tempo of selection revealed in viral gene genealogies. BMC Evol Biol 11:220.

18. Koelle K, Cobey S, Grenfell B, & Pascual M (2006) Epochal evolution shapes the phylodynamics of interpandemic influenza A (H3N2) in humans. Science 314(5807):1898–1903.

19. Rota PA, et al. (2016) Measles. Nat Rev Dis Primers 2:16049.

20. Hamblin SR, White PA, & Tanaka MM (2014) Viral niche construction alters hosts and ecosystems at multiple scales. Trends Ecol Evol 29(11):594–599.

21. Volz EM, Koelle K, & Bedford T (2013) Viral phylodynamics. PLoS Comput Biol 9(3):e1002947.

22. Houldcroft CJ, Beale MA, & Breuer J (2017) Clinical and biological insights from viral genome sequencing. Nat Rev Microbiol 15(3):183–192.

23. Liu L (2014) Fields Virology, 6th Edition. Clinical Infectious Diseases 59(4):613–613.

24. Hatcher EL, et al. (2017) Virus Variation Resource - improved response to emergent viral outbreaks. Nucleic Acids Res 45(D1):D482–d490.

25. Shu Y & McCauley J (2017) GISAID: Global initiative on sharing all influenza data–from vision to reality. Eurosurveillance 22(13):30494.

26. Katoh K, Misawa K, Kuma Ki, & Miyata T (2002) MAFFT: a novel method for rapid multiple sequence alignment based on fast Fourier transform. Nucleic acids research 30(14):3059–3066.

27. Nguyen L-T, Schmidt HA, Von Haeseler A, & Minh BQ (2015) IQ-TREE: a fast and effective stochastic algorithm for estimating maximum-likelihood phylogenies. Molecular biology and evolution 32(1):268–274.

28. Scholle SO, Ypma RJ, Lloyd AL, & Koelle K (2013) Viral substitution rate variation can arise from the interplay between within-host and epidemiological dynamics. Am Nat 182(4):494–513.

29. Organisation WH (2021) Ebola virus disease fact-sheets. (who).

30. Jacob ST, et al. (2020) Ebola virus disease. Nat Rev Dis Primers 6(1):13.

31. To T-H, Jung M, Lycett S, & Gascuel O (2016) Fast dating using least-squares criteria and algorithms. Systematic biology 65(1):82–97.

32. Schneider AB, et al. (2019) Updated Phylogeny of Chikungunya Virus Suggests Lineage-Specific RNA Architecture. Viruses 11(9).

33. Chancey C, Grinev A, Volkova E, & Rios M (2015) The global ecology and epidemiology of West Nile virus. Biomed Res Int 2015:376230.

34. Di Giallonardo F, et al. (2016) Fluid Spatial Dynamics of West Nile Virus in the United States: Rapid Spread in a Permissive Host Environment. J Virol 90(2):862–872.

35. Wang W, et al. (2021) Assembly and infection efficacy of hepatitis B virus surface protein exchanges in eight hepatitis D virus genotype isolates. J Hepatol.

36. Moriyama M, Hugentobler WJ, & Iwasaki A (2020) Seasonality of Respiratory Viral Infections. Annu Rev Virol 7(1):83–101.

37. Lin S & Zhang YJ (2021) Advances in Hepatitis E Virus Biology and Pathogenesis. Viruses 13(2).

38. Kubinski M, et al. (2020) Tick-Borne Encephalitis Virus: A Quest for Better Vaccines against a Virus on the Rise. Vaccines (Basel) 8(3).

39. Dolan PT, Whitfield ZJ, & Andino R (2018) Mechanisms and Concepts in RNA Virus Population Dynamics and Evolution. Annu Rev Virol 5(1):69–92.

40. Lynch M, et al. (2016) Genetic drift, selection and the evolution of the mutation rate. Nat Rev Genet 17(11):704–714.

41. Kimura M (1991) The neutral theory of molecular evolution: a review of recent evidence. Jpn J Genet 66(4):367–386.

42. Hurst LD (2002) The Ka/Ks ratio: diagnosing the form of sequence evolution. Trends Genet 18(9):486.

43. Hejase HA, Dukler N, & Siepel A (2020) From Summary Statistics to Gene Trees: Methods for Inferring Positive Selection. Trends Genet 36(4):243–258.

44. Yang Z (2007) PAML 4: phylogenetic analysis by maximum likelihood. Molecular biology and evolution 24(8):1586–1591.

45. Lin JJ, Bhattacharjee MJ, Yu CP, Tseng YY, & Li WH (2019) Many human RNA viruses show extraordinarily stringent selective constraints on protein evolution. Proc Natl Acad Sci U S A 116(38):19009–19018.

46. Takeuchi K, Kadota SI, Takeda M, Miyajima N, & Nagata K (2003) Measles virus V protein blocks interferon (IFN)-alpha/beta but not IFN-gamma signaling by inhibiting STAT1 and STAT2 phosphorylation. FEBS Lett 545(2-3):177–182.

47. Papenburg J, et al. (2013) Genetic diversity and molecular evolution of the major human metapneumovirus surface glycoproteins over a decade. J Clin Virol 58(3):541–547.

48. Primadharsini PP, Nagashima S, & Okamoto H (2021) Mechanism of Cross-Species Transmission, Adaptive Evolution and Pathogenesis of Hepatitis E Virus. Viruses 13(5).

49. Kyriakopoulou Z, Pliaka V, Amoutzias GD, & Markoulatos P (2015) Recombination among human non-polio enteroviruses: implications for epidemiology and evolution. Virus Genes 50(2):177–188.

50. Holmes EC (2003) Patterns of intra- and interhost nonsynonymous variation reveal strong purifying selection in dengue virus. J Virol 77(20):11296–11298.

51. Kryazhimskiy S & Plotkin JB (2008) The population genetics of dN/dS. PLoS Genet 4(12):e1000304.

52. Rochman ND, et al. (2021) Ongoing global and regional adaptive evolution of SARS-CoV-2. Proc Natl Acad Sci U S A 118(29).

53. Oberste MS, Maher K, Kilpatrick DR, & Pallansch MA (1999) Molecular evolution of the human enteroviruses: correlation of serotype with VP1 sequence and application to picornavirus classification. J Virol 73(3):1941–1948.

54. Wertz GW & Moudy RM (2004) Antigenic and genetic variation in human respiratory syncytial virus. Pediatr Infect Dis J 23(1 Suppl):S19–24.

55. Neufeldt CJ, Cortese M, Acosta EG, & Bartenschlager R (2018) Rewiring cellular networks by members of the Flaviviridae family. Nat Rev Microbiol 16(3):125–142.

56. Shrivastava G, et al. (2017) NS2A comprises a putative viroporin of Dengue virus 2. Virulence 8(7):1450–1456.

57. Fros JJ & Pijlman GP (2016) Alphavirus Infection: Host Cell Shut-Off and Inhibition of Antiviral Responses. Viruses 8(6).

58. Wang J, Santiago E, & Caballero A (2016) Prediction and estimation of effective population size. Heredity (Edinb) 117(4):193–206.

59. Ishida Y & Rosales A (2020) The origins of the stochastic theory of population genetics: The Wright-Fisher model. Stud Hist Philos Biol Biomed Sci 79:101226.

60. Moran PAP (1958) Random processes in genetics. Mathematical Proceedings of the Cambridge Philosophical Society 54(1):60–71.

61. Strimmer K & Pybus OG (2001) Exploring the demographic history of DNA sequences using the generalized skyline plot. Mol Biol Evol 18(12):2298–2305.

62. Pybus OG, Rambaut A, & Harvey PH (2000) An integrated framework for the inference of viral population history from reconstructed genealogies. Genetics 155(3):1429–1437.

63. McCrone JT & Lauring AS (2018) Genetic bottlenecks in intraspecies virus transmission. Curr Opin Virol 28:20–25.

64. Wertheim JO & Kosakovsky Pond SL (2011) Purifying selection can obscure the ancient age of viral lineages. Mol Biol Evol 28(12):3355–3365.

65. Tsetsarkin KA, et al. (2014) Multi-peaked adaptive landscape for chikungunya virus evolution predicts continued fitness optimization in Aedes albopictus mosquitoes. Nat Commun 5:4084.

66. Salje H, et al. (2017) Dengue diversity across spatial and temporal scales: Local structure and the effect of host population size. Science 355(6331):1302–1306.

67. Bao Y, et al. (2008) The influenza virus resource at the National Center for Biotechnology Information. J Virol 82(2):596–601.

68. Li W & Godzik A (2006) Cd-hit: a fast program for clustering and comparing large sets of protein or nucleotide sequences. Bioinformatics 22(13):1658–1659.

69. simplemaps (2021) World Cities Database.

70. Price MN, Dehal PS, & Arkin AP (2010) FastTree 2–approximately maximum-likelihood trees for large alignments. PloS one 5(3):e9490.

71. Fitch WM (1971) Toward defining the course of evolution: minimum change for a specific tree topology. Systematic Biology 20(4):406–416.

72. Wolf YI, et al. (2018) Origins and Evolution of the Global RNA Virome. mBio 9(6).

