## Supplementary material for "Human pathogenic RNA viruses establish non-competing lineages by occupying independent niches": SI Appendix

**This PDF file includes:**

Extended Methods
Figures S1 to S29

**Additional supplementary information can be found:**

https://doi.org/10.5281/zenodo.5711959
**or**
https://ftp.ncbi.nih.gov/pub/wolf/_suppl/virNiches/

This repository includes:

Tables

Table S1: Virus characteristics including abbreviation, family, order, tax id, sequence length threshold (for nearly complete genomes), vaccination status, mode of transmission, circulation (human or human/zoonotic), available treatment, and disease progression (acute or chronic).

Table S2: Sequence IDs of outgroup constituents.

Table S3: GISAID acknowledgements for the SARS-CoV-2 sequences used in this study.

Table S4: Summary of manual sequence curation indicating lab- or vaccine-related keywords used to prune sequences.

Table S5: Sequence IDs for EVA and H3N2 e10, e100, d10, and d100 subtrees.

Table S6: Sequence IDs for ML and GL lineages.

Table S7: Sequence IDs for the H3N2 “new” and “old” subtrees.

Table S8: Mutation rates (in units of nucleotide substitutions per site per year) and estimated dates for the last common ancestor of all tree/subtrees. Negative dates represent years prior to 0 CE.

Table S9: Mean dN/dS values.

Table S10: Yearly viral cases and generation time estimations (with references).

Table S11: Metadata including the sequence ID, host, date of isolation, location of isolation, and subtype.

Directories

Alignments ORFeome alignments (excluding stop codons) with the exception of SARS-CoV-2.

Correlated Subtrees Subtrees representing all correlated-clades (genealogical lineages; GL) in

Newick format.

Diversity TMRCA and Skyline Plots .png files.

e/d/n/o Subtrees EVA and H3N2 subtrees, evenly and diversely sampled as well as “early” and “late” subtrees for H3N2.

Genealogical Trees Including global topologies (.main), subtrees, and grafted trees (.grafted).

ORF References The first and last nucleotide of each ORF in the reference sequence.

Redundancy Tables List of all redundant isolates and the corresponding representative in the

ORFeome-unique alignment.

Reference Sequences Genbank page for all reference sequences.

Rooted Trees Global topologies for each virus.
Ultrametric Trees For each global topology.

### Extended Methods

#### Multiple Sequence Alignments of RNA Virus Genomes

Nearly complete viral genomes were retrieved for all viruses except influenza A virus H3N2 and SARS-CoV-2 from NCBI virus (1) with the following minimal length requirements on nucleotide sequence (27000: MERS, BCoV1; 1700: Ebola; 15500: MMV; 12000: HRSV, HRV3, MRV; 11000: HMPV; 9500: TBEV; 8500: DENV, CHIKV, HCV, WNV, YFV, ZIKV; 5500: EVA, EVD, Norwalk, PeVA, SV; 5600: EVB, RVA; 6000: EVC; 4700: OHVA; 1200: HDV; ). Genomes were retrieved on 10/28/2020, except for MERS, which was retrieved on 12/01/2020. Isolates from non-human hosts were included to take into account vector transmission and putative zoonotic reservoirs. Members of related viral families were used to construct an outgroup if sequence identity of the largest protein between outgroup candidate and main group was at least 30%. Up to 20 diverse reference viruses were selected among the outgroup candidates. Outgroup sequence IDs are listed in Supplementary Data, Table S2. For HDV and HRSV, no outgroup was available meeting these criteria.

Samples for Influenza A virus H3N2 (flu H3N2) segment HA were retrieved from the NCBI flu database (2) on 05/18/2021. Only (nearly) complete HA genome segments were considered (larger than 1600 nucleotides). No outgroup was considered forH3N2.

Due to the exceptionally large number of SARS-CoV-2 genomes sequenced by 2021, the SARS-CoV-2 tree and alignment analyzed in this work was subsampled from a larger alignment consisting of all high quality genomes that were available as of January 8, 2021 in the GISAID database(3), as previously described (4). Subsampling was conducted to maximize the sequence diversity (based on Hamming distance). Acknowledgments for the GISAID deposited sequences used in this study are displayed in Supplementary Data, Table S3.

Subalignments were considered for H3N2 and EVA, principally for the purpose of effective population size analysis (see below). For each, subalignments were constructed to sample an even number of isolates from every year at 10 and 100 isolates per year with three randomly sampled replicates (6 alignments in total). Two additional alignments were constructed to maximize the sequence diversity (based on Hamming distance, as for SARS-CoV-2) over the same number of isolates as the first set of subalignments given 10 and 100 isolates per year respectively.

In all cases sequences were harmonized to DNA (e.g. U was transformed to T to amend software compatibility) and aligned with MAFFT(5), using default settings.

Sequences were clustered according to 100% identity with no coverage threshold using CD-HIT (6), and otherwise default settings for MERS and H3N2. The longest sequence from each cluster was selected as a representative. For all other viruses, all sequences were considered for the next steps. Exterior ambiguous characters (preceding/succeeding the first/last defined nucleotide) were removed, and sequences with more than 10 remaining interior, completely ambiguous characters (“N”) were discarded. MAFFT was run with default settings to align the cleared unique sequences. Outliers based on hamming distance to the nearest neighbor and consensus were identified and removed from the set. Remaining sequences were aligned with MAFFT, default settings, together with the respective outgroup.

Note that, in principle, the redundancy introduced by this clustering step can introduce ambiguity in mapping metadata on the tree. We show that this is not an issue for these datasets below.

Exterior ambiguous characters (preceding/succeeding the first/last defined nucleotide) were removed. Sites corresponding to protein-coding ORFs were then mapped to the alignments from the reference sequences excluding stop codons (see Supplementary Data). Noncoding regions were discarded.

These resulting alignments contained out-of-frame gaps. We first identified gaps in the reference sequences corresponding to multiples of three nucleotides present in no more than 90% of all sequences (and thus likely corresponding to “true” insertions relative to the reference at the same position in at least 10% of all sequences). Sites corresponding to less common insertions and non-triplet gaps in the reference were discarded. The remaining gaps represent deletions relative to the reference sequence. Similarly, codons in the frame of the reference sequences where fewer than 10% of sequences in any of the three sites were represented by non-gap characters were removed. Gaps shorter than three nucleotides in the remaining sites were replaced with the ambiguous character N. Longer gaps were shifted into frame and padded with ambiguous characters on either end of the gap to minimize the number of characters changed. Sequences composed of more than 25% gaps were then discarded.

Outliers among the remaining sequences were then identified based on the Hamming distance (excluding gaps and ambiguous characters) to the nearest neighbor and removed

Unique alignment rows were then identified, and redundant rows were discarded. Such redundancy, in addition to the redundancy introduced by prior clustering for MERS and H3N2, can introduce ambiguity in mapping metadata on the tree as a single, random representative isolate is selected to correspond to each unique row. As discussed above, the SARS-CoV-2 data were subsampled and matched with representative metadata accordingly. Among the remaining twenty-six alignments, only 7 had more than 10% of all isolates removed due to redundancy: Ebola (37%), EVC (15%), HRSV (13%), MERS (16%), MMV (20%), MRV (25%), and H3N2 (42%). For EVC, all but 5 redundant sequences mapped to rows in the alignment which were later pruned (see the following section on pruning lab-related isolates and select isolates from non-human hosts). For the remaining 6 it was confirmed that the sequencing dates for the redundant isolates correspond to those of the representative isolates mapped to the tree (see Figure S29) and thus the selection of these representatives does not introduce significant ambiguity. Tables listing all redundant sequence IDs and the sequence ID representing each redundant sequence in the respective alignment are available in the Supplementary Data.

#### Pruning Lab-Related Isolates from Non-Human Hosts

Samples related to laboratory experiments, vaccine-related sequences and patents were pruned based on an automated keyword search (see Supplementary Data, Table S4) followed by manual curation. Further, samples related to vaccine-derived poliomyelitis were removed from the EVC set as well as samples related to patients under treatment with direct-acting antivirals from the HCV set. The supplementary material contains analysis for trees maintaining "every" sequence (EVCe, HCVe) as well as only the "reduced" (EVCr, HCVr) set. Main figures display results for the reduced set.

#### Selecting subalignments for EVA and H3N2

In order to analyze the effect of sampling efforts on the effective population size (see below), several subalignments were generated for EVA and H3N2. To maintain an even representation over time, up to either 10 or 100 isolates (not unique sequences) were randomly selected ("even"/ "e"). To construct subalignments maximizing sequence diversity, we collected the same number of sequences as each evenly sampled subalignment through an iterative process maximizing the minimum hamming distance at each step ("diverse"/ "d"). The complete alignment was reduced to the samples within each subset (keeping the outgroup) to retrieve eight final subalignments per virus (3x e_10, 3x e_100 and 1x d_10 and d100). The corresponding sequence IDs for each subalignment can be found in Supplementary Data, Table S5.

#### Metadata Analysis

Dates and locations of isolation are available for many isolates reported as calendar dates and city or country/administrative region of origin. The fraction of isolates for which metadata is available varies across the alignments. Calendar dates may be reported as the day, month, or only year. If the day or month was not specified, the collection date was assumed to be the midpoint of the window. These dates are referenced as calendar dates in the main text and date indices (number of days before/after January 1, 1950) in the supplement for convenience. In practice, for the analysis presented in this work, only the year collected is relevant.

For the regional analysis, the latitude and longitude of each city of origin or a representative city for each country/administrative region of origin was identified from simplemaps (https://simplemaps.com/data/world-cities) (7). This information was used to calculate the distance on the globe between pairs of sequencing locations (the great circle distance, not respecting geographic or political boundaries).

#### Tree Construction

With the exception of SARS-CoV-2 and H3N2, tree topology was optimized using IQ-TREE (8) with the evolutionary model fixed to GTR+F+G4 and the minimum branch length decreased from the default 10e-6 to 10e-7 (options: -m GTR+F+G4 -st DNA -

blmin 0.0000001). For SARS-CoV-2, the tree was drawn from the global topology previously described (4) corresponding to the selected subalignment. For H3N2, trees for both subalignments (evenly sampled and maximum diversity) were constructed as described using IQ-TREE. The global H3N2 tree was approximated using FastTree(9) specifying GTR; a 4 category gamma distribution; no support values; and using the previously-constructed maximum diversity subtree as a constraint (compiled at double precision, options: -nt -gtr -gamma -cat 4 -nosupport -constraints).

Trees were rooted according to the position of an outgroup selected from representative sequences of the closest relative species whenever the percent identity of the largest protein or polyprotein was at least 30%. In three cases, the outgroup did not meet these criteria. We applied midpoint tooting for HDV andHRSV) and rooted at the oldest sample for H3N2.

**Manual Parsing of Trees**

Viral lineages were manually selected representing viral sero- or genotypes, known geographic subtypes, and large, monophyletic clades. No manual lineages were selected for EVB and Rhinovirus A (RVA). Sequences of each manual lineage (“ML”) are listed in Supplementary Data, Table S6.

**Parsing Trees into Correlated Subsets Defined by Sequencing Date and Root Distance (“genealogical lineages”)**

In addition to these manually established subtypes, we sought to identify individual “outbreaks” within the trees with a well defined evolutionary “trajectory” composed of a monophyletic clade such that within this clade, the distance to the tree root is well correlated with the date of isolation.

Moving from root to tip, at each node of the global topology, we calculated the Pearson correlation coefficient between the root distance and the isolation date for all sequences descendent from that node. For a predefined threshold, *C*, if the correlation was greater than *C*, this clade was added to the ensemble of correlated clades and the search was stopped at that node. If the correlation was below *C*, the search would continue until the descendent clades contained fewer than 15 descendent sequences. Such clades with fewer than 15 sequences were not considered further.

The threshold, *C*, was then determined for each tree to be either 0.8 or the largest value such that 30% of all sequences with known isolation dates were included in a correlated clade, whichever was smaller with the exception of SARS-CoV-2. The threshold for SARS-CoV-2 was set to 0.75 as between 0.75 and 0.8 this parsing scheme dramatically differed from returning a single, inclusive correlated clade to returning over 40 small clades. At 0.75, all but one sequence was included, and consequently we considered all sequences in SARS-CoV-2 to belong to a single correlated clade. *C* was found to be less than 0.2 for two trees (HDV/0.18 and RVA/0.16); less than 0.65 for five more (Ebola/0.59, EVB/0.41, HCV/0.63, PeVA/0.39 and TBEV/0.49); and 0.8 for all others (with the exception of SARS-CoV-2). H3N2 sequences display a very high root distance/sequencing date correlation (Fig. S7) and nearly all H3N2 sequences formed a single correlated clade even at a threshold value near 1 with the method described above; however, due to the ladder-like tree structure with relatively fewer sequences near the root, parsing near the root with this method is sensitive to outliers and rooting. Consequently, we considered all sequences in H3N2 to belong to a single correlated clade. Note as discussed in the main text, we refer to these correlated clades as “genealogical lineages” or GL.

Subtrees corresponding to the subalignments described above for H3N2 and EVA were additionally considered as well as two subtrees for H3N2 representing “newer” and “older” sequences (H3N2n/H3N2o) roughly determined to be sequenced after/before 2010 respectively. Sequences of each correlated/ genealogical lineage (“GL”) are listed in Supplementary Data, Table S6 and sequences for H3N2n and H3N2o are in Supplementary Data, Table S7.

**Construction of Date-Constrained Genealogical Trees**

All trees, global topologies for each alignment as well as the selected subtrees, with branch lengths corresponding to the number of nucleotide substitutions per site, were paired with the known dates of isolation to construct date-constrained, genealogical trees using least-square dating (with software LSD2) (10) associated with a single mutation rate for each tree/subtree and, consequently, a predicted date for the LCA of each tree/subtree (Supplementary Data, Table S8). When not known, the isolation dates were estimated and unconstrained so that, in some trees, select sequences are predicted to have been isolated in the future.

Many of these clades are far from the tree root with few sequences interspersed between the clade root and the tree root. The mutation rate along these long, deep connecting branches may differ significantly from that estimated over the whole tree or among the GLs. If present, such a rate mismatch would substantially change the predicted date for the LCA. This date, as well as the predicted dates of other deep nodes, is used to estimate the effective population size. We are primarily interested in establishing a lower bound, but not necessarily an upper bound, for the effective population size (see below) and given the sparsity of these branches, a partitioned evolutionary model is unlikely to yield statistically significant results. Alternatively, we constructed truncated global genealogical trees, or “grafted trees”, as follows. First the deepest root (oldest LCA) of any GL was identified. This was taken to be the root for the grafted tree. Next, the remaining GLs were connected to this (now multifurcated) root by branches reflecting the difference between the LCA of each clade and the root. These grafted trees represent an upper bound on the mutation rates over these branches and the effective population size estimated over these grafted trees represents the lower bound for the effective population size of each virus respecting the genealogical trees estimated over each GL.

Given the original trees as well as the date-constrained, genealogical trees, we sought to construct a measure representing the rate at which older GLs were displaced by more recent GLs (for those trees with at least 2 GLs identified). This is related to the rate at which clades go extinct. We considered the Shannon entropy of the clade distribution calculated over sliding windows containing the nearest 5% of all isolates (including redundant sequences and not limited to tree leaves) based on the known or estimated date of isolation $S_t^j$ OR distance to the tree root $S_d^j$ (see main text). Dates/root distances were sampled from that corresponding to the earliest or deepest isolate in any GL to the latest or shallowest respectively. The distance to the root was linearly scaled to represent the date of isolation corresponding to each sequence in an alternative genealogical tree where the date of isolation for each sequence exactly corresponds to the distance to the root for that sequence (mapped to the established date range).

The entropy is reported in units of base *N*, the number of GLs in the tree so that the maximum value is 1 and the minimum value is 0 for all trees. A mean entropy near 0 represents a genealogical tree composed of clades which quickly displace one another. A mean entropy near 1 represents a genealogical tree where all clades are equally distributed at every time point. We observe $\langle S_t^j \rangle_j > \langle S_d^j \rangle_j$ in all but one case (HDV) indicating that the entropy is larger than that expected from an analysis of the tree structure with no known dates of isolation (see Fig. 3D).

**Reconstruction of Ancestral Genome Sequences**

Fitch Traceback (11) was used to estimate ancestral states. Briefly, character sets were constructed moving from leaf to root. Each node was assigned the intersection of the descendant character sets if not empty and the union otherwise. Moving from root to leaf, nodes with more than one character in their set were assigned the consensus character if present in their set or a randomly chosen representative character otherwise. Substitutions between states were identified and placed in the middle of the branch bridging the pair of nodes.

**Mutational Signatures**

In an effort to distinguish intra-GL and inter-GL mutational signatures, three classes of amino acid sites were identified on the basis of the nonsynonymous mutations within each site. Redundant sequences were not evaluated in this analysis. 1) Multiple, deep (MD) substitutions are “lineage defining”, being conserved in at least 90% of the samples within at least two GLs, but represented by different amino acid residues in each of these GLs. For example, consider the third amino acid of the CHIKV ORF gp1. 97% of the sequences in GL 1 contain a Serine in that site while 96% of the sequences in GL2 contain a Proline in that site.2) Multiple, shallow sites. Consider a nonsynonymous mutation in site *M* which appears independently *N* times across all GLs combined (but may appear only once within any GL). If *N* is no less than the threshold defined to be the 90$^{th}$ percentile of the distribution over all *N* or 2 (whichever is larger), site *M* is included in this class of repeat, shallow sites. 3) All shallow sites bearing at least one nonsynonymous mutation which occurs within a GL. Only viruses with at least 2 GLs were included in this analysis. We then computed the site density of each class over a sliding window of 101 amino acid sites respecting mature peptide boundaries. The site densities across the genomes are displayed (normalized with respect to the mean site density for each class over the entire genome) in Supplemental Figures S18-24.

**Computation of *dN/dS***

For each mature protein sequence within each alignment, 10 reduced alignments were constructed as follows. Sequences were iteratively included in each reduced alignment to maximize diversity by maximizing the minimum hamming distance to the ensemble at each step. The first 10 sequences are conserved across every alignment and the remaining 40 are unique to each alignment. The reference sequence, was additionally added to each reduced alignment and PAML(12) was used to estimate tN, tS, *dN/dS*, N, S, and N/S for each segment and every reduced alignment. For sub-lineages (manually selected and GL) containing less than 410 sequences, the maximal number of reduced alignments was constructed. The reference sequence was not added to reduced alignments of sub-lineages.

Runs with total tree lengths above 50 (with Branch length defined as number of nucleotide substitutions per codon) were removed (resulting in fewer than 10 independent estimates for some viruses). These cases most likely represent ML estimates which did not converge. Median *dN/dS* is calculated for each peptide as well as mean *dN/dS* over all proteins (normalized to their length). Whenever saturation of *dS* (>1.5) is observed for any reduced alignment, the corresponding peptides are highlighted in the respective figures. No *dN/dS* is given for the following peptides: EVCr_sub3 2B and VPg, OHVA_sub1 A1pp, OHVA_sub3 Viral helicase and TBEV_sub2 ancC. In these cases, the sub lineages contained fewer than 90 samples resulting in only one reduced alignment showing an unreliable total branch length above 50 nucleotide substitutions per codon. Mean *dN/dS* ratios for all peptides of all lineages can be found in Supplementary Data, Table S9.

**Estimation of the Total and Effective Population Size**

The effective population size *Ne* and the ratio of the census population size *N* over *Ne* was estimated as previously described (13). In brief, genealogical trees were analyzed with the PACT package (http://www.trevorbedford.com/pact). Here, coalescence rate (Cr) is defined by the number of possible coalescent events (equal to the number of internal branches) divided by the opportunity for a coalescent event in a given fixed window of time.

The opportunity is defined as

$$opportunity = \sum_i \frac{kn(n-1)}{2}$$

With *k* being the length of the interval, *n* the number of concurrent lineages and *i* the discrete set of intervals. In our case, each tree is sliced into 1000 equal parts (*i*) from root to most distant tip with the tree specific length of the interval *k* in years. Given

$$Ne * t \sim 1/Cr$$

where *t* is the viral generation time, one can estimate *Ne* in generations which is equal to the number of individuals assuming the Fisher-Wright model (see, for example, (14)). Generation time *t* was estimated for each virus based on available epidemiological data. To retrieve *N/Ne* ratios, yearly cases were estimated for each virus and N was calculated as

$$N = D * t$$

with *D* being estimated yearly cases. *t* and *N* estimations can be found in Supplementary Data, Table S10, including references.

#### Estimation of diversity and TMRCA over time in skyline plots

Viral diversity (average time of any pair of leaves at a given timepoint to find their LCA) and average time to most recent common ancestor (TMRCA) over time were calculated with the PACT package(http://www.trevorbedford.com/pact). Within PACT, step size was set to 2 years to avoid windows with only one branch for which no opportunity could be calculated (see above). The analysis was repeated four times with a 0.25-year shift for the 2-year window to allow optimal resolution. At each timepoint, a cross-section of the tree is taken. Branches cut at this section are treated as terminal leaves. Then, diversity and mean TMRCA are calculated for this timepoint. Skyline plots for main virus populations, estimations from grafted trees, and all GLs can be found in the Supplementary Data.

#### Simulating trees with respect to selection strength and sampling density

In order to demonstrate the potential equivalency between the impacts of selection strength and sampling density on effective population size, we simulated an ensemble of trees as follows. Beginning with a rooted, bifurcated tree consisting of 2 leaves with uniformly distributed random branch lengths (between 0 and 1), the tree was extended by adding a multifurcated node at the “end” of a leaf so that the terminal branch length became the branch leading to the added multifurcation. The branch lengths of all leaves within the multifurcation were also random and uniformly distributed between 0 and 1. This process was continued for 500 iterations. At each iteration up to 10 leaves were selected for extension. To represent increased sampling, the multifurcation multiplicity was varied from 2 to 6. To represent increased selection strength, leaves were selected for extension only if they were positioned beyond a given threshold distance to the root of the tree. This threshold was set to be the 0th, 25th, 50th, 75th, or 95th percentile of the root distance distribution for the entire tree at that iteration. Note that in practice, multifurcations were represented as bifurcating subtrees with arbitrary topology and arbitrarily short interior branch lengths.

**Metadata**

Metadata including the sequence ID, host, date of isolation, location of isolation, and subtype for all sequences is available in Supplementary Data, Table S11.

#### List of abbreviations

**AS:** All, shallow

**BCoV1:** Betacoronavirus 1

**CHIKV**: Chikungunya virus

**DENV:** Dengue virus

**dN/dS:** ratio of non-synonymous to synonymous substitution rates

**DUF:** Domain of unknown function

**Ebola:** Zaire ebolavirus

**EVA:** Enterovirus A

**EVAd:** Enterovirus A 'diverse'

**EVAe:** Enterovirus A 'even'

**EVB:** Enterovirus B

**EVC:** Enterovirus C

**EVCe:** Enterovirus C 'every' sample

**EVCr:** Enterovirus C 'reduced' samples

**EVD:** Enterovirus D

**GL:** Genealogical lineage

**H3N2d:** H3N2 'divers'

**H3N2e:** H3N2 'even'

**HBV:** Hepatitis B virus

**HCV:** Hepatitis C virus

**HCVe:** Hepatitis C virus 'every' sample

**HCVr:** Hepatitis C virus 'reduced' samples

**HDV:** Hepatitis D virus

**HMPV:** Human metapneumovirus

**HRSV:** Human respiratory syncytial virus

**HRV3:** Human respirovirus 3

**H3N2:** Influenza A Virus H3N2

**LCA:** Last common ancestor

**LHDAg:** Large human delta antigen

**LSD:** least-square distance

**MD:** Multiple, deep

**MERS:** Middle East respiratory syndrome-related coronavirus

**ML:** Manual lineage

**MMV:** Measles morbillivirus

**MRCA:** Most recent common ancestor

**MRV:** Mumps rubulavirus

**MS:** Multiple, shallow

**N:** Census population size

**Ne:** Effective population size

**Norwalk:** Norwalk virus

**NS:** Non-structural protein

**NSP:** Non-structural protein

**OHVA:** Orthohepevirus A

**PeVA:** Parechovirus A

**RVA:** Rhinovirus A

**SARS-CoV-2:** Severe acute respiratory syndrome-related coronavirus 2

**SV:** Sapporo virus

**TMRCA:** Time to most recent common ancestor

**YFV:** Yellow fever virus

**ZIKV:** Zika virus

### Declarations

#### Ethics approval and consent to participate

Not applicable.

#### Consent for publication

Not applicable.

#### Data Availability

The datasets generated and/or analyzed during the current study are available as supplementary data at Zenodo, https://doi.org/10.5281/zenodo.5711959, as well as through FTP, https://ftp.ncbi.nih.gov/pub/wolf/_suppl/virNiches/. Original virus sequences are publicly available for all viruses except SARS-CoV-2 at NCBI virus (1). SARS-CoV-2 sequences are available at GISAID (3).

#### Competing interests

The authors declare that they have no competing interests.

#### Funding

NDR, YIW PM, and EVK are supported by the Intramural Research Program of the National Institutes of Health (National Library of Medicine).

#### Authors' contributions

PM, NDR, and GF collected data; PM, NDR, YIW, GF, FZ, and EVK analyzed data; PM, NDR, and EVK wrote the manuscript that was edited and approved by all authors.

#### Acknowledgements

The authors thank Koonin group members for helpful discussions.

### References


1. Hatcher EL*, et al.* (2017) Virus Variation Resource - improved response to emergent viral outbreaks. *Nucleic Acids Res* 45(D1):D482-d490.
2. Bao Y*, et al.* (2008) The influenza virus resource at the National Center for Biotechnology Information. *J Virol* 82(2):596-601.
3. Shu Y & McCauley J (2017) GISAID: Global initiative on sharing all influenza data–from vision to reality. *Eurosurveillance* 22(13):30494.
4. Rochman ND*, et al.* (2021) Ongoing global and regional adaptive evolution of SARS-CoV-2. *Proc Natl Acad Sci U S A* 118(29).
5. Katoh K, Misawa K, Kuma Ki, & Miyata T (2002) MAFFT: a novel method for rapid multiple sequence alignment based on fast Fourier transform. *Nucleic acids research* 30(14):3059-3066.
6. Li W & Godzik A (2006) Cd-hit: a fast program for clustering and comparing large sets of protein or nucleotide sequences. *Bioinformatics* 22(13):1658-1659.
7. simplemaps (2021) World Cities Database.
8. Nguyen L-T, Schmidt HA, Von Haeseler A, & Minh BQ (2015) IQ-TREE: a fast and effective stochastic algorithm for estimating maximum-likelihood phylogenies. *Molecular biology and evolution* 32(1):268-274.
9. Price MN, Dehal PS, & Arkin AP (2010) FastTree 2–approximately maximum-likelihood trees for large alignments. *PloS one* 5(3):e9490.
10. To T-H, Jung M, Lycett S, & Gascuel O (2016) Fast dating using least-squares criteria and algorithms. *Systematic biology* 65(1):82-97.
11. Fitch WM (1971) Toward defining the course of evolution: minimum change for a specific tree topology. *Systematic Biology* 20(4):406-416.
12. Yang Z (2007) PAML 4: phylogenetic analysis by maximum likelihood. *Molecular biology and evolution* 24(8):1586-1591.
13. Bedford T, Cobey S, & Pascual M (2011) Strength and tempo of selection revealed in viral gene genealogies. *BMC Evol Biol* 11:220.
14. Ishida Y & Rosales A (2020) The origins of the stochastic theory of population genetics: The Wright-Fisher model. *Stud Hist Philos Biol Biomed Sci* 79:101226.

#### Supplemental Figure Legends

**Figure S1.** The fraction of sequences included in any correlated-clade (genealogical lineage) as a function of the Pearson correlation coefficient threshold. Left. All sequences. Right. Sequences with metadata.

**Figure S2-7.** The distance to the tree root vs. the date of isolation. Each point represents a leaf in the tree. Filled circles represent leaves included in a correlated-clade and are colored accordingly. Open circles represent leaves which are not included in any correlate clade. For SARS-CoV-2 and H3N2 the entire tree was considered to constitute a single, correlated-clade.

**Figure S8.** The ratio of the mean inter- vs. intra-lineage map distance for all pairs of lineages for manually selected lineages on top and genealogical lineages (GL) below. The mean inter-lineage map distance is the mean great circle distance between sequencing locations taken over all pairs of isolates spanning the pair of lineages. The mean intra-lineage map distance is taken over all pairs of isolates within each lineage and averaged over the two mean values for each lineage. Filled circles represent pairs of lineages where the colors represents the number of isolates within the pair. Open circles represent the mean ratio weighted by the number of isolates in each pair.

**Figure S9.** *dN/dS* ratios were analysed for each protein of each virus and are represented by a single dot. Black line represents mean over all proteins (normalized to length). Black dots represent proteins with non-reliable *dN/dS* due to saturation of synonymous substitutions.

**Figure S10-13.** *dN/dS* ratios retrieved from trees covering the whole population (corresponding to Figure S9) and trees representing genealogical lineages (GL). Each protein is represented by a single dot, black line represents mean over all proteins. Black coloured dots represent proteins with non-reliable *dN/dS* due to saturation of synonymous substitutions.

**Figure S14-17.** *dN/dS* ratios retrieved from trees covering the whole population (corresponding to Figure S9) and trees representing manually selected lineages (correlating to viral sero- or genotypes). Each protein is represented by a single dot, black line represents mean over all proteins. Black coloured dots represent proteins with non-reliable *dN/dS* due to saturation of synonymous substitutions.

**Figure S18-24.** The site density of three classes of mutations – Multiple, deep (MD, red), multiple, shallow (MS, yellow), and all shallow (AS, blue, see Methods and Figure 2) – computed over a sliding window of 101 amino acid sites respecting mature peptide boundaries and normalized with respect to the mean site density for each class over the entire genome.

**Figure S25.** Mutation rates (as substitutions per site per year) and estimated date of the last common ancestor for **A.** both HCV and EVC trees considered (e/r with and without manual pruning, see Methods). **B.** H3N2 diverse and evenly sampled subtrees (d/e) as well as new/old subtrees (n/o). **C.** EVA diverse and evenly sampled subtrees (d/e). Whole trees are shown green, subtrees in red.

**Figure S26. A.** Ne replicates for simulated trees, depending on sampling multiplicity and root distance threshold (representing selection). Largest repetition in red, middle in black and smallest in yellow. **B.** Sample multiplicity for each replicate of simulated trees plotted against *Ne* and number of leaves in the final tree. Sampling multiplicities shown in dark blue, light blue, green, brown, and yellow for multiplicities of 2, 3, 4, 5, and 6 respectively. **C.** Root distance threshold plotted against *Ne* and number of leaves of final simulated trees. Root distance thresholds shown in dark blue, light blue, green, brown, and yellow for thresholds of 0, 25, 50, 75 and 95, respectively. **D.** Increase of Ne with increased sampling density relative to selection strength (root distance thresholds 0, 25, 50, 75 and 95).

**Figure S27. A**. Schematic of genealogical tree construction. Phylogenetic tree (1) which is either directly converted into genealogical tree (2) or split into clades (3) (as described in Fig. 2). Genealogical trees for each clade are calculated (4) and clades grafted at the root of oldest clade, keeping date of each leave intact (5). **B.** *Ne* estimates (according to Fisher-Wright model) for whole trees, clades and grafted trees. **C.** Estimated generation time *t* in days. Bars represent 0.5-5 times the value of the filled circle. **D.** Estimation of yearly viral cases. Bars represent 0.5-5 times the value of the filled circle.

**Figure S28. A**. Mean diversity per year based on genealogical trees for the main population, genealogical lineages (GL) and grafted trees. Diversity is calculated with PACT (the average time in years for any 2 pairs at a given timepoint to coalesce). **B-D**. Representative skyline plots for diversity and time to most recent common ancestor (TMRCA) of selected viral main populations or genealogical lineages. Plots for all sets can be found in the Supplementary data.

**Figure S29.** The date of isolation for redundant isolates vs. the date of isolation for the leaf in the tree to which these sequences are mapped for all virus species where more than 10% of sequences are redundant. Isolates and leaves with no known date of isolation are excluded. Number next to virus species gives fraction of samples with redundancy.

#### Supplementary Data

#### Tables

**Table S1:** Virus characteristics including abbreviation, family, order, tax id, sequence length threshold (for nearly complete genomes), vaccination status, mode of transmission, circulation (human or human/zoonotic), available treatment, and disease progression (acute or chronic).

**Table S2:** Sequence IDs of outgroup constituents.

**Table S3:** GISAID acknowledgements for the SARS-CoV-2 sequences used in this study.

**Table S4:** Summary of manual sequence curation indicating lab- or vaccine-related keywords used to prune sequences.

**Table S5:** Sequence IDs for EVA and H3N2 e10, e100, d10, and d100 subtrees.

**Table S6:** Sequence IDs for ML and GL lineages.

**Table S7:** Sequence IDs for the H3N2 “new” and “old” subtrees.

**Table S8:** Mutation rates (in units of nucleotide substitutions per site per year) and estimated dates for the last common ancestor of all tree/subtrees. Negative dates represent years prior to 0 CE.

**Table S9:** Mean dN/dS values.

**Table S10:** Yearly viral cases and generation time estimations (with references).

**Table S11:** Metadata including the sequence ID, host, date of isolation, location of isolation, and subtype.

#### Directories

**Alignments** ORFeome alignments (excluding stop codons) with the exception of SARS-CoV-2.

**Correlated Subtrees** Subtrees representing all correlated-clades (genealogical lineages; GL) in Newick format.

**Diversity TMRCA and Skyline Plots** .png files.

**e/d/n/o Subtrees** EVA and H3N2 subtrees, evenly and diversely sampled as well as "early" and "late" subtrees for H3N2.

**Genealogical Trees** Including global topologies (.main), subtrees, and grafted trees (.grafted).

**ORF References** The first and last nucleotide of each ORF in the reference sequence.

**Redundancy Tables** List of all redundant isolates and the corresponding representative in the ORFeome-unique alignment.

**Reference Sequences** Genbank page for all reference sequences.

**Rooted Trees** Global topologies for each virus.

**Ultrametric Trees** For each global topology.

**S1**

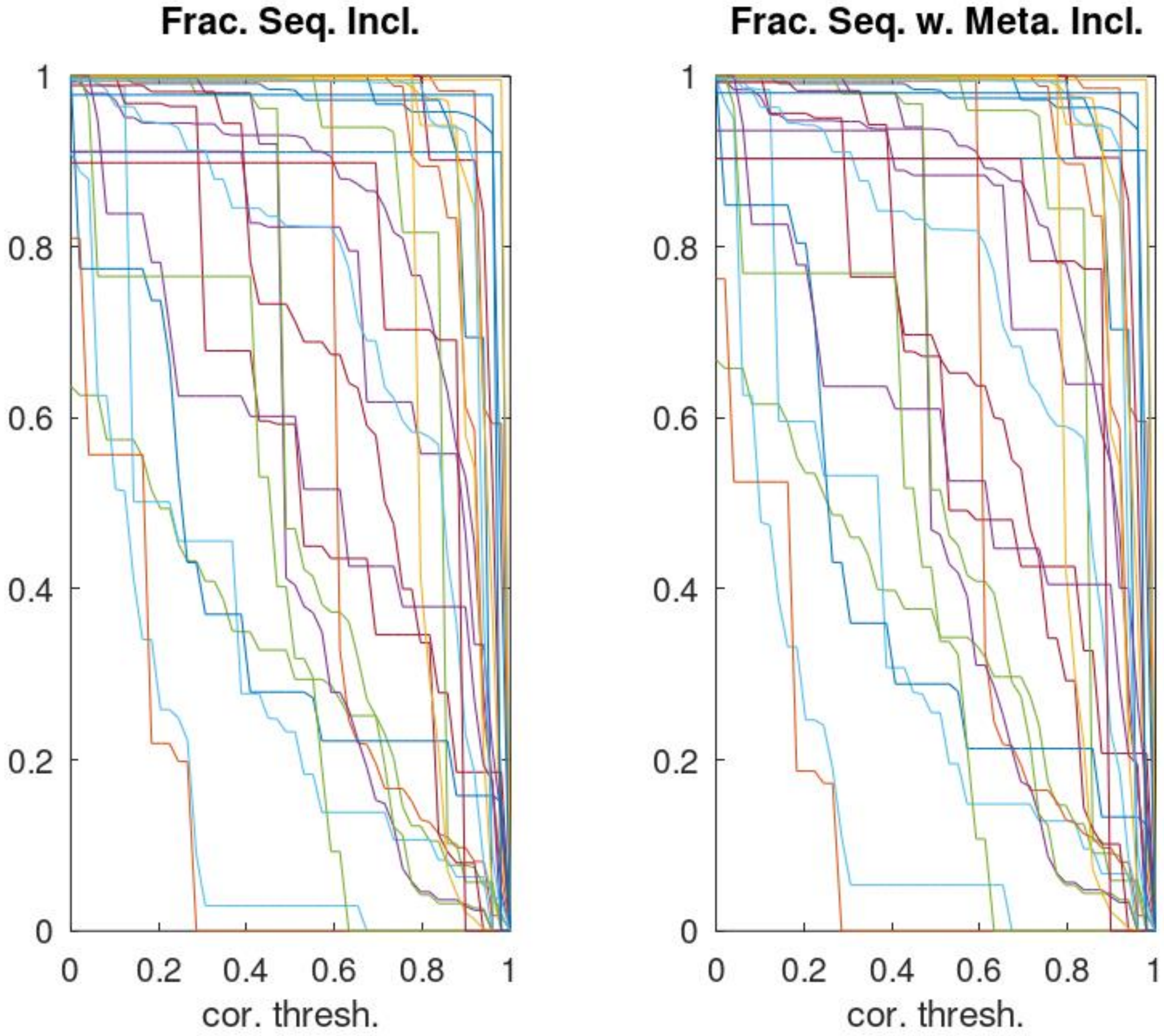
Frac. Seq. Incl.
1
0.8
0.6
0.4
0.2
0
0
0.2
0.4
0.6
0.8
1
cor. thresh.
Frac. Seq. w. Meta. Incl.
1
0.8
0.6
0.4
0.2
0
0
0.2
0.4
0.6
0.8
1
cor. thresh.

S2

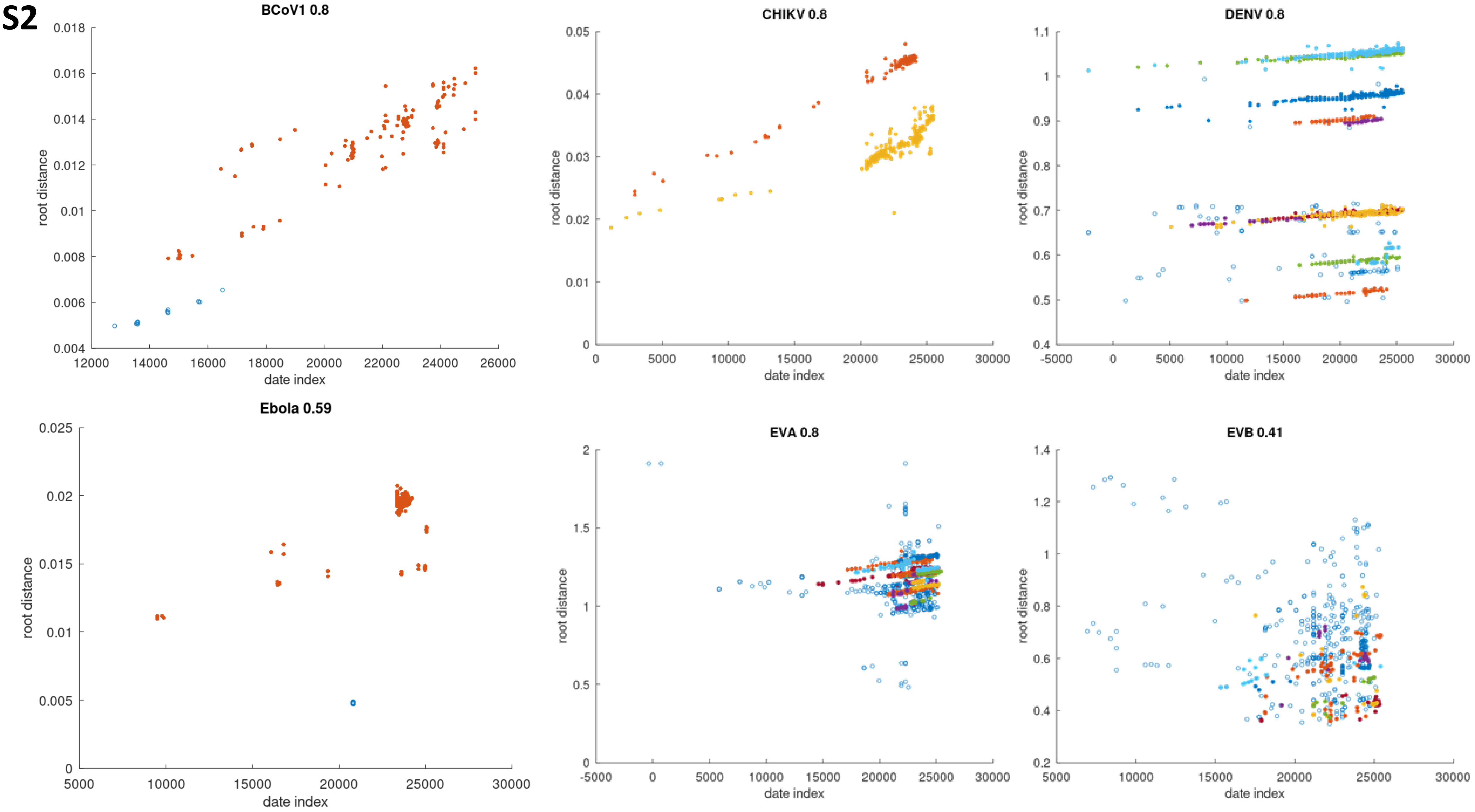
BCoV1 0.8
CHIKV 0.8
DENV 0.8
Ebola 0.59
EVA 0.8
EVB 0.41
root distance
date index

S3

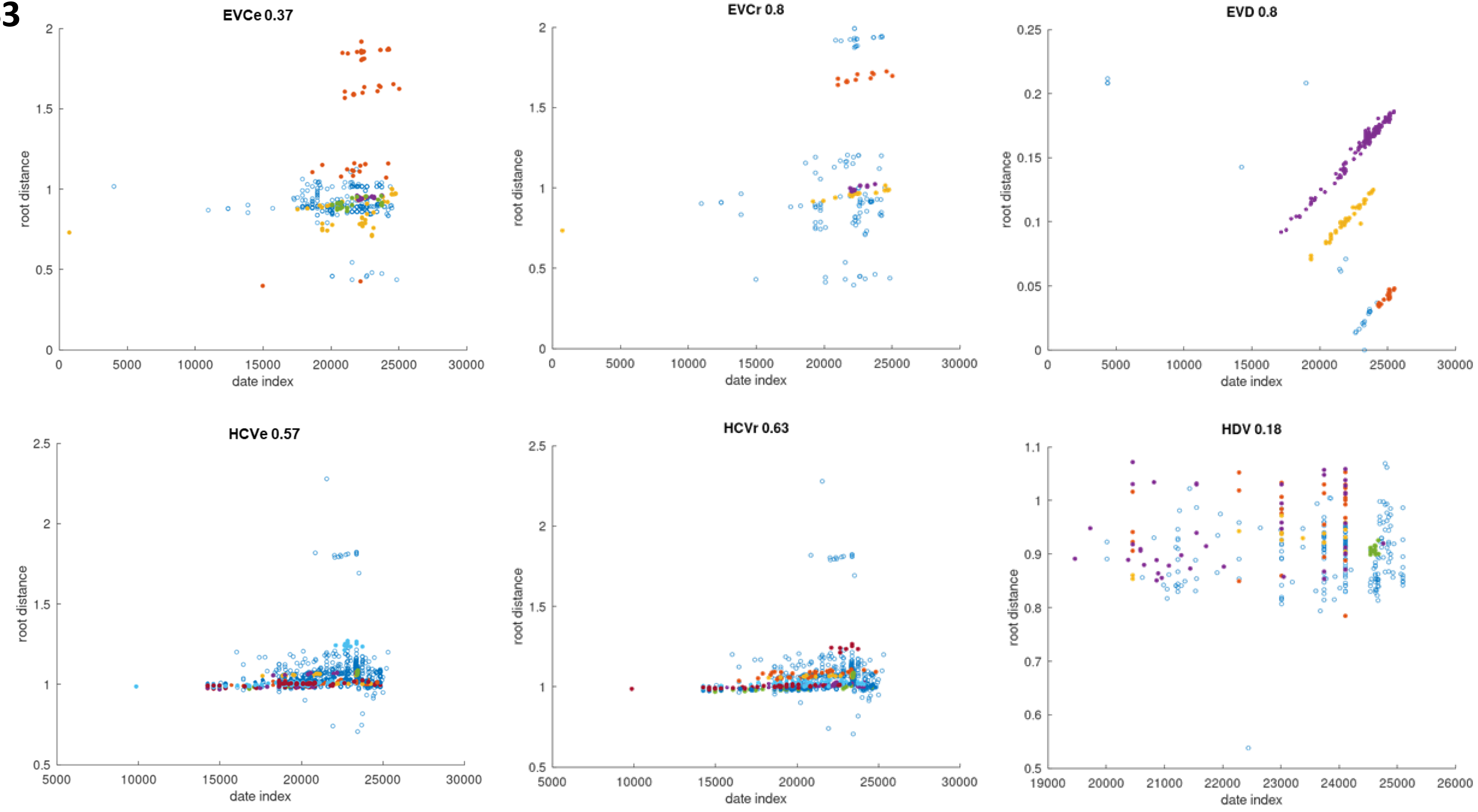
EVCe 0.37
root distance
date index
EVCr 0.8
root distance
date index
EVD 0.8
root distance
date index
HCVe 0.57
root distance
date index
HCVr 0.63
root distance
date index
HDV 0.18
root distance
date index

**S4**

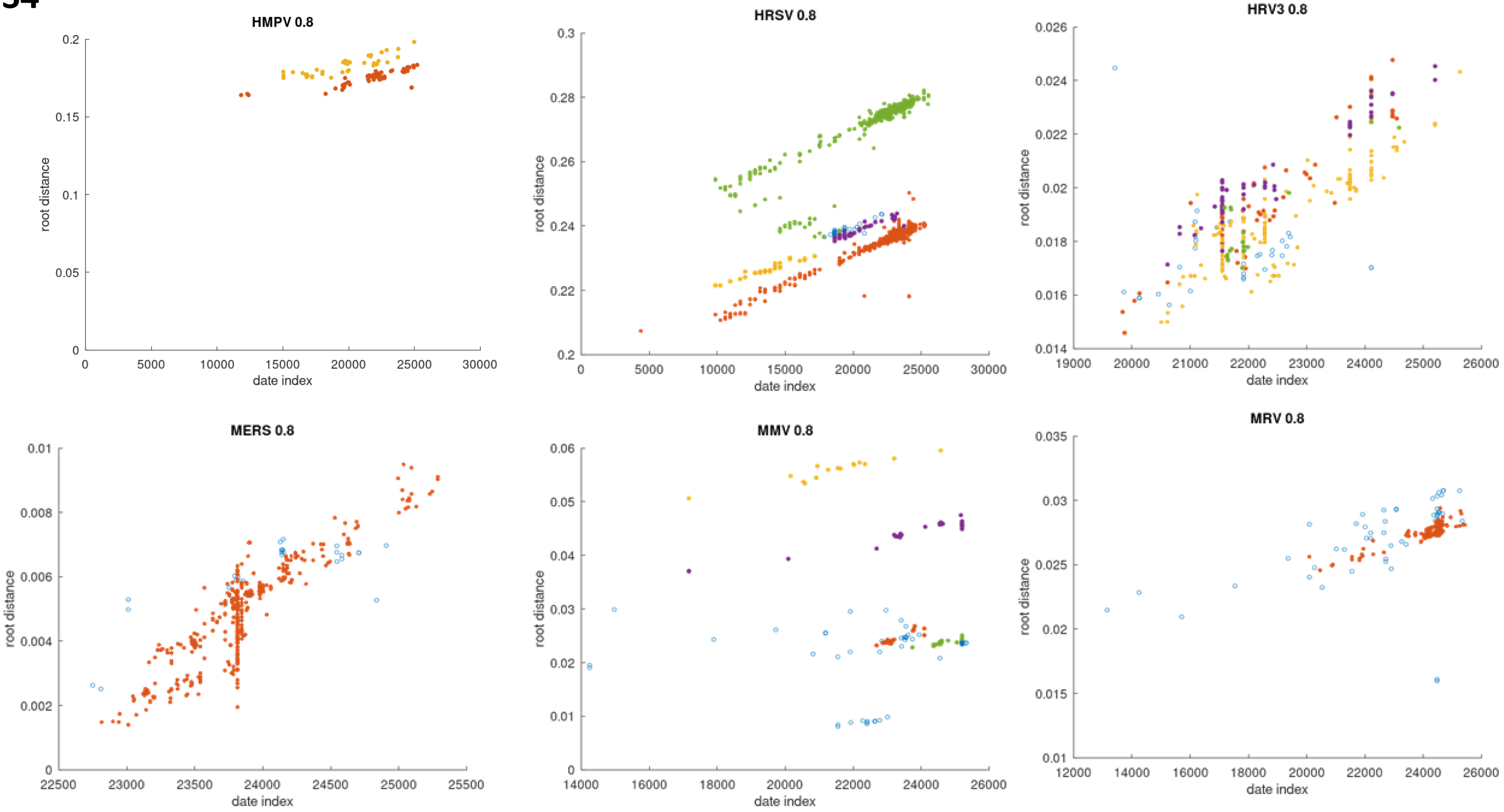

HMPV 0.8
HRSV 0.8
HRV3 0.8
MERS 0.8
MMV 0.8
MRV 0.8
root distance
date index

**S5**

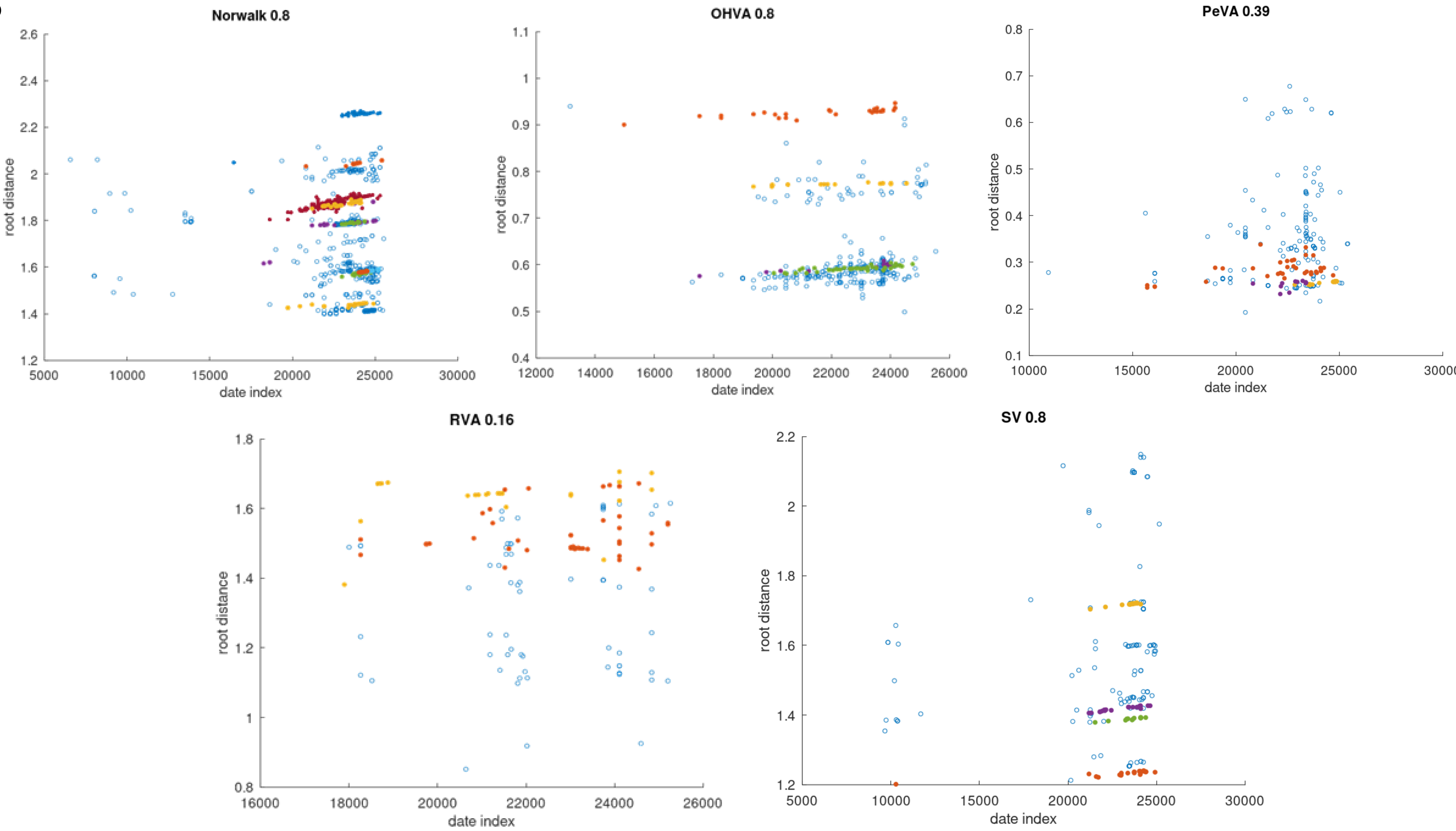
Norwalk 0.8
OHVA 0.8
PeVA 0.39
RVA 0.16
SV 0.8
root distance
date index

S6

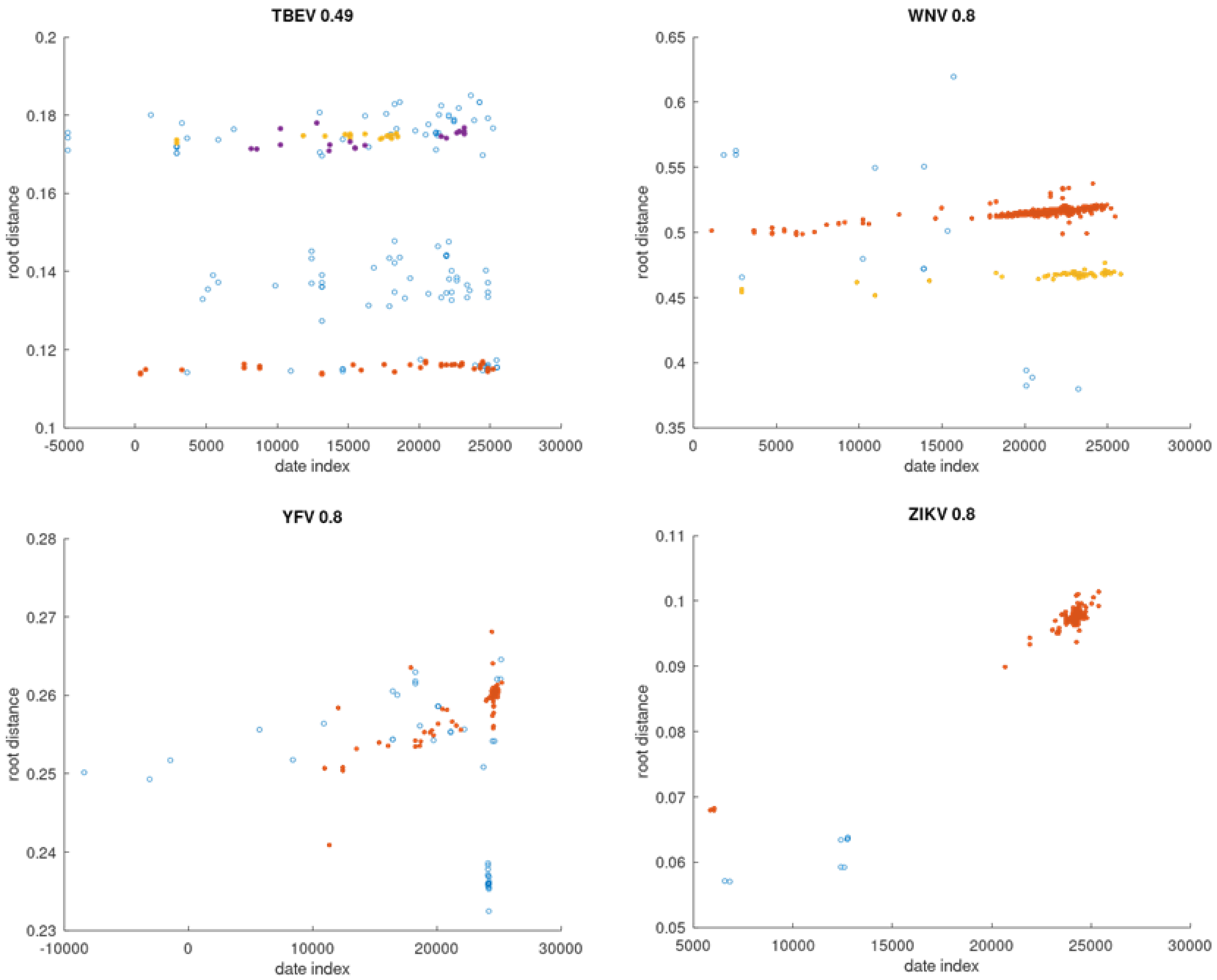

**S7**

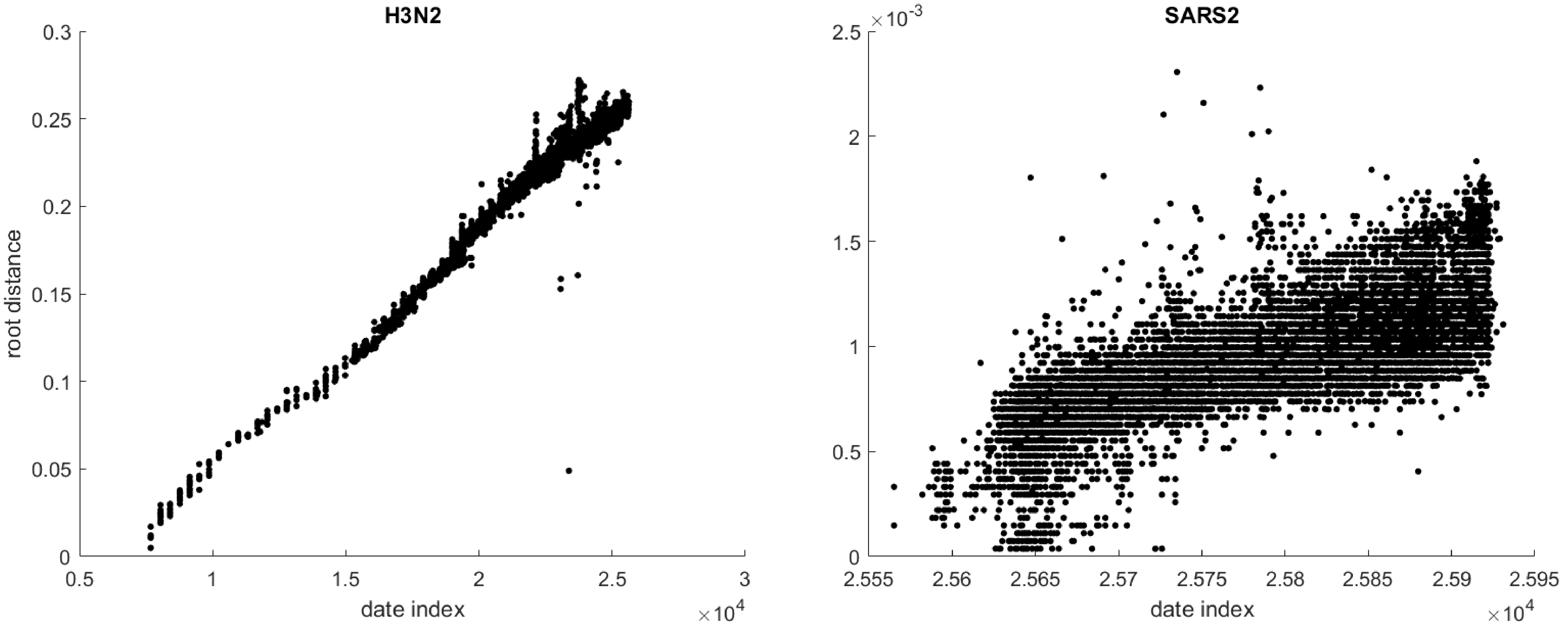
H3N2
root distance
date index
×10^4
0.3
0.25
0.2
0.15
0.1
0.05
0
0.5
1
1.5
2
2.5
3
SARS2
×10^-3
2.5
2
1.5
1
0.5
0
2.555
2.56
2.565
2.57
2.575
2.58
2.585
2.59
2.595
date index
×10^4

S8

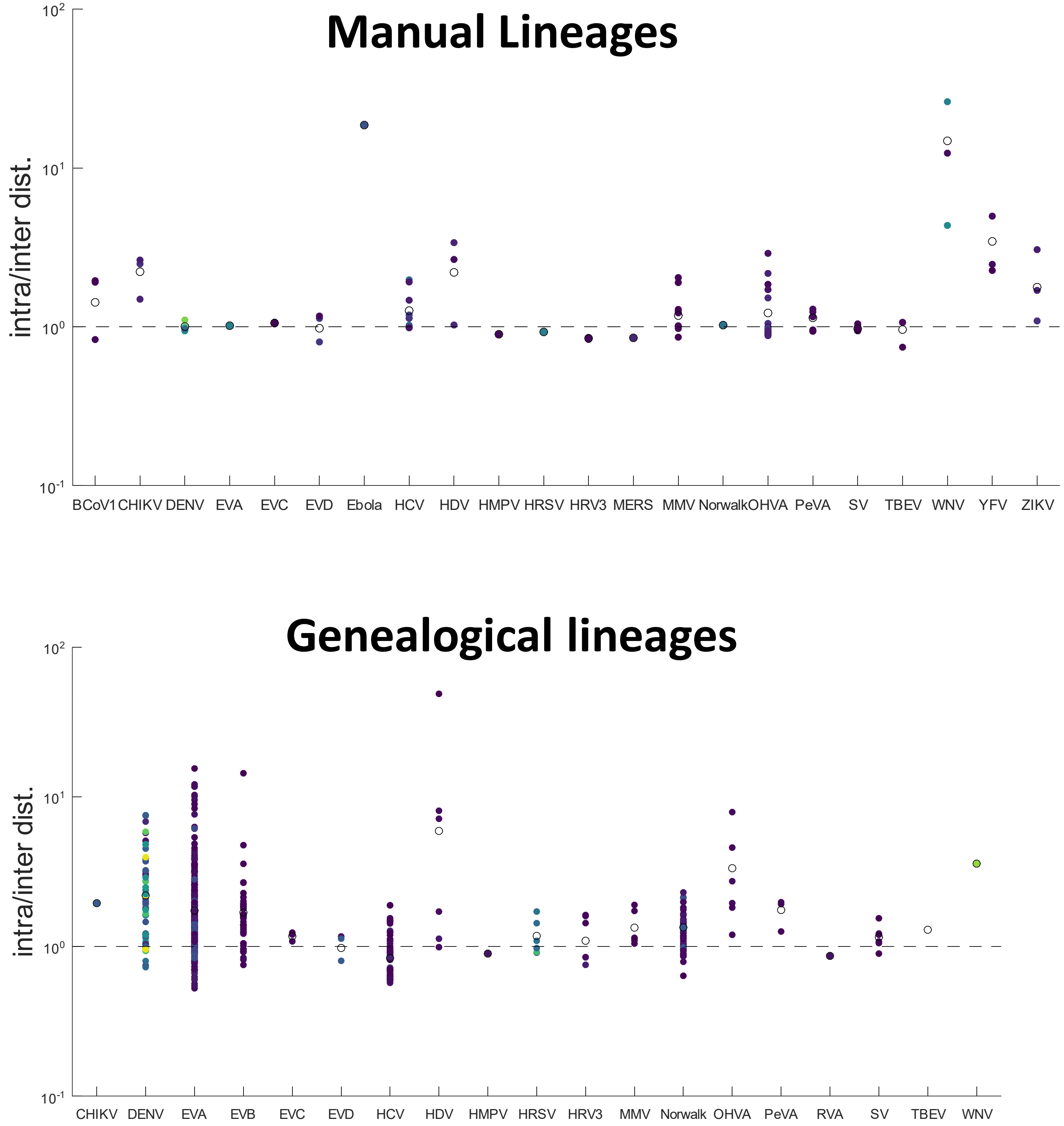
Manual Lineages
intra/inter dist.
$10^2$
$10^1$
$10^0$
$10^{-1}$
BCoV1 CHIKV DENV EVA EVC EVD Ebola HCV HDV HMPV HRSV HRV3 MERS MMV Norwalk OHVA PeVA SV TBEV WNV YFV ZIKV
Genealogical lineages
intra/inter dist.
$10^2$
$10^1$
$10^0$
$10^{-1}$
CHIKV DENV EVA EVB EVC EVD HCV HDV HMPV HRSV HRV3 MMV Norwalk OHVA PeVA RVA SV TBEV WNV

**S9**

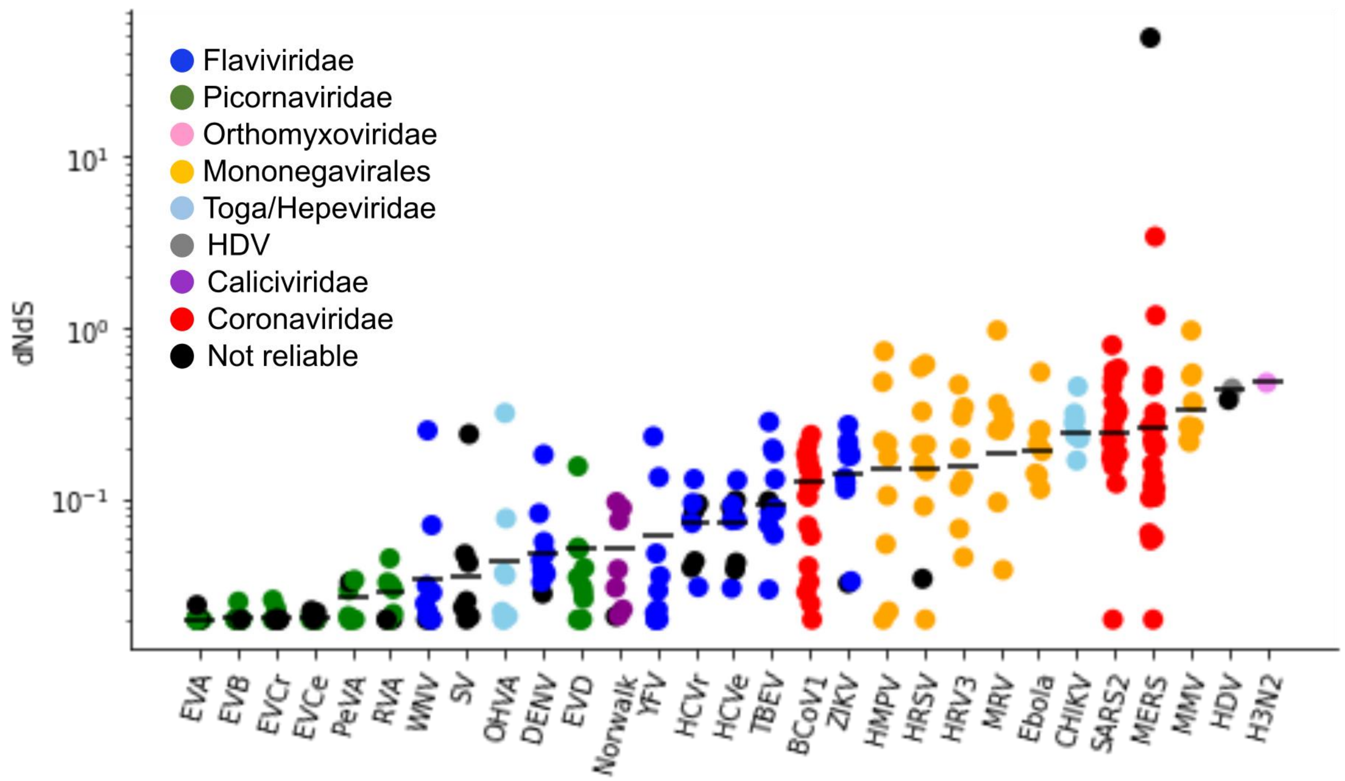

Flaviviridae
Picornaviridae
Orthomyxoviridae
Mononegavirales
Toga/Hepeviridae
HDV
Caliciviridae
Coronaviridae
Not reliable
dNdS
$10^1$
$10^0$
$10^{-1}$
EVA
EVB
EVCr
EVCe
PeVA
RVA
WNV
SV
OHVA
DENV
EVD
Norwalk
YFV
HCVr
HCVe
TBEV
BCoV1
ZIKV
HMPV
HRSV
HRV3
MRV
Ebola
CHIKV
SARS2
MERS
MMV
HDV
H3N2

# S10

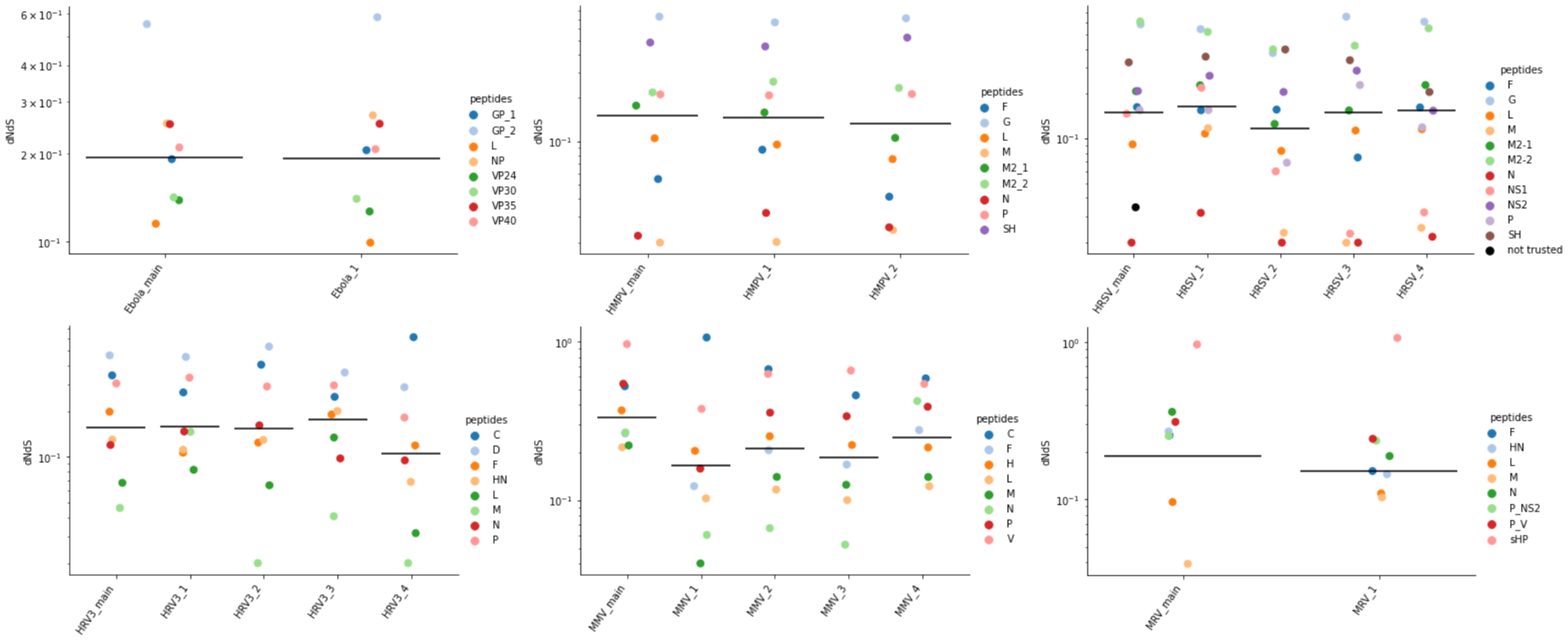

dNdS
Ebola_main
Ebola_1
peptides
GP_1
GP_2
L
NP
VP24
VP30
VP35
VP40
HMPV_main
HMPV_1
HMPV_2
F
G
M
M2_1
M2_2
N
P
SH
HRSV_main
HRSV_1
HRSV_2
HRSV_3
HRSV_4
M2-1
M2-2
NS1
NS2
not trusted
HRV3_main
HRV3_1
HRV3_2
HRV3_3
HRV3_4
C
D
HN
MMV_main
MMV_1
MMV_2
MMV_3
MMV_4
H
V
MRV_main
MRV_1
P_NS2
P_V
sHP

# S11

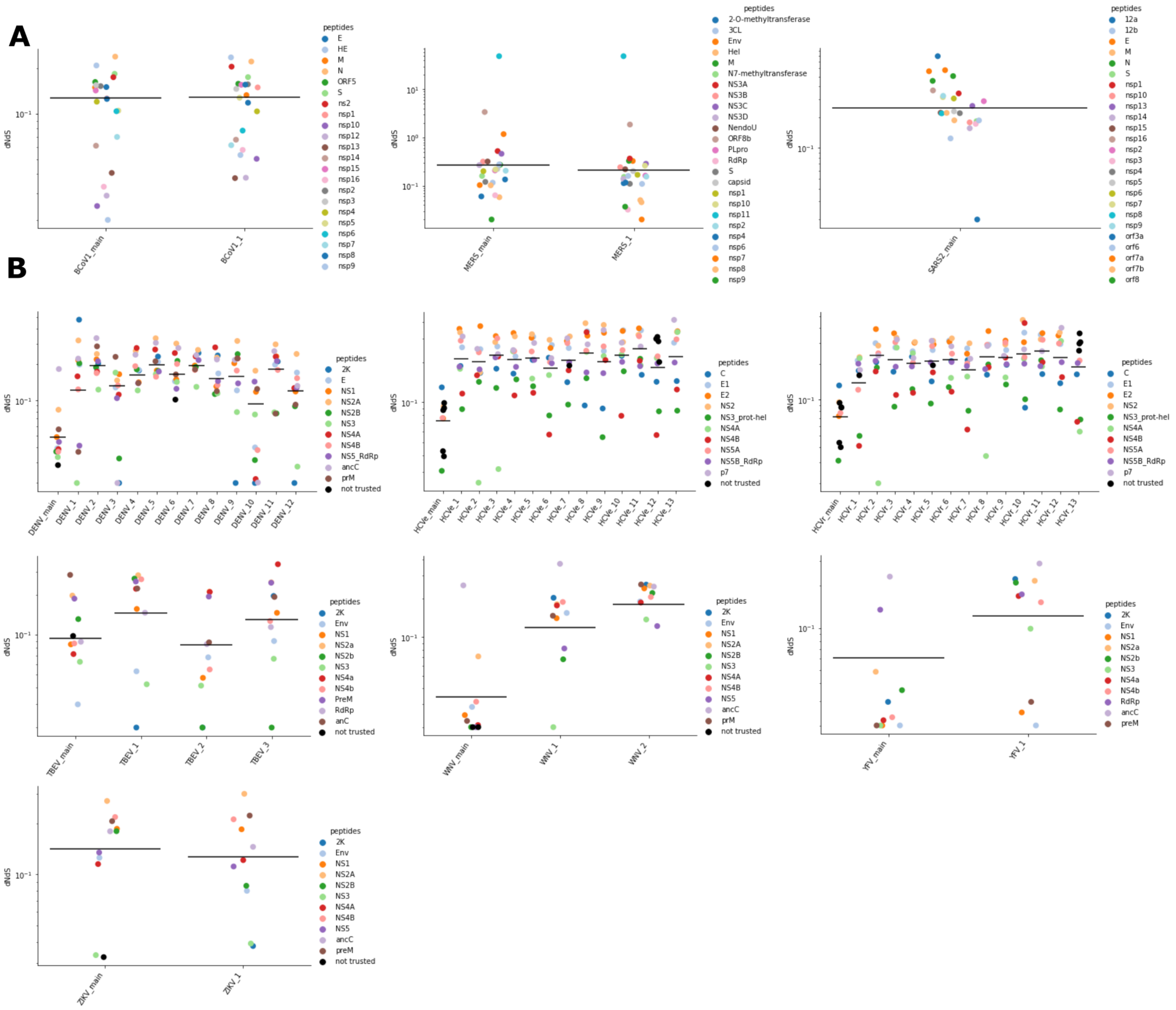
A
dNdS
BCoV1_main
BCoV1_1
peptides
E
HE
M
N
ORF5
S
ns2
nsp1
nsp10
nsp12
nsp13
nsp14
nsp15
nsp16
nsp2
nsp3
nsp4
nsp5
nsp6
nsp7
nsp8
nsp9
MERS_main
MERS_1
peptides
2-O-methyltransferase
3CL
Env
Hel
M
N7-methyltransferase
NS3A
NS3B
NS3C
NS3D
NendoU
ORF8b
PLpro
RdRp
S
capsid
nsp1
nsp10
nsp11
nsp2
nsp4
nsp6
nsp7
nsp8
nsp9
SARS2_main
peptides
12a
12b
E
M
N
S
nsp1
nsp10
nsp13
nsp14
nsp15
nsp16
nsp2
nsp3
nsp4
nsp5
nsp6
nsp7
nsp8
nsp9
orf3a
orf6
orf7a
orf7b
orf8
B
DENV_main
DENV_1
DENV_2
DENV_3
DENV_4
DENV_5
DENV_6
DENV_7
DENV_8
DENV_9
DENV_10
DENV_11
DENV_12
peptides
2K
E
NS1
NS2A
NS2B
NS3
NS4A
NS4B
NS5_RdRp
ancC
prM
not trusted
HCVe_main
HCVe_1
HCVe_2
HCVe_3
HCVe_4
HCVe_5
HCVe_6
HCVe_7
HCVe_8
HCVe_9
HCVe_10
HCVe_11
HCVe_12
HCVe_13
peptides
C
E1
E2
NS2
NS3_prot-hel
NS4A
NS4B
NS5A
NS5B_RdRp
p7
not trusted
HCVr_main
HCVr_1
HCVr_2
HCVr_3
HCVr_4
HCVr_5
HCVr_6
HCVr_7
HCVr_8
HCVr_9
HCVr_10
HCVr_11
HCVr_12
HCVr_13
TBEV_main
TBEV_1
TBEV_2
TBEV_3
peptides
2K
Env
NS1
NS2a
NS2b
NS3
NS4a
NS4b
PreM
RdRp
anC
not trusted
WNV_main
WNV_1
WNV_2
peptides
2K
Env
NS1
NS2A
NS2B
NS3
NS4A
NS4B
NS5
ancC
prM
not trusted
YFV_main
YFV_1
peptides
2K
Env
NS1
NS2a
NS2b
NS3
NS4a
NS4b
RdRp
ancC
preM
ZIKV_main
ZIKV_1
peptides
2K
Env
NS1
NS2A
NS2B
NS3
NS4A
NS4B
NS5
ancC
preM
not trusted

# S12

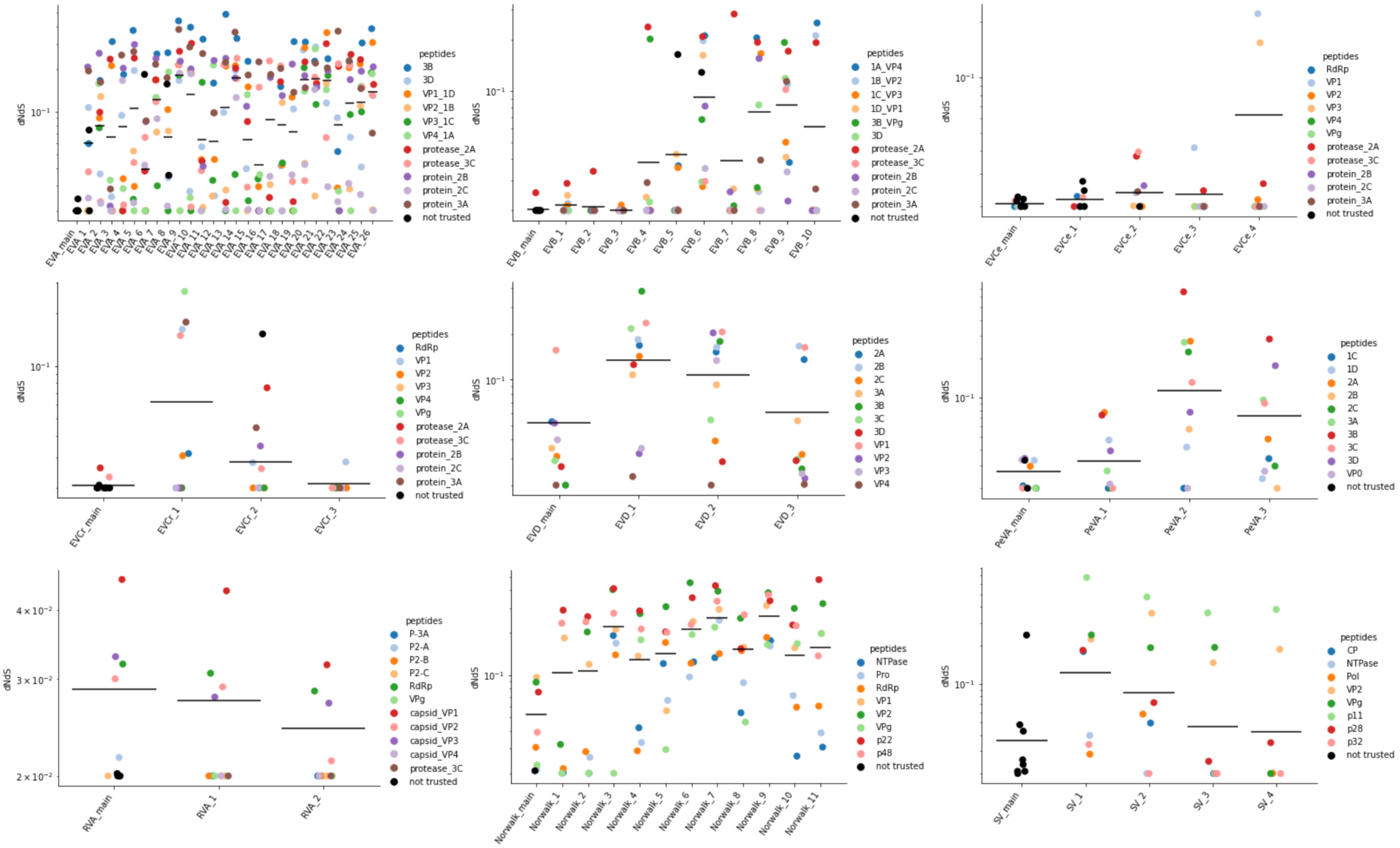

dNdS
$10^{-1}$
EVA_main EVA_1 EVA_2 EVA_3 EVA_4 EVA_5 EVA_6 EVA_7 EVA_8 EVA_9 EVA_10 EVA_11 EVA_12 EVA_13 EVA_14 EVA_15 EVA_16 EVA_17 EVA_18 EVA_19 EVA_20 EVA_21 EVA_22 EVA_23 EVA_24 EVA_25 EVA_26
peptides
3B
3D
VP1_1D
VP2_1B
VP3_1C
VP4_1A
protease_2A
protease_3C
protein_2B
protein_2C
protein_3A
not trusted
dNdS
$10^{-1}$
EVB_main EVB_1 EVB_2 EVB_3 EVB_4 EVB_5 EVB_6 EVB_7 EVB_8 EVB_9 EVB_10
peptides
1A_VP4
1B_VP2
1C_VP3
1D_VP1
3B_VPg
3D
protease_2A
protease_3C
protein_2B
protein_2C
protein_3A
not trusted
dNdS
$10^{-1}$
EVCe_main EVCe_1 EVCe_2 EVCe_3 EVCe_4
peptides
RdRp
VP1
VP2
VP3
VP4
VPg
protease_2A
protease_3C
protein_2B
protein_2C
protein_3A
not trusted
dNdS
$10^{-1}$
EVCr_main EVCr_1 EVCr_2 EVCr_3
peptides
RdRp
VP1
VP2
VP3
VP4
VPg
protease_2A
protease_3C
protein_2B
protein_2C
protein_3A
not trusted
dNdS
$10^{-1}$
EVD_main EVD_1 EVD_2 EVD_3
peptides
2A
2B
2C
3A
3B
3C
3D
VP1
VP2
VP3
VP4
dNdS
$10^{-1}$
PeVA_main PeVA_1 PeVA_2 PeVA_3
peptides
1C
1D
2A
2B
2C
3A
3B
3C
3D
VP0
not trusted
dNdS
$4\times10^{-2}$
$3\times10^{-2}$
$2\times10^{-2}$
RVA_main RVA_1 RVA_2
peptides
P-3A
P2-A
P2-B
P2-C
RdRp
VPg
capsid_VP1
capsid_VP2
capsid_VP3
capsid_VP4
protease_3C
not trusted
dNdS
$10^{-1}$
Norwalk_main Norwalk_1 Norwalk_2 Norwalk_3 Norwalk_4 Norwalk_5 Norwalk_6 Norwalk_7 Norwalk_8 Norwalk_9 Norwalk_10 Norwalk_11
peptides
NTPase
Pro
RdRp
VP1
VP2
VPg
p22
p48
not trusted
dNdS
$10^{-1}$
SV_main SV_1 SV_2 SV_3 SV_4
peptides
CP
NTPase
Pol
VP2
VPg
p11
p28
p32
not trusted

# S13

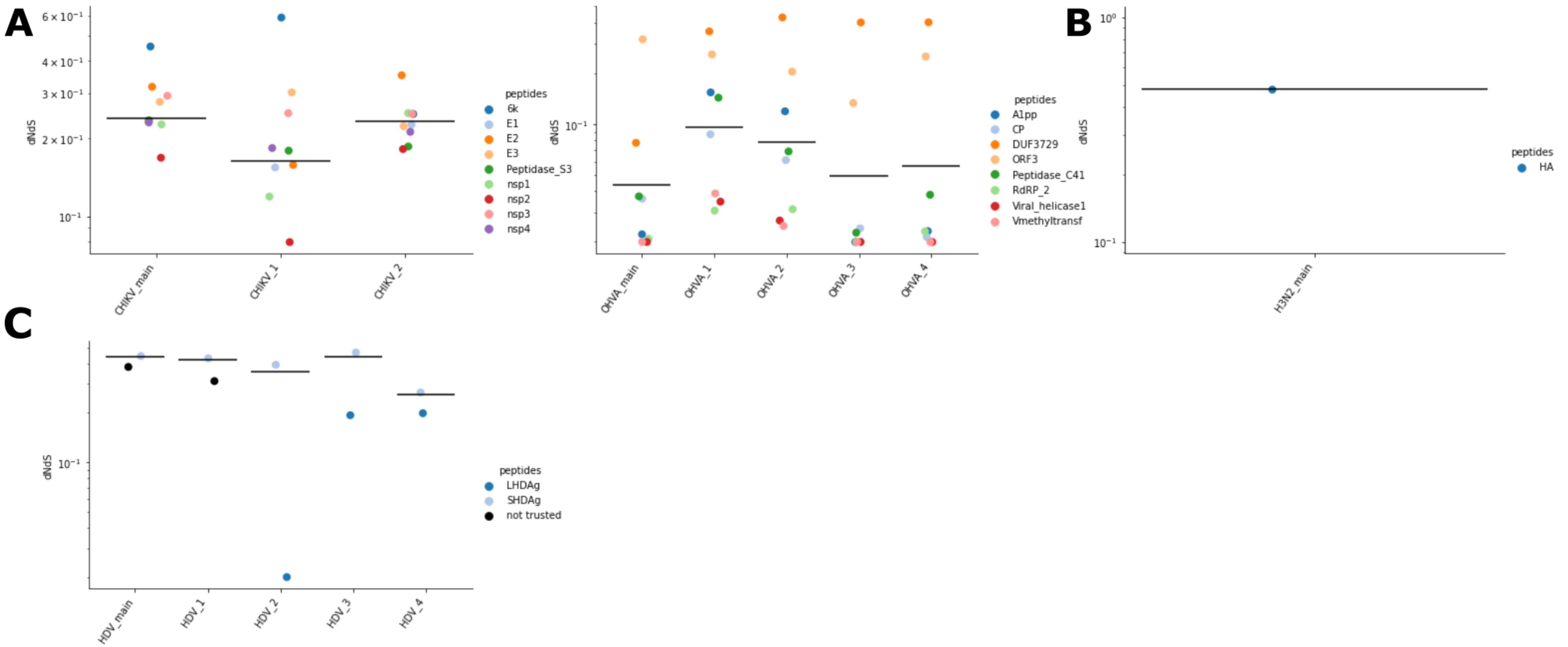
A
B
C
dNdS
$6\times10^{-1}$
$4\times10^{-1}$
$3\times10^{-1}$
$2\times10^{-1}$
$10^{-1}$
$10^{0}$
CHIKV_main
CHIKV_1
CHIKV_2
peptides
6k
E1
E2
E3
Peptidase_S3
nsp1
nsp2
nsp3
nsp4
OHVA_main
OHVA_1
OHVA_2
OHVA_3
OHVA_4
A1pp
CP
DUF3729
ORF3
Peptidase_C41
RdRP_2
Viral_helicase1
Vmethyltransf
H3N2_main
HA
HDV_main
HDV_1
HDV_2
HDV_3
HDV_4
LHDAg
SHDAg
not trusted

# S14

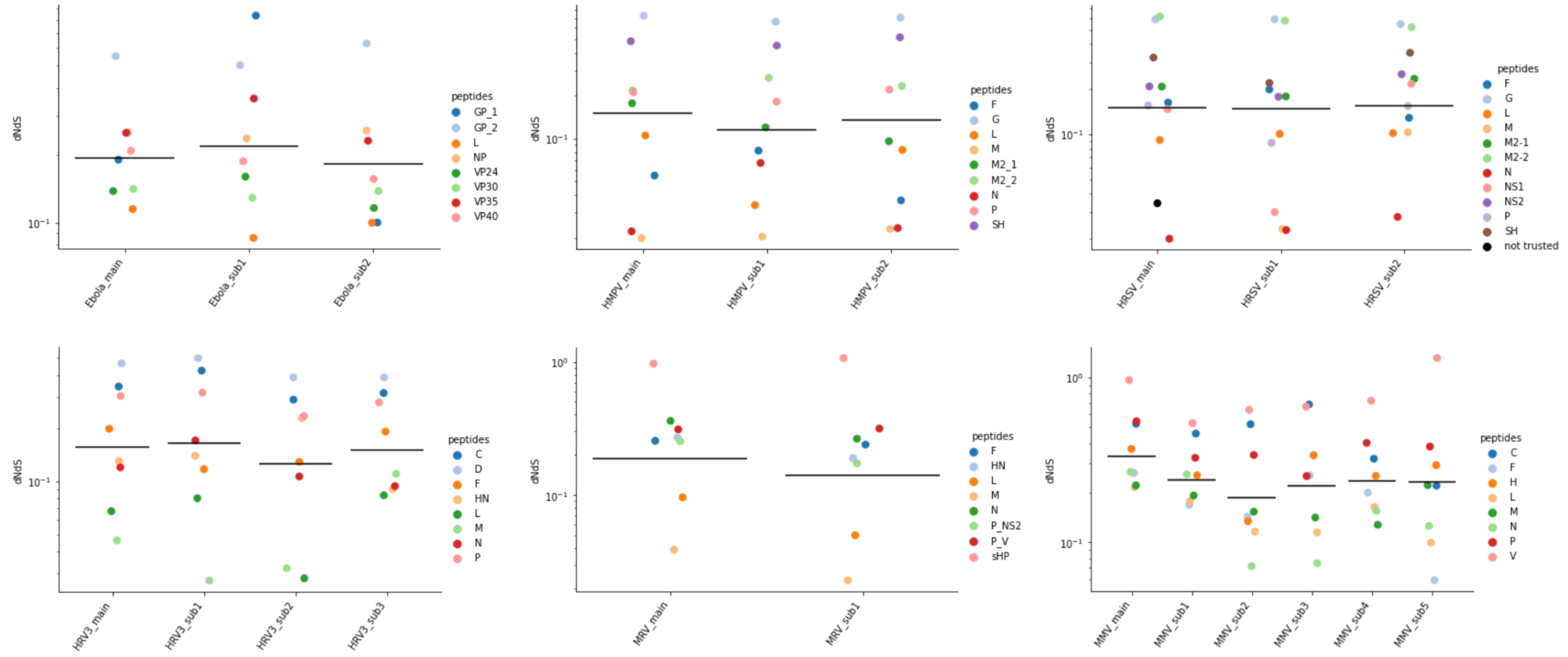
dNdS
$10^{-1}$
Ebola_main
Ebola_sub1
Ebola_sub2
peptides
GP_1
GP_2
L
NP
VP24
VP30
VP35
VP40
HMPV_main
HMPV_sub1
HMPV_sub2
peptides
F
G
L
M
M2_1
M2_2
N
P
SH
HRSV_main
HRSV_sub1
HRSV_sub2
peptides
F
G
L
M
M2-1
M2-2
N
NS1
NS2
P
SH
not trusted
HRV3_main
HRV3_sub1
HRV3_sub2
HRV3_sub3
peptides
C
D
F
HN
L
M
N
P
$10^{0}$
MRV_main
MRV_sub1
peptides
F
HN
L
M
N
P_NS2
P_V
sHP
MMV_main
MMV_sub1
MMV_sub2
MMV_sub3
MMV_sub4
MMV_sub5
peptides
C
F
H
L
M
N
P
V

# S15

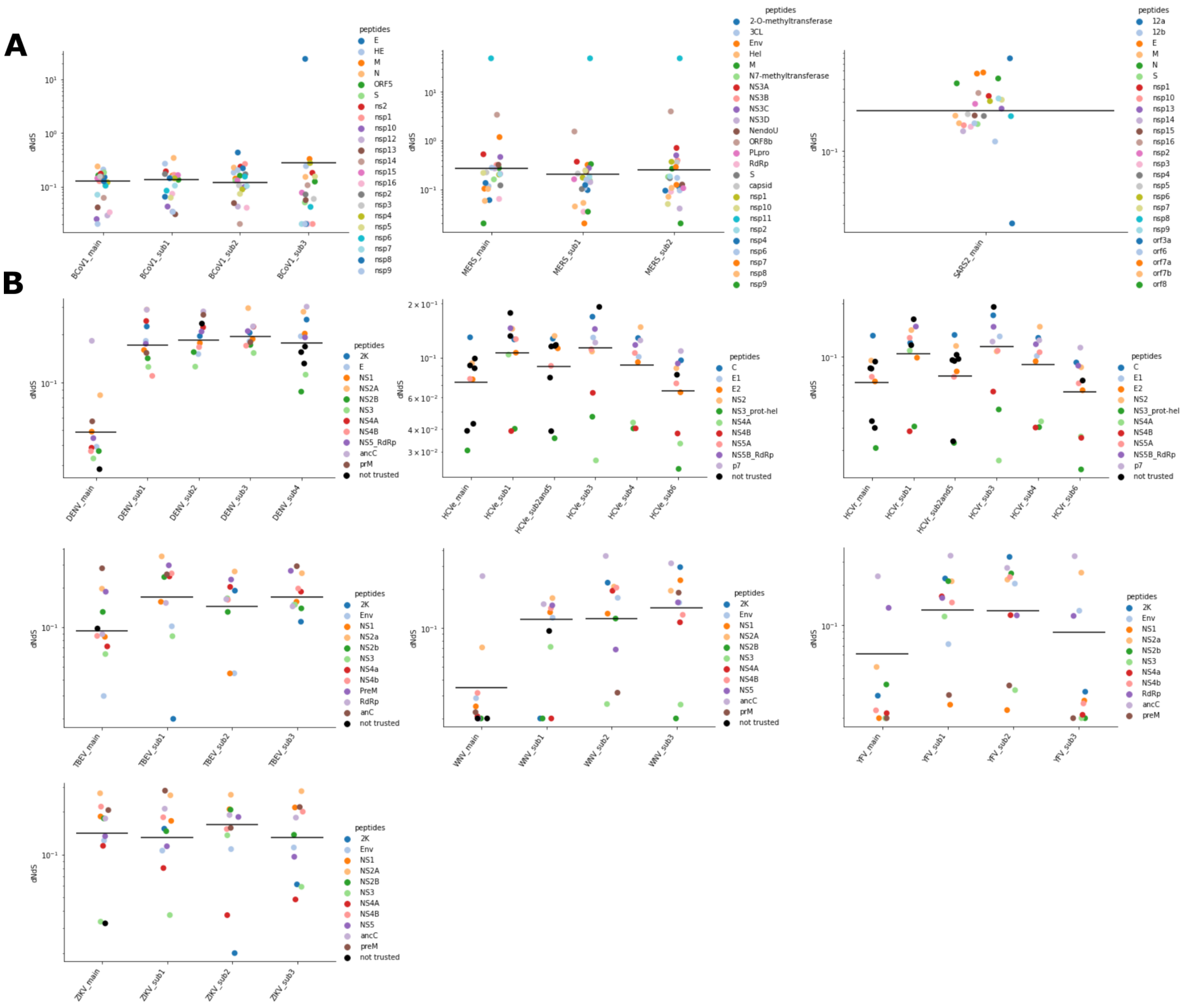
A
B
dNdS
peptides
BCoV1_main
BCoV1_sub1
BCoV1_sub2
BCoV1_sub3
MERS_main
MERS_sub1
MERS_sub2
SARS2_main
DENV_main
DENV_sub1
DENV_sub2
DENV_sub3
DENV_sub4
HCVe_main
HCVe_sub1
HCVe_sub2and5
HCVe_sub3
HCVe_sub4
HCVe_sub6
HCVr_main
HCVr_sub1
HCVr_sub2and5
HCVr_sub3
HCVr_sub4
HCVr_sub6
TBEV_main
TBEV_sub1
TBEV_sub2
TBEV_sub3
WNV_main
WNV_sub1
WNV_sub2
WNV_sub3
YFV_main
YFV_sub1
YFV_sub2
YFV_sub3
ZIKV_main
ZIKV_sub1
ZIKV_sub2
ZIKV_sub3
not trusted

# S16

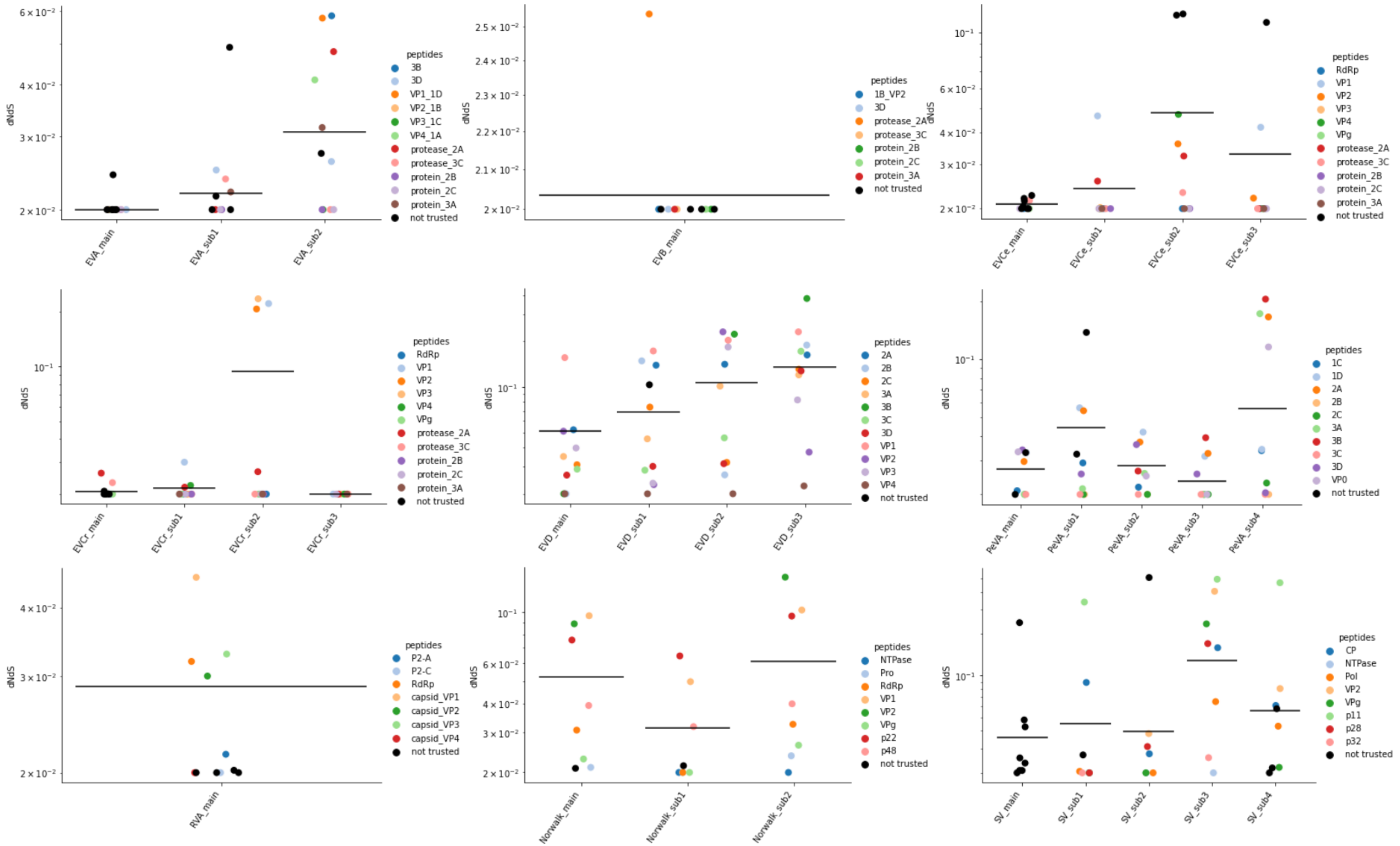
dNdS
EVA_main
EVA_sub1
EVA_sub2
peptides
3B
3D
VP1_1D
VP2_1B
VP3_1C
VP4_1A
protease_2A
protease_3C
protein_2B
protein_2C
protein_3A
not trusted
EVB_main
1B_VP2
3D
protease_2A
protease_3C
protein_2B
protein_2C
protein_3A
not trusted
EVCe_main
EVCe_sub1
EVCe_sub2
EVCe_sub3
RdRp
VP1
VP2
VP3
VP4
VPg
protease_2A
protease_3C
protein_2B
protein_2C
protein_3A
not trusted
EVCr_main
EVCr_sub1
EVCr_sub2
EVCr_sub3
EVD_main
EVD_sub1
EVD_sub2
EVD_sub3
2A
2B
2C
3A
3B
3C
3D
VP1
VP2
VP3
VP4
not trusted
PeVA_main
PeVA_sub1
PeVA_sub2
PeVA_sub3
PeVA_sub4
1C
1D
2A
2B
2C
3A
3B
3C
3D
VP0
not trusted
RVA_main
P2-A
P2-C
RdRp
capsid_VP1
capsid_VP2
capsid_VP3
capsid_VP4
not trusted
Norwalk_main
Norwalk_sub1
Norwalk_sub2
NTPase
Pro
RdRp
VP1
VP2
VPg
p22
p48
not trusted
SV_main
SV_sub1
SV_sub2
SV_sub3
SV_sub4
CP
NTPase
Pol
VP2
VPg
p11
p28
p32
not trusted

# S17

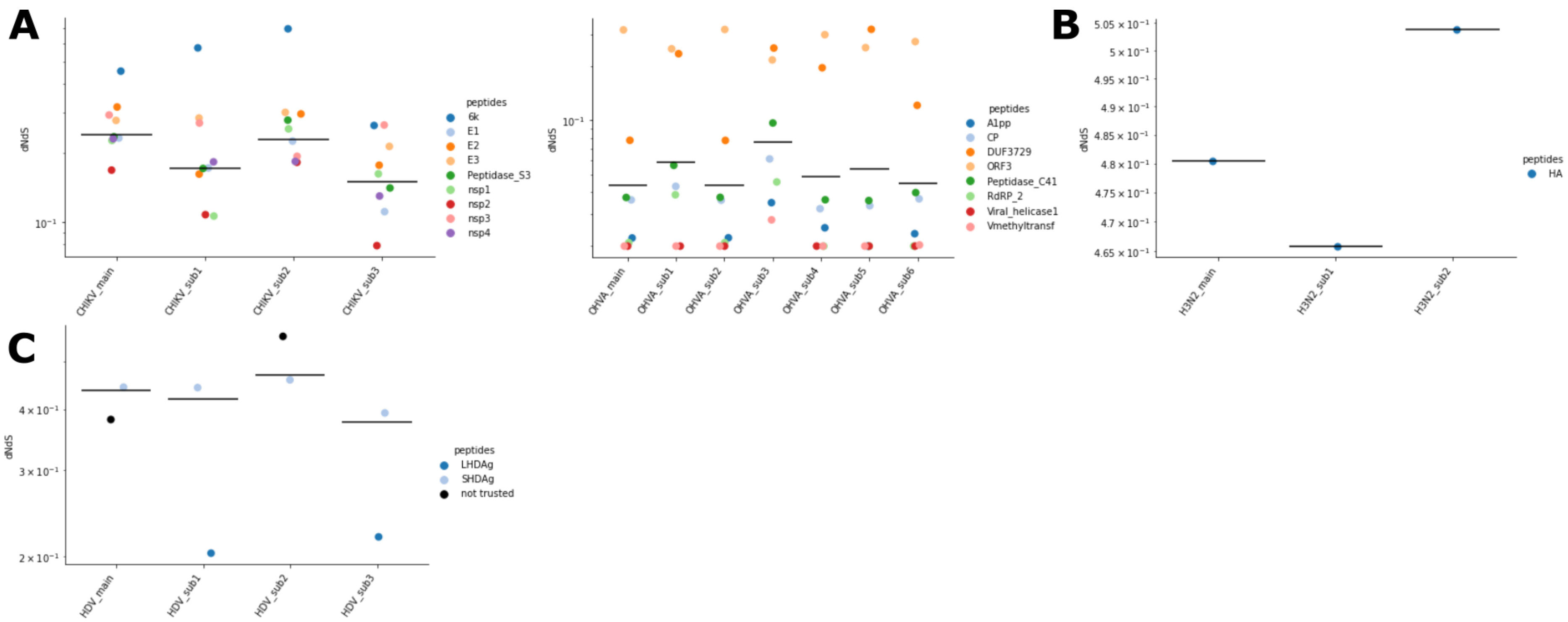

A
B
C
dNdS
peptides
6k
E1
E2
E3
Peptidase_S3
nsp1
nsp2
nsp3
nsp4
CHIKV_main
CHIKV_sub1
CHIKV_sub2
CHIKV_sub3
A1pp
CP
DUF3729
ORF3
Peptidase_C41
RdRP_2
Viral_helicase1
Vmethyltransf
OHVA_main
OHVA_sub1
OHVA_sub2
OHVA_sub3
OHVA_sub4
OHVA_sub5
OHVA_sub6
HA
H3N2_main
H3N2_sub1
H3N2_sub2
LHDAg
SHDAg
not trusted
HDV_main
HDV_sub1
HDV_sub2
HDV_sub3

# S18

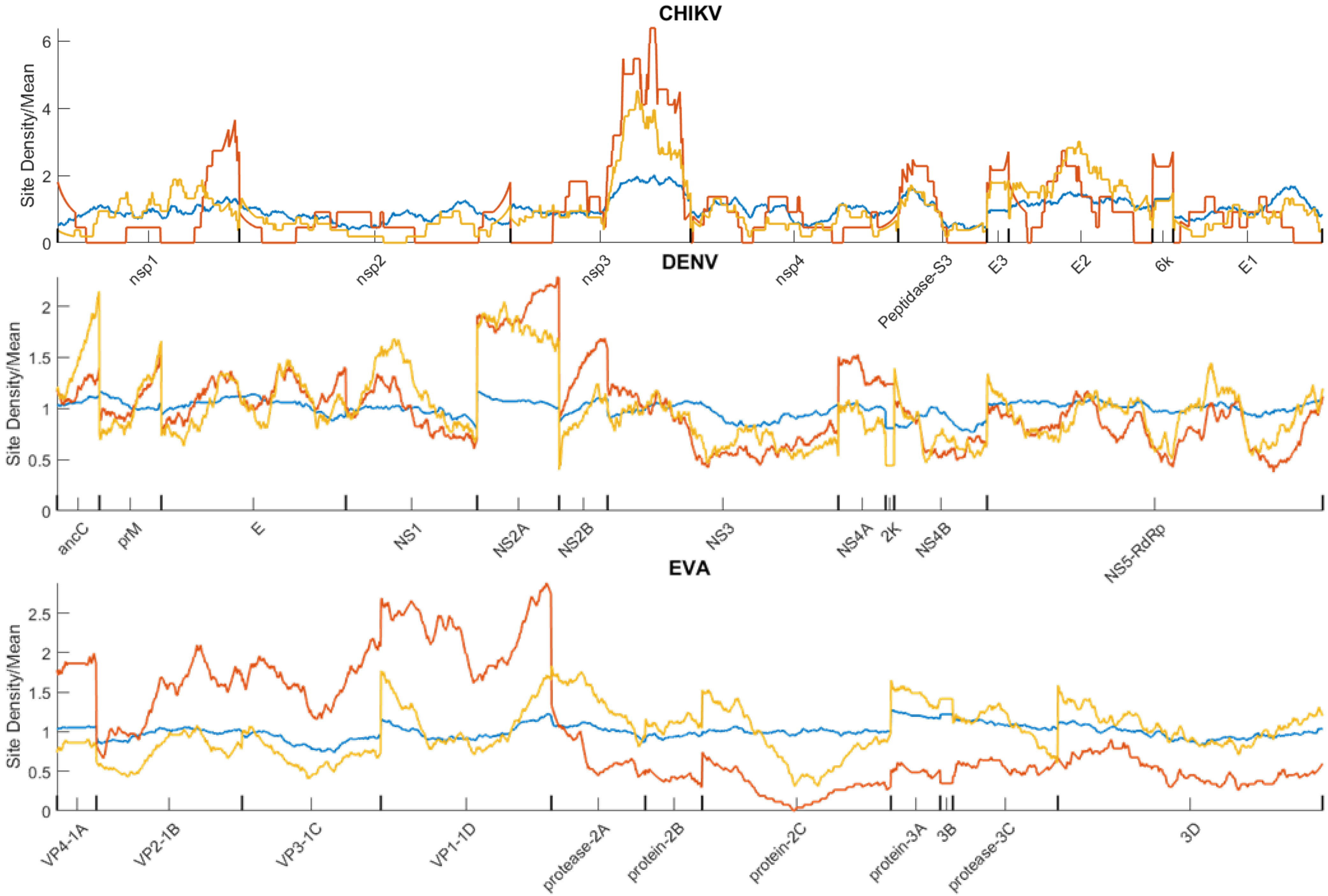
CHIKV
Site Density/Mean
0
2
4
6
nsp1
nsp2
nsp3
nsp4
Peptidase-S3
E3
E2
6k
E1
DENV
Site Density/Mean
0
0.5
1
1.5
2
ancC
prM
E
NS1
NS2A
NS2B
NS3
NS4A
2K
NS4B
NS5-RdRp
EVA
Site Density/Mean
0
0.5
1
1.5
2
2.5
VP4-1A
VP2-1B
VP3-1C
VP1-1D
protease-2A
protein-2B
protein-2C
protein-3A
3B
protease-3C
3D

**S19**

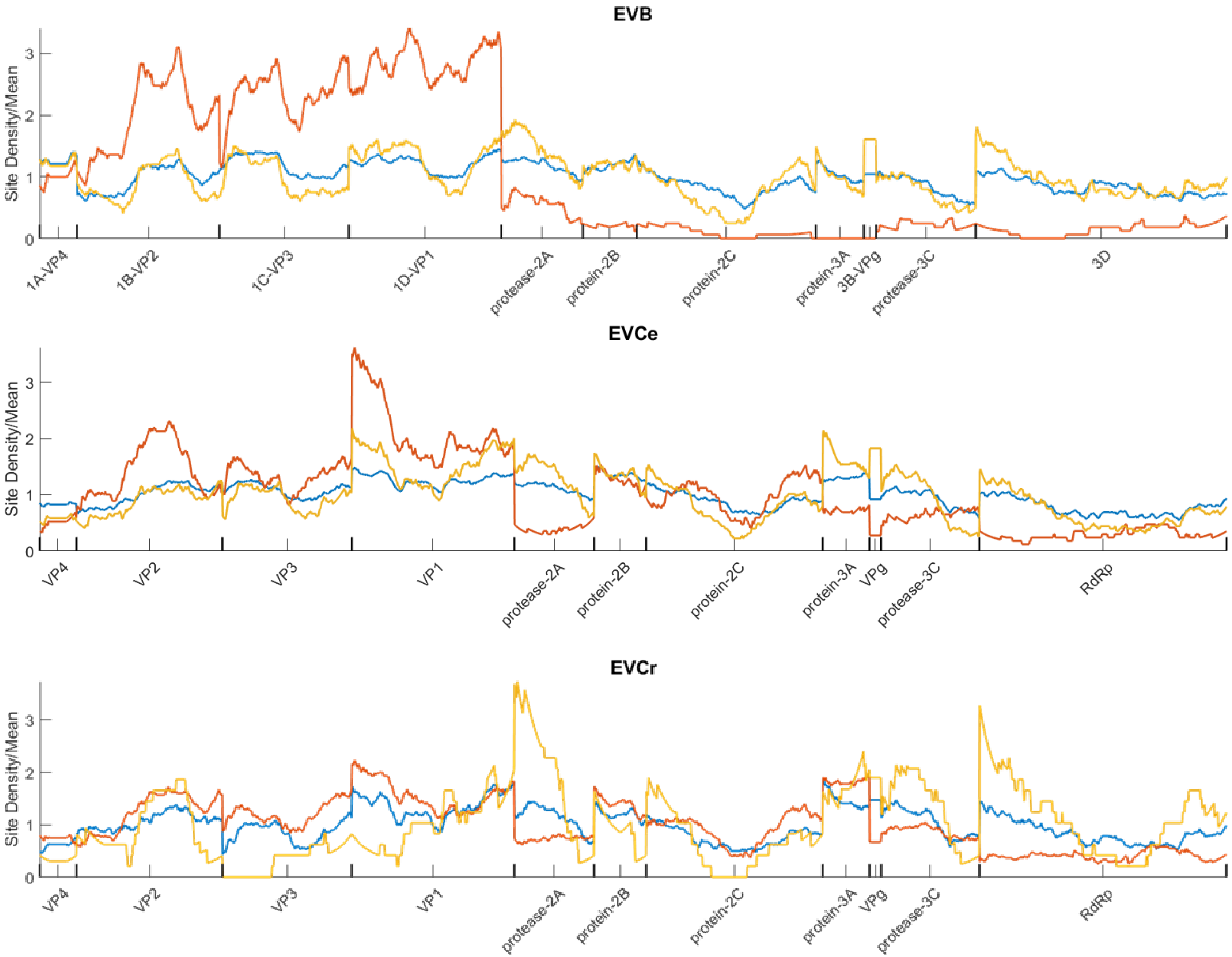

EVB
Site Density/Mean
0
1
2
3
1A-VP4
1B-VP2
1C-VP3
1D-VP1
protease-2A
protein-2B
protein-2C
protein-3A
3B-VPg
protease-3C
3D
EVCe
Site Density/Mean
0
1
2
3
VP4
VP2
VP3
VP1
protease-2A
protein-2B
protein-2C
protein-3A
VPg
protease-3C
RdRp
EVCr
Site Density/Mean
0
1
2
3
VP4
VP2
VP3
VP1
protease-2A
protein-2B
protein-2C
protein-3A
VPg
protease-3C
RdRp

S20

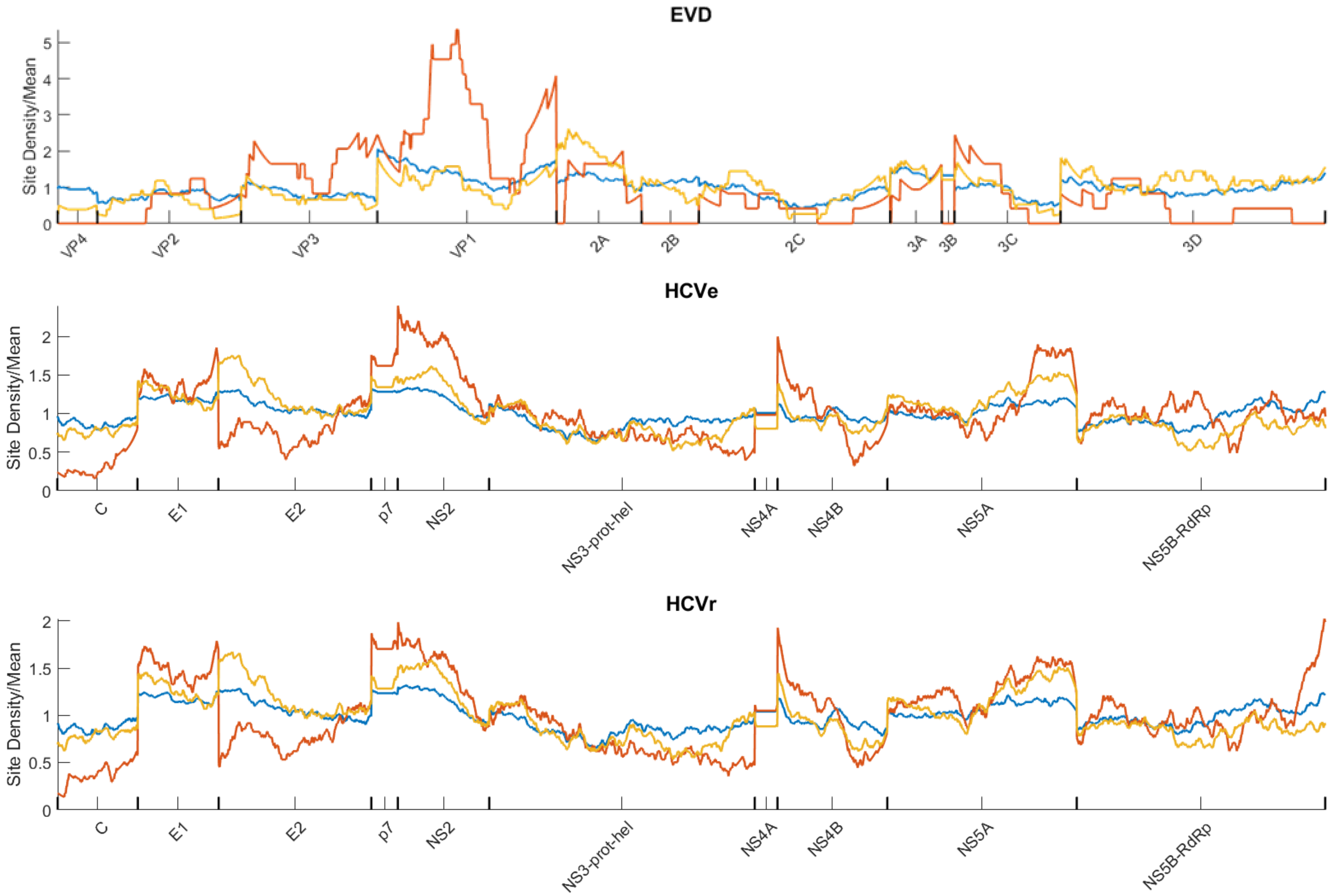
EVD
Site Density/Mean
5
4
3
2
1
0
VP4
VP2
VP3
VP1
2A
2B
2C
3A
3B
3C
3D
HCVe
Site Density/Mean
2
1.5
1
0.5
0
C
E1
E2
p7
NS2
NS3-prot-hel
NS4A
NS4B
NS5A
NS5B-RdRp
HCVr
Site Density/Mean
2
1.5
1
0.5
0
C
E1
E2
p7
NS2
NS3-prot-hel
NS4A
NS4B
NS5A
NS5B-RdRp

**S21**

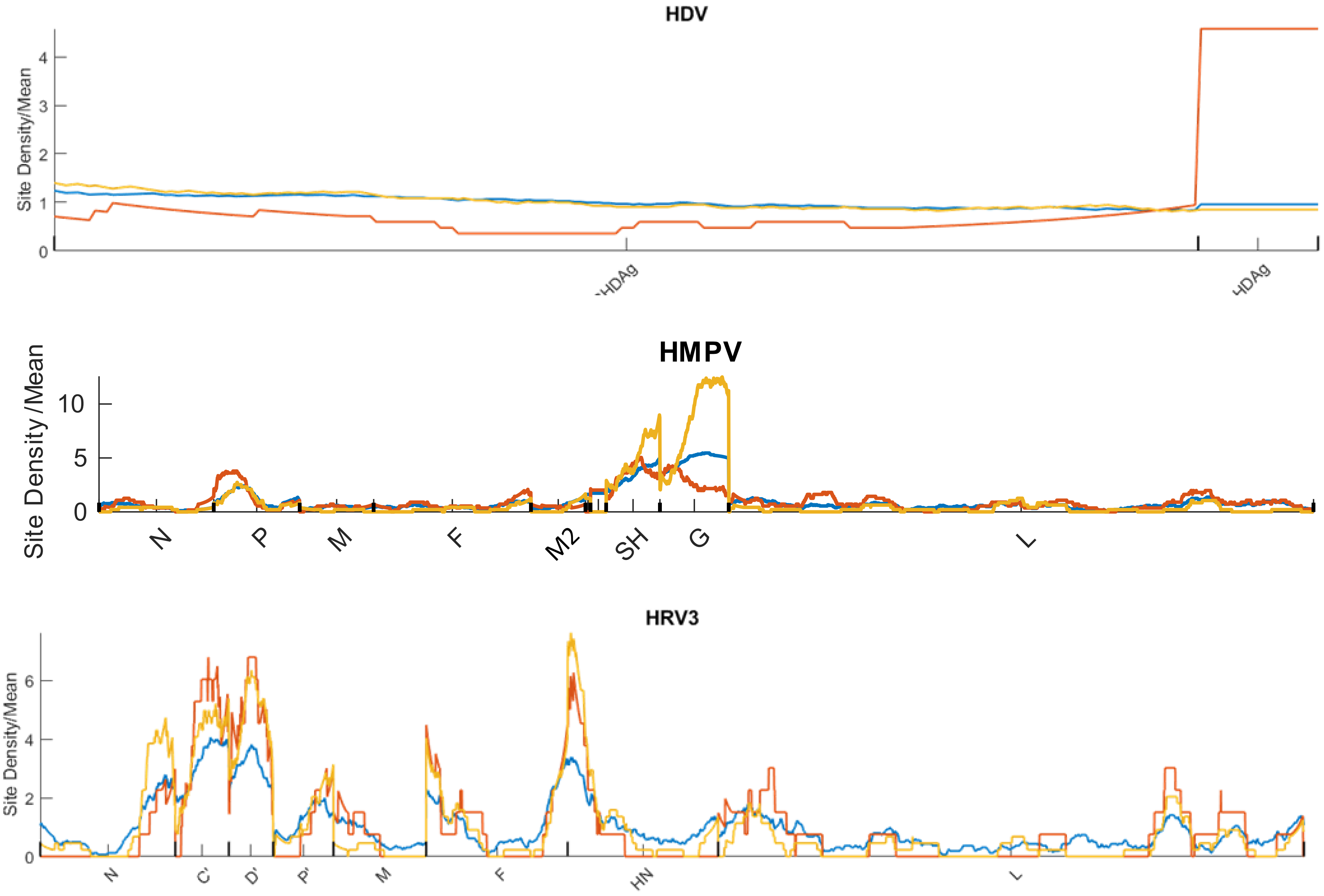
HDV
Site Density/Mean
0
1
2
3
4
HDAg
HDAg
HMPV
Site Density/Mean
0
5
10
N
P
M
F
M2
SH
G
L
HRV3
Site Density/Mean
0
2
4
6
N
C'
D'
P'
M
F
HN
L

**S22**

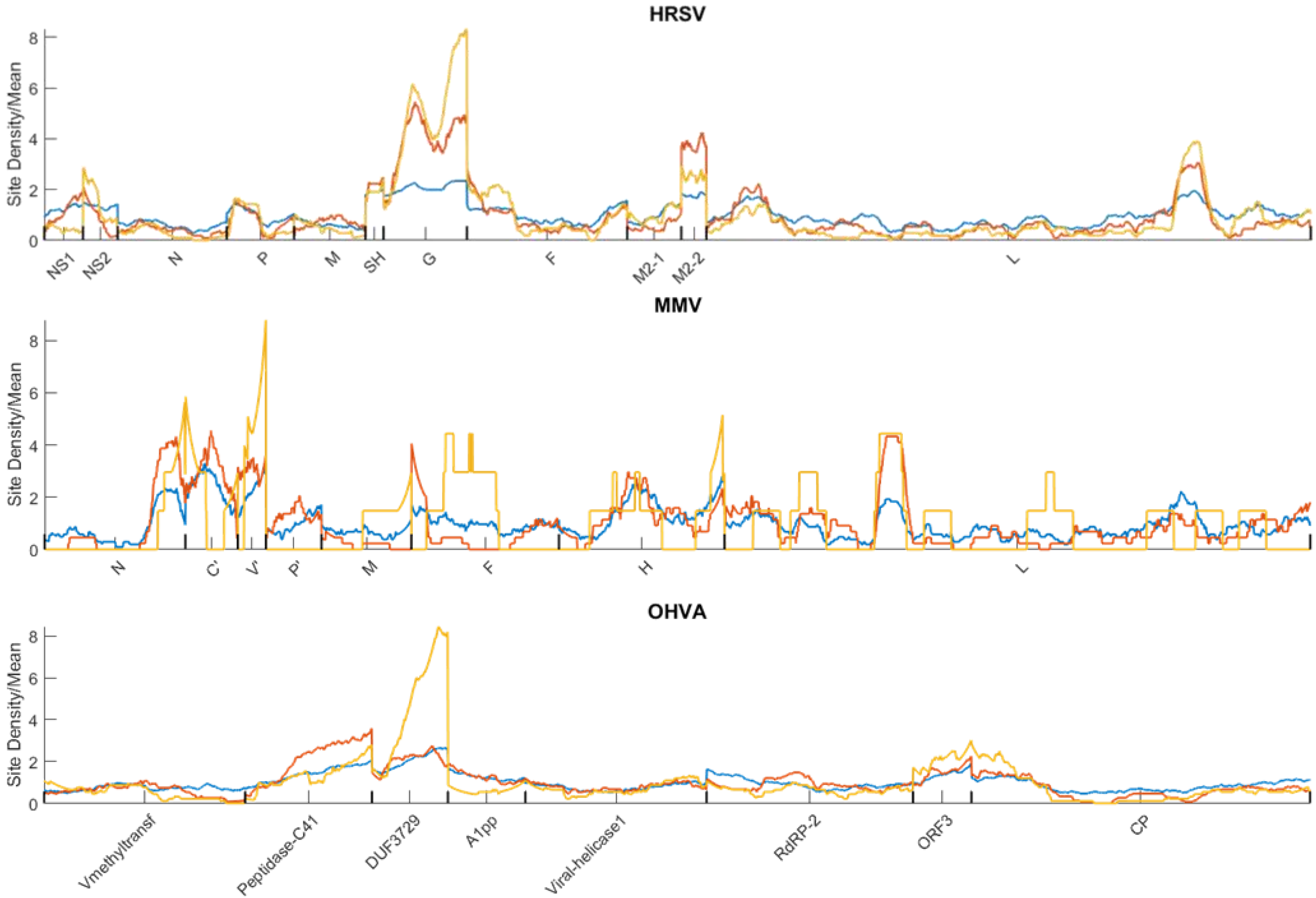
HRSV
Site Density/Mean
8
6
4
2
0
NS1
NS2
N
P
M
SH
G
F
M2-1
M2-2
L
MMV
Site Density/Mean
8
6
4
2
0
N
C'
V'
P'
M
F
H
L
OHVA
Site Density/Mean
8
6
4
2
0
Vmethyltransf
Peptidase-C41
DUF3729
A1pp
Viral-helicase1
RdRP-2
ORF3
CP

S23

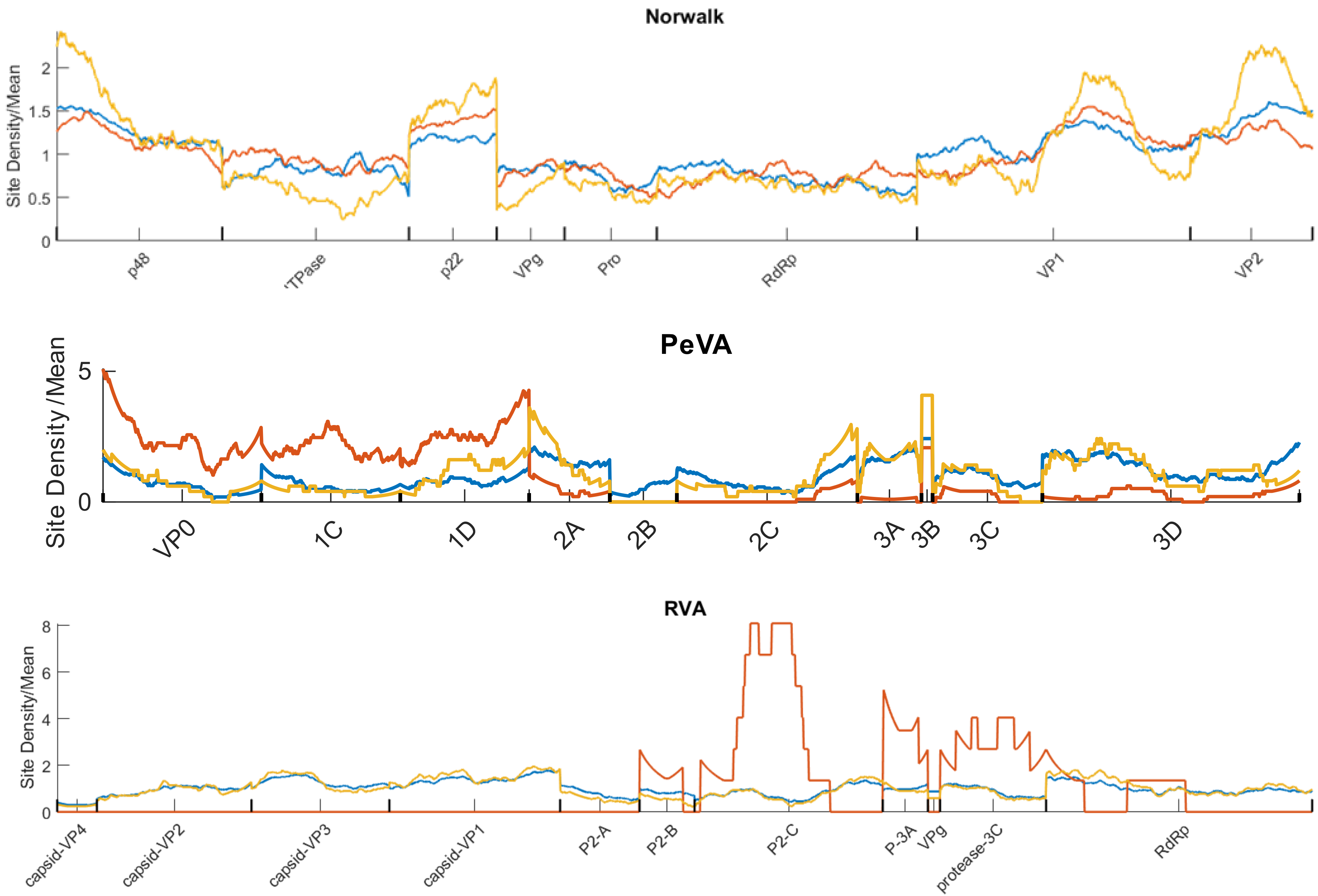

Norwalk
Site Density/Mean
0
0.5
1
1.5
2
p48
TPase
p22
VPg
Pro
RdRp
VP1
VP2
PeVA
Site Density /Mean
0
5
VP0
1C
1D
2A
2B
2C
3A
3B
3C
3D
RVA
Site Density/Mean
0
2
4
6
8
capsid-VP4
capsid-VP2
capsid-VP3
capsid-VP1
P2-A
P2-B
P2-C
P-3A
VPg
protease-3C
RdRp

**S24**

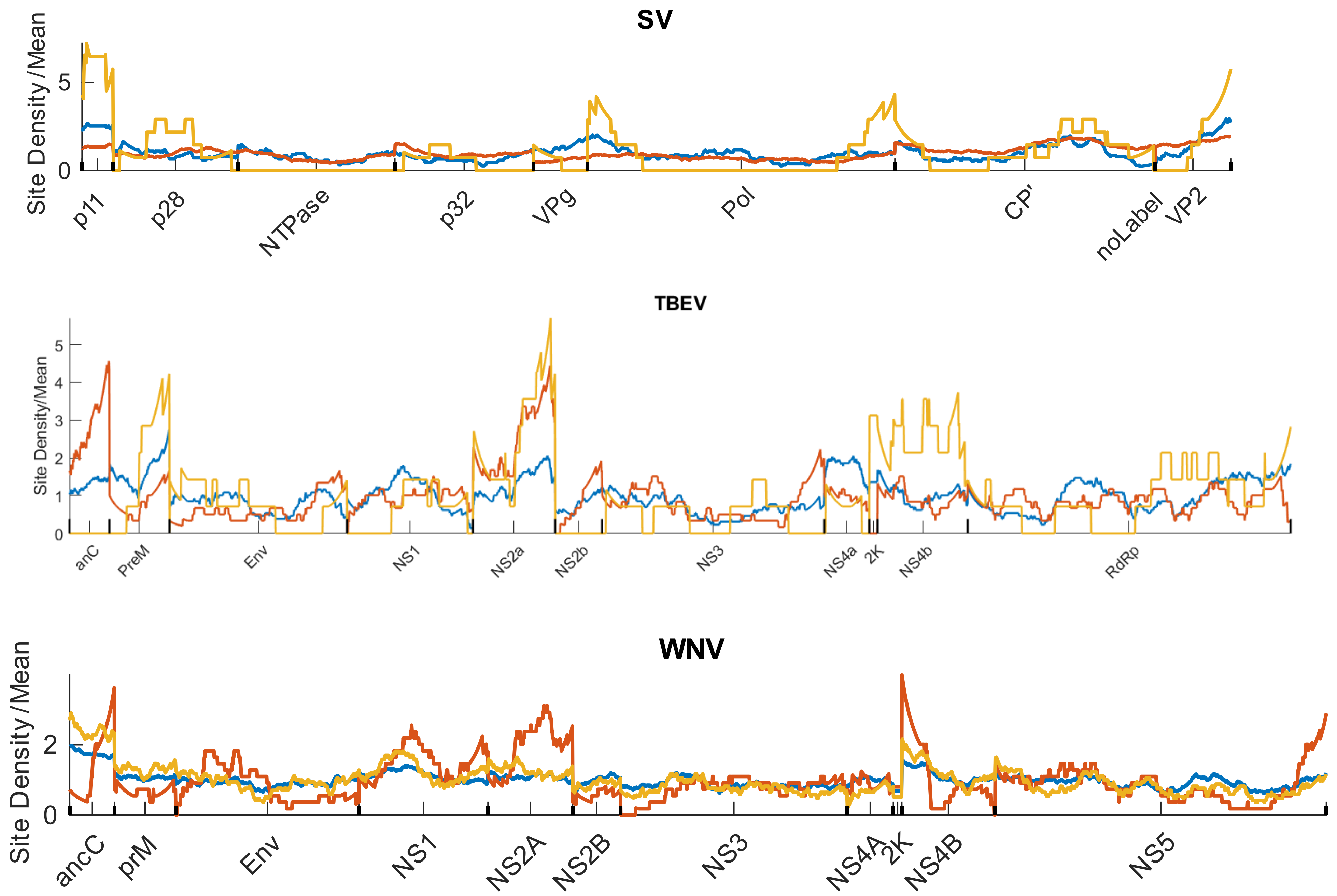
SV
Site Density /Mean
5
0
p11
p28
NTPase
p32
VPg
Pol
CP'
noLabel
VP2
TBEV
Site Density/Mean
5
4
3
2
1
0
anC
PreM
Env
NS1
NS2a
NS2b
NS3
NS4a
2K
NS4b
RdRp
WNV
Site Density /Mean
2
0
ancC
prM
Env
NS1
NS2A
NS2B
NS3
NS4A
2K
NS4B
NS5

# S25

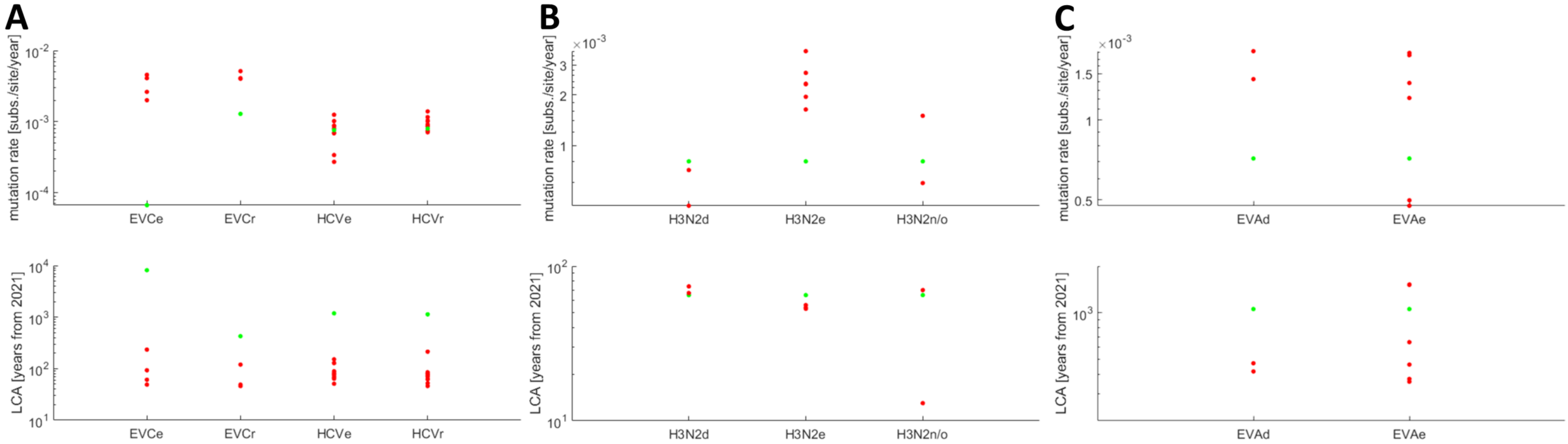

A
B
C
mutation rate [subs./site/year]
10^-2
10^-3
10^-4
EVCe
EVCr
HCVe
HCVr
×10^-3
3
2
1
H3N2d
H3N2e
H3N2n/o
×10^-3
1.5
1
0.5
EVAd
EVAe
LCA [years from 2021]
10^4
10^3
10^2
10^1
EVCe
EVCr
HCVe
HCVr
10^2
10^1
H3N2d
H3N2e
H3N2n/o
10^3
EVAd
EVAe

# S26

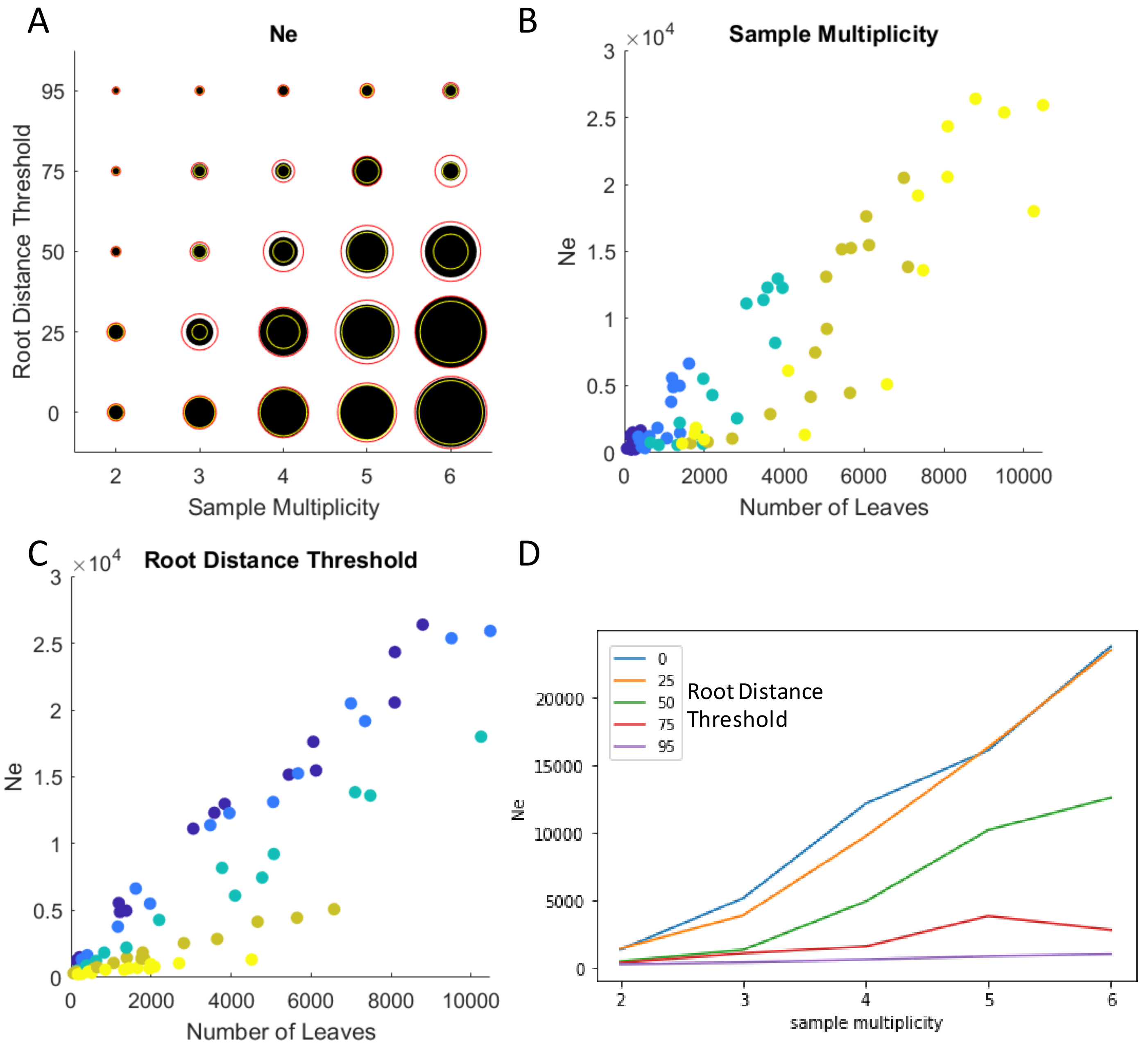
A
Ne
Root Distance Threshold
95
75
50
25
0
2
3
4
5
6
Sample Multiplicity
B
Sample Multiplicity
×10^4
Ne
Number of Leaves
C
Root Distance Threshold
×10^4
Ne
Number of Leaves
D
0
25
50
75
95
Root Distance Threshold
Ne
sample multiplicity

# S27

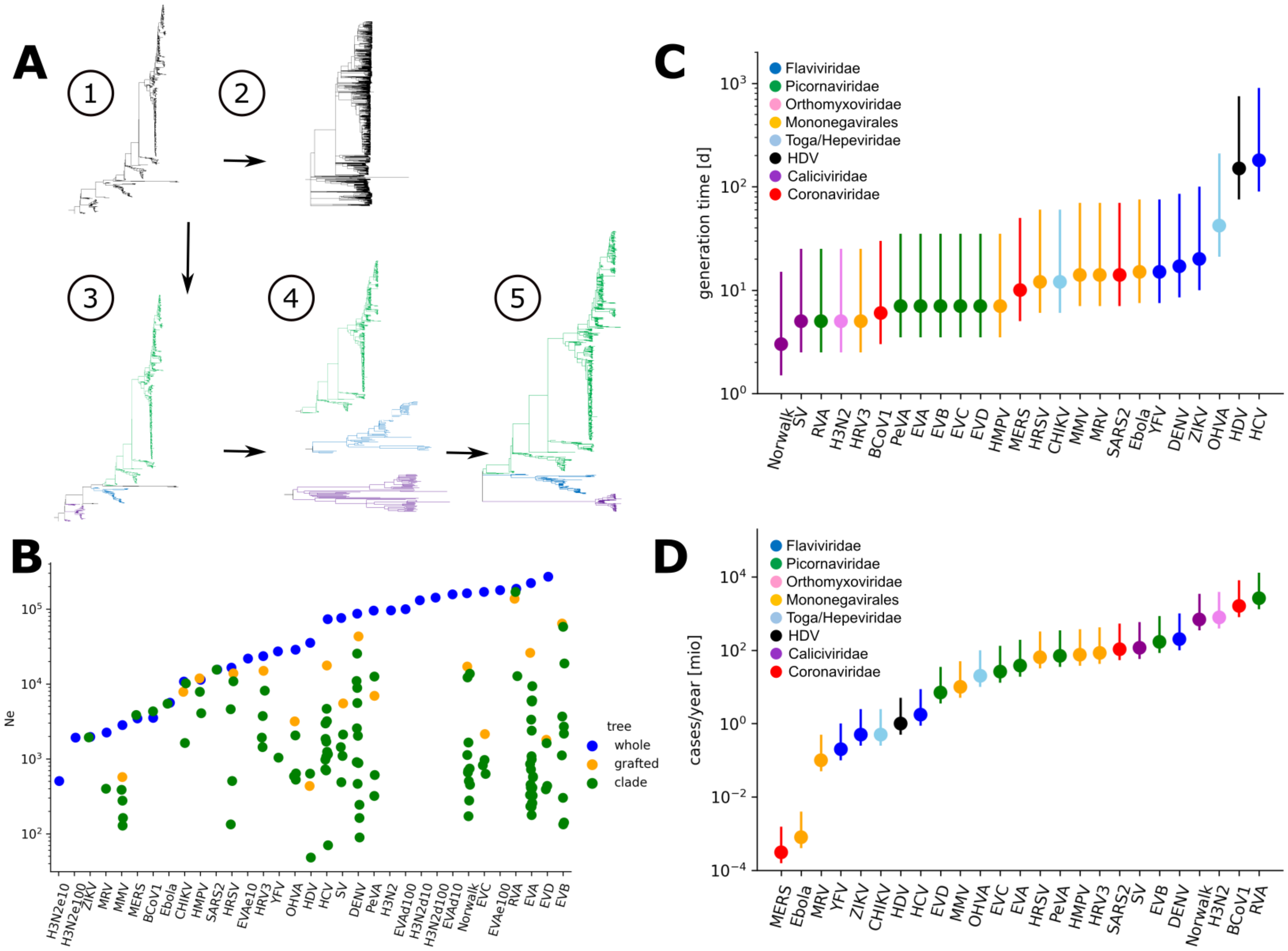
A
1
2
3
4
5
B
Ne
tree
whole
grafted
clade
C
generation time [d]
Flaviviridae
Picornaviridae
Orthomyxoviridae
Mononegavirales
Toga/Hepeviridae
HDV
Caliciviridae
Coronaviridae
D
cases/year [mio]

# S28

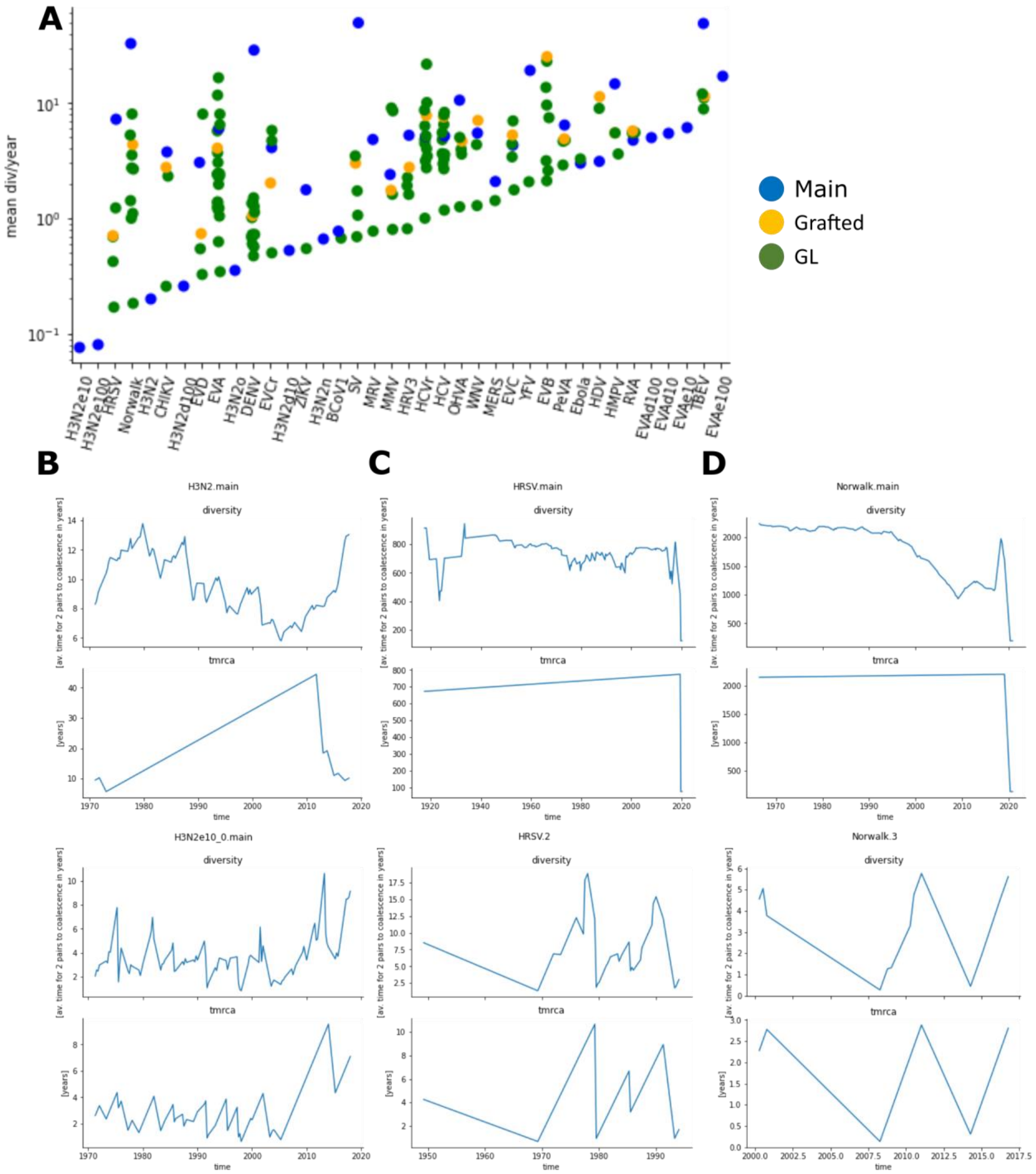
A
mean div/year
Main
Grafted
GL
H3N2e10
H3N2e100
HRSV
Norwalk
H3N2
CHIKV
H3N2d100
EVD
EVA
H3N2o
DENV
EVCr
H3N2d10
ZIKV
H3N2n
BCoV1
SV
MRV
MMV
HRV3
HCVr
HCV
OHVA
WNV
MERS
EVC
YFV
EVB
PeVA
Ebola
HDV
HMPV
RVA
EVAd100
EVAd10
EVAe10
TBEV
EVAe100
B
H3N2.main
diversity
tmrca
time
H3N2e10_0.main
C
HRSV.main
HRSV.2
D
Norwalk.main
Norwalk.3
[av. time for 2 pairs to coalescence in years]
[years]

# S29

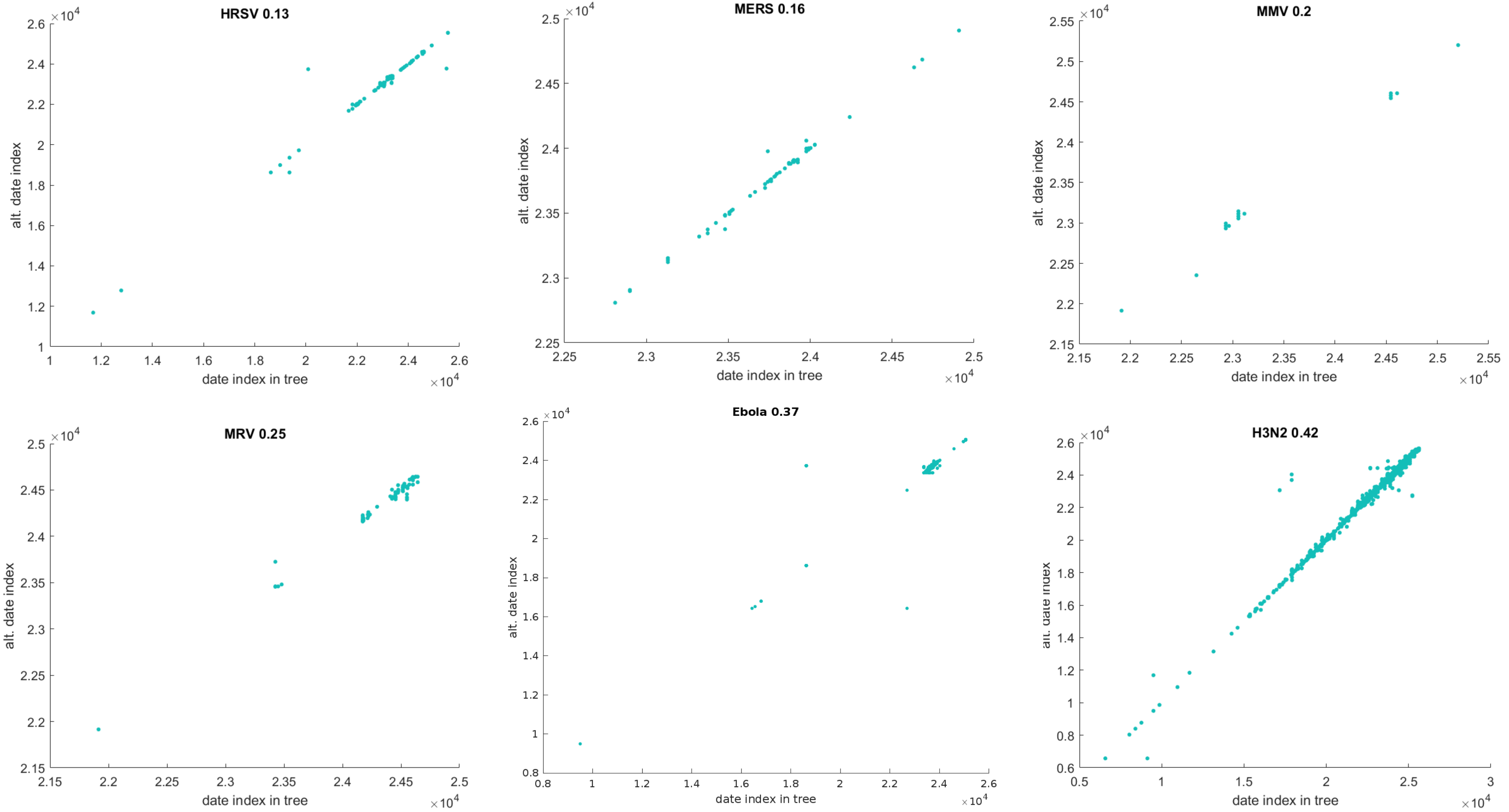
HRSV 0.13
MERS 0.16
MMV 0.2
MRV 0.25
Ebola 0.37
H3N2 0.42
alt. date index
date index in tree
×10^4
